# Robust integrated intracellular organization of the human iPS cell: where, how much, and how variable

**DOI:** 10.1101/2020.12.08.415562

**Authors:** Matheus P. Viana, Jianxu Chen, Theo A. Knijnenburg, Ritvik Vasan, Calysta Yan, Joy E. Arakaki, Matte Bailey, Ben Berry, Antoine Borensztejn, Jackson M. Brown, Sara Carlson, Julie A. Cass, Basudev Chaudhuri, Kimberly R. Cordes Metzler, Mackenzie E. Coston, Zach J. Crabtree, Steve Davidson, Colette M. DeLizo, Shailja Dhaka, Stephanie Q. Dinh, Thao P. Do, Justin Domingus, Rory M. Donovan-Maiye, Tyler J. Foster, Christopher L. Frick, Griffin Fujioka, Margaret A. Fuqua, Jamie L. Gehring, Kaytlyn A. Gerbin, Tanya Grancharova, Benjamin W. Gregor, Lisa J. Harrylock, Amanda Haupt, Melissa C. Hendershott, Caroline Hookway, Alan R. Horwitz, Chris Hughes, Eric J. Isaac, Gregory R. Johnson, Brian Kim, Andrew N. Leonard, Winnie W. Leung, Jordan J. Lucas, Susan A. Ludmann, Blair M. Lyons, Haseeb Malik, Ryan McGregor, Gabe E. Medrash, Sean L. Meharry, Kevin Mitcham, Irina A. Mueller, Timothy L. Murphy-Stevens, Aditya Nath, Angelique M. Nelson, Luana Paleologu, T. Alexander Popiel, Megan M. Riel-Mehan, Brock Roberts, Lisa M. Schaefbauer, Magdalena Schwarzl, Jamie Sherman, Sylvain Slaton, M. Filip Sluzewski, Jacqueline E. Smith, Youngmee Sul, Madison J. Swain-Bowden, W. Joyce Tang, Derek J. Thirstrup, Daniel M. Toloudis, Andrew P. Tucker, Veronica Valencia, Winfried Wiegraebe, Thushara Wijeratna, Ruian Yang, Rebecca J. Zaunbrecher, Allen Institute for Cell Science, Graham T. Johnson, Ruwanthi N. Gunawardane, Nathalie Gaudreault, Julie A. Theriot, Susanne M. Rafelski

## Abstract

Despite the intimate link between cell organization and function, the principles underlying intracellular organization and the relation between organization, gene expression and phenotype are not well understood. We address this by creating a benchmark for mean cell organization and the natural range of cell-to-cell variation. This benchmark can be used for comparison to other normal or abnormal cell states. To do this, we developed a reproducible microscope imaging pipeline to generate a high-quality dataset of 3D, high-resolution images of over 200,000 live cells from 25 isogenic human induced pluripotent stem cell (hiPSC) lines from the Allen Cell Collection. Each line contains one fluorescently tagged protein, created via endogenous CRISPR/Cas9 gene editing, representing a key cellular structure or organelle. We used these images to develop a new multi-part and generalizable analysis approach of the locations, amounts, and variation of these 25 cellular structures. Taking an integrated approach, we found that both the extent to which a structure’s individual location varied (“stereotypy”) and the extent to which the structure localized relative to all the other cellular structures (“concordance”) were robust to a wide range of cell shape variation, from flatter to taller, smaller to larger, or less to more polarized cells. We also found that these cellular structures varied greatly in how their volumes scaled with cell and nuclear size. These analyses create a data-driven set of quantitative rules for the locations, amounts, and variation of 25 cellular structures within the hiPSC as a normal baseline for cell organization.

## Introduction

A living cell must organize all of its millions of subcellular components and processes in space and time through as many as four orders of magnitude. At the nanometer scale, specific molecular interactions permit the assembly of macromolecules and organelles to perform and regulate cell function. More global cell behaviors, however, can occur over scales of tens of microns, such as the coordinated protrusion of a cell front and retraction of a rear during cell migration (Lauffenburger and Horwitz, 1996). Identifying the rules of cell organization and understanding how they facilitate global behaviors across this broad span of spatial scales is an immensely complex and daunting task that must also incorporate dynamic changes across a broad temporal spectrum. However, to understand cell organization at the level of the major intracellular machinery and organelles (cellular structures), requires the study of only ∼25-50 of these structures. This enormously reduces the dimensional complexity, making feasible the quest for an interpretable and testable set of rules that govern cell organization and how this organization changes as cells transition to alternative normal or abnormal cell behaviors. For example, measuring the locations of each of these cellular structures relative to all the others, as well as the total volume occupied by each structure, creates a rich set of quantitative rule-building constraints for generating and testing models of cell organization (Johnson et al., 2015; Macklin et al., 2020).

A significant potential challenge, even for this approach, is that cells must behave robustly yet respond sensitively to their ever-changing environments. As a result, a population of normal, putatively identical cells might exhibit significant cell-to-cell variability. Thus, it is important to establish a baseline with which different kinds of cells can be compared. This baseline should represent the typical, or mean, cell within the population, as well as the full range of normal variation of the population itself. An abnormal cell phenotype may exhibit not only a shift in the mean but also a shift in the variation (Roggiani and Goulian, 2015). Therefore, a meaningful and useful description of normal cell organization requires quantitative measurements, not just of the locations or amounts of each of the cellular structures, but also how they vary within a large group of normal cells.

To establish a normal baseline for cell organization, we turned to human induced pluripotent stem cells (hiPSCs), which represent an early embryonic cell state and an ideal human model system. hiPSCs are naturally immortal, karyotypically normal, and can be induced to differentiate into other cell types (Drubin and Hyman, 2017). We previously developed methods to generate a series of isogenic clonal hiPSC lines expressing fluorescent protein tags for visualizing specific organelles and cellular structures via endogenous CRISPR/Cas9 gene editing. We performed extensive quality control on these lines to create the *Allen Cell Collection* (Roberts et al., 2017a). In this work, we imaged 25 lines from this collection, each containing one fluorescently tagged protein representing a key cellular structure or organelle.

Here we present the *hiPSC Single-Cell Image Dataset*, an unprecedented collection of high-resolution, 3D images of over 200,000 live cells. To analyze this large-scale dataset, we develop generalizable, quantitative methods that permit direct comparisons of the similarity of overall cell and nuclear shapes for 3D cell image data, to build a simple and human-interpretable “shape space”. This approach facilitates the robust identification of clusters of cells that are most similar to each other in their overall shape. We also introduce a generalizable method to parameterize fluorescence intensity distributions in 3D cell images; this method allows the actual distribution observed for a particular cell to be robustly “morphed” into another similar cell shape, without losing substantial quantitative information about fluorescence localization. We then apply these methods to determine which structures are most highly stereotyped with respect to their cell-to-cell variation, and also which pairs of structures are most similar to each other, throughout the normal hiPSC shape space. These analyses create a data-driven set of quantitative rule-building constraints for the locations, amounts, and variation of 25 cellular structures within the hiPSC as a normal baseline for cell organization and a fundamental benchmark for comparison with future analyses of cell shape and cell organization for cells in different states.

## Results

### An hiPSC Single-Cell Image Dataset contains over 200,000 live, high-resolution, 3D cells spanning 25 cellular structures

We previously developed methods and quality control workflows to create the Allen Cell Collection (Roberts et al., 2017a) and www.allencell.org) of hiPSC lines, each expressing a single endogenously tagged protein representing a particular organelle or cellular structure. Here we created 15 additional cell lines and used a total set of 25 cell lines permitting a holistic view of cells at the level of their the major organelles, cellular structures, and compartments (Table 1).

**Table 1:** Fluorescently tagged cellular structures used to create the hiPSC Single-Cell Image Dataset.

| Structure | Protein | Gene | Location | AICS line |
| --- | --- | --- | --- | --- |
| nucleoli (DFC) | fibrillarin | *FBL* | nuclear | **AICS-0014 cl. 6** |
| nucleoli (GC) | nucleophosmin | *NPM1* | nuclear | AICS-0057 cl. 50 |
| nuclear speckles | SON | *SON* | nuclear | AICS-0094 cl. 24 |
| cohesins | SMC-1A | *SMC1A* | nuclear | AICS-0068 cl. 9 |
| histones | H2B | *H2BC11* | nuclear | AICS-0061 cl. 36 |
| nuclear envelope | lamin B1 | *LMNB1* | nuclear periphery | **AICS-0013 cl. 210** |
| nuclear pores | Nup153 | *NUP153* | nuclear periphery | AICS-0069 cl. 88 |
| ER (Sec61 beta) | Sec61 beta | *SEC61B* | cytoplasm | **AICS-0010 cl. 55** |
| ER (SERCA2) | SERCA2 | *ATP2A2* | cytoplasm | AICS-0046 cl. 51 |
| mitochondria | Tom20 | *TOMM20* | cytoplasm | **AICS-0011 cl. 27** |
| peroxisomes | PMP34 | *SLC25A17* | cytoplasm | AICS-0033 cl. 115 |
| endosomes | Rab-5A | *RAB5A* | cytoplasm | AICS-0040 cl. 35* |
| lysosomes | LAMP-1 | *LAMP1* | cytoplasm | AICS-0022 cl. 37 |
| Golgi | sialyltransferase 1 | *ST6GAL1* | cytoplasm | AICS-0025 cl. 44* |
| centrioles | centrin-2 | *CETN2* | cytoplasm | AICS-0032 cl. 19*** |
| microtubules | alpha-tubulin | *TUBA1B* | cytoplasm | **AICS-0012 cl. 105** |
| plasma membrane | CAAX | *AAVS1* | cell periphery | AICS-0054 cl. 91*** |
| actin filaments | beta-actin | *ACTB* | cell periphery | **AICS-0016 cl. 184** |
| actin bundles | alpha-actinin-1 | *ACTN1* | cell periphery | AICS-0007 cl. 79** |
| actomyosin bundles | non-muscle myosin IIB | *MYH10* | cell periphery | **AICS-0024 cl. 80** |
| gap junctions | connexin-43 | *GJA1* | cell periphery | AICS-0053 cl. 16 |
| tight junctions | ZO-1 | *TJP1* | cell periphery | **AICS-0023 cl. 20** |
| desmosomes | desmoplakin | *DSP* | cell periphery | **AICS-0017 cl. 65** |
| adherens junctions | beta-catenin | *CTNNB1* | cell periphery | AICS-0058 cl. 67 |
| matrix adhesions | paxillin | *PXN* | cell periphery | **AICS-0005 cl. 50** |

\* biallelic edit, \*\* line not yet released, \*\*\* mTagRFP-T fluorophore tag instead of mEGFP fluorophore tag Bold AICS line number indicates cell lines published previously (Roberts, et al., 2017).

hiPSCs grow in tightly packed, epithelial-like monolayer colonies (Roberts et al., 2017a), requiring well-defined imaging assay guidelines for reproducible data collection. We grow these cells on Matrigel-coated glass plates compatible with high-resolution imaging while preserving their normal pluripotent state (Roberts et al., 2017a). We built an, automated microscopy imaging pipeline to Table 1: Fluorescently tagged cellular structures used to create the hiPSC Single-Cell Image Dataset reproducibly generate the living colonies, imaged the cells in 3D using spinning-disk confocal microscopes to collect standardized 3D information, and processed these images to create the hiPSC Single-Cell Image Dataset (Figure 1). We mostly imaged cells halfway towards the centers of large, well-packed colonies, as cells behaved most consistently in this region. We also captured variations in colony area and locations within the colony, and enriched for images with mitotic cells when necessary (Figure 1A). To keep track of the position of each image within each colony, we collected low magnification overview images of the entire well prior to the high magnification imaging.

**Figure 1.**
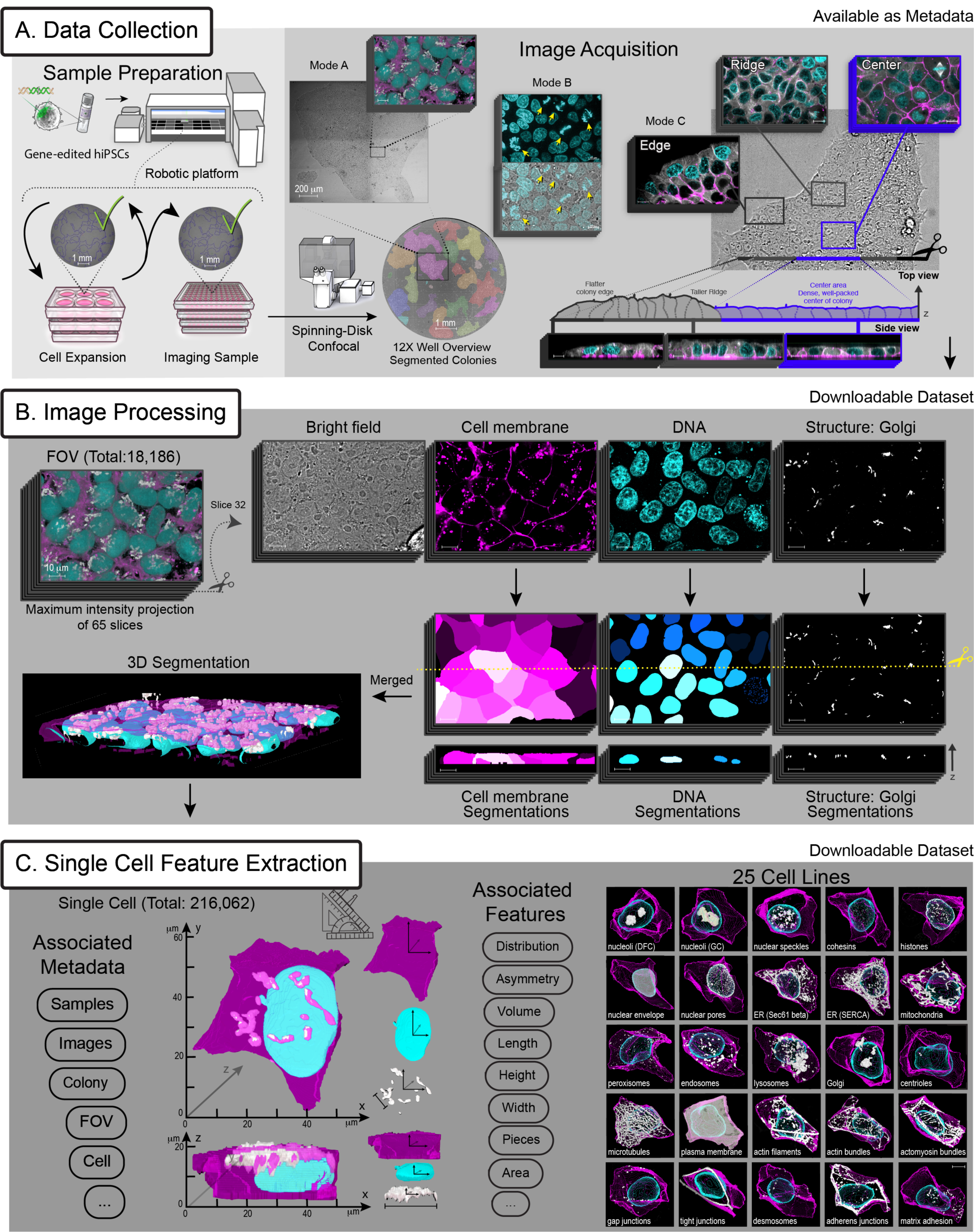
An hiPSC Single-Cell Image Dataset contains over 200,000 live, high-resolution, 3D cells spanning 25 cellular structures. The dataset was generated by a microscopy pipeline composed of three main parts; Data Collection, Image Processing and Single Cell Feature Extraction. **A)** Data Collection: the sample preparation starts with a vial of frozen gene-edited hiPSCs from a line from the Allen Cell Collection, expressing an endogenous, fluorescently tagged protein representing a particular cellular structure. The cell cultures are expanded in 6-well plates on an automated cell culture platform. At each passage cells are seeded into optical grade, glass bottom 96-well plates to create imaging samples. Bright field overview images of each well are inspected and only wells meeting pre-determined quality controls are passaged from the 6-well plates and imaged from the 96-well plates. The image acquisition of live cells starts with a 12X overview image of each well on a spinning-disk confocal microscope. Imaging sessions are conducted using three modes. In mode A, the 12X overview images of colonies are segmented by an automated script to generate sets of coordinates for positions within imageable colonies, located approximately halfway between the colony edge and colony center. Imageable colonies are those that meet size, morphology, and position-within-a-well criteria. In mode B, the microscope operator adjusts the location of the field of view (FOV) to enrich for mitotic cells via appropriate cell and DNA morphology visible with live bright field viewing and confirmed by DNA staining (yellow arrows). In mode C, three regions of colonies are imaged, the edge, ridge (just inward from the edge), and center. Cells were labeled with fluorescent DNA and membrane dyes and then imaged at each pre-selected colony position. Z-stacks were acquired at 120X in four channels, representing the bright field, cell membrane dye (magenta), DNA dye (cyan) and the fluorescently tagged cellular structure (grayscale), also shown in (B). Mode A and C panels show Golgi (via sialyltransferase) and microtubules (via alpha-tubulin), respectively. **B)** Image Processing: The hiPSC Single-Cell Image Dataset consists of a total of 18,186 curated FOVs, which are available for download. An example z-stack is shown. On the left is the maximum intensity projection of all 65 slices with all fluorescent channels combined, in the colors indicated in the panels on the right. “Cutting” the z-stack in half exposes the view of a single slice (slice 32) in the middle of the stack, shown for each individual channel, including the bright field channel. We applied 3D segmentation algorithms to each of the fluorescent channels to identify boundaries in 3D of the cells via the membrane dye (magenta), the nuclei and mitotic DNA via the DNA dye (cyan), and each of the 25 cellular structures via their fluorescent protein tag (grayscale; Golgi shown here). Resulting 3D segmentations for cell membrane, DNA, and structure channels are also shown as a side view, the xz-cross-section along the yellow dotted line. All segmentation algorithms were developed and performed using the Allen Cell and Structure Segmenter. **C)** Single Cell Feature Extraction: A total of 216,062 single cells were segmented from the FOVs. Metadata related to the sample, experiment, and microscopy was collected and associated with each individual cell. Appropriate features were extracted for each cell from the cell, the nucleus or mitotic DNA, and the cellular structure segmentations, including measurements such as the height and volume. These cells, including both the images and the segmentations as well as the metadata and features are all available for download. The hiPSC Single-Cell Image Dataset includes 25 cell lines representing key organelles and cellular structures located throughout all of the major compartments of the cell. One representative cell example per structure is shown as a 3D visualization in the 5×5 grid. Scale bars are 10 µm unless otherwise noted.

We included fluorescent cell membrane and DNA dyes to reference the locations of intracellular fluorescent protein (FP)-tagged structures relative to the cell boundary and the nucleus or mitotic chromosomes. Cells were imaged live and in 3D at high resolution (120x magnification, 1.25 numerical aperture, NA), generating 18,186 fields of view (FOVs) in four acquisition channels, representing the fluorescently tagged protein, the cell membrane and DNA dyes, and the transmitted light channel (Figure 1A&B). Measuring the locations of each of the 25 cellular structures within cells required segmentation approaches that demarcate structure, as well as the cell and nuclear boundaries within these 3D images. To do this, we used the *Allen Cell and Structure Segmenter* (the Segmenter), a fully-accessible, Python-based 3D segmentation software package (Chen et al., 2018). For each of the 25 cellular structures, we used the tagged protein to identify the location and morphology of the structure, rather than the location of the FP-tagged protein, itself (**Figure S1**). The tightly packed, epithelial-like nature of hiPSCs, as well as the need for highly-accurate 3D cell boundaries to minimize cellular structure misassignment to neighboring cells required deep learning-based segmentation approaches to create a robust, scalable, and highly accurate 3D cell and nuclear segmentation algorithm ((Chen et al., 2018) and *3D Segmentation* in Methods) applicable to all FOVs in this dataset (**Figure S1**).

From each FOV, individual cells were segmented using the plasma membrane dye, resulting in a single-cell image dataset consisting of 216,062 cells (Figure 1C). Every individual cell is labeled with a unique ID, permitting the persistent association of relevant metadata including the full set of experimental parameters, position within the original FOV, and structure segmentations with versioned software captured for future data provenance. Additional cell, nuclear, and structural features were extracted and associated with each cell ID, e.g. cellular structure volume, generating a rich single-cell image dataset for analysis. Both the FOV images and the single-cell dataset are available for use by the community as downloadable files (allencell.org/data-downloading.html) and through interactive online visual analysis tools that require no software installation or expertise (cfe.allencell.org). For the analyses described below, we used the subset of 203,737 interphase cells, excluding the 11,238 cells undergoing mitosis and a few outliers (**Table S1**).

### A PCA-based cell and nuclear shape space reveals interpretable modes of shape variation

To embrace the great diversity of these 203,737 3D images of cells spanning 25 cellular structures and directly compare cellular organization across this large population, we built a cell and nuclear shape-based coordinate system (Figure 2), adapting a standard Principal Component Analysis (PCA)-based dimensional reduction approach (Pincus and Theriot, 2007). First, we aligned all cells to their centroids, preserving both, the biologically relevant, apical-basal axis (z-axis in lab frame of reference) and the longest axis of the cell perpendicular to that axis (longest axis in x-y plane). Next, we used a spherical harmonic expansion (SHE, (Marshall et al., 1996; Ruan and Murphy, 2019)) to accurately parameterize each 3D cell and nuclear shape with a set of orthogonal periodic basis set functions, defined on the surface of a sphere. For each cell and nuclear boundary, we retained the first 16 degrees of the SHE, corresponding to 289 coefficients for each shape. The joint vector of 578 SHE coefficients for each cell was sufficient for accurate reconstruction of the original cell and nuclear shape with high spatial precision (Figure 2A&B). The joint vectors for all cells (578 SHE coefficients) were then subjected to PCA. We found that the first eight principal components represented about 70% of the total variation in cell and nuclear shape (Figure 2C). With this dimensionality reduction, the cell and nuclear shapes for each individual cell can be approximately reconstructed from a small vector with only eight components. This dimensionality reduction also organizes the cells into a simple, intuitive 8-dimensional generative “shape space”. For example, the origin (0,0,0,0,0,0,0,0) of the shape space can be reconstructed via the values of the SHE coefficients representing this location in the 8-dimensional coordinate system, and then be visualized as an idealized cell shape that statistically represents the average, or mean, shape of all of the cells in the data set (Figure 2D). Similarly, idealized shapes can be reconstructed by traversing across each of the eight orthogonal axes in the shape space.

**Figure 2.**
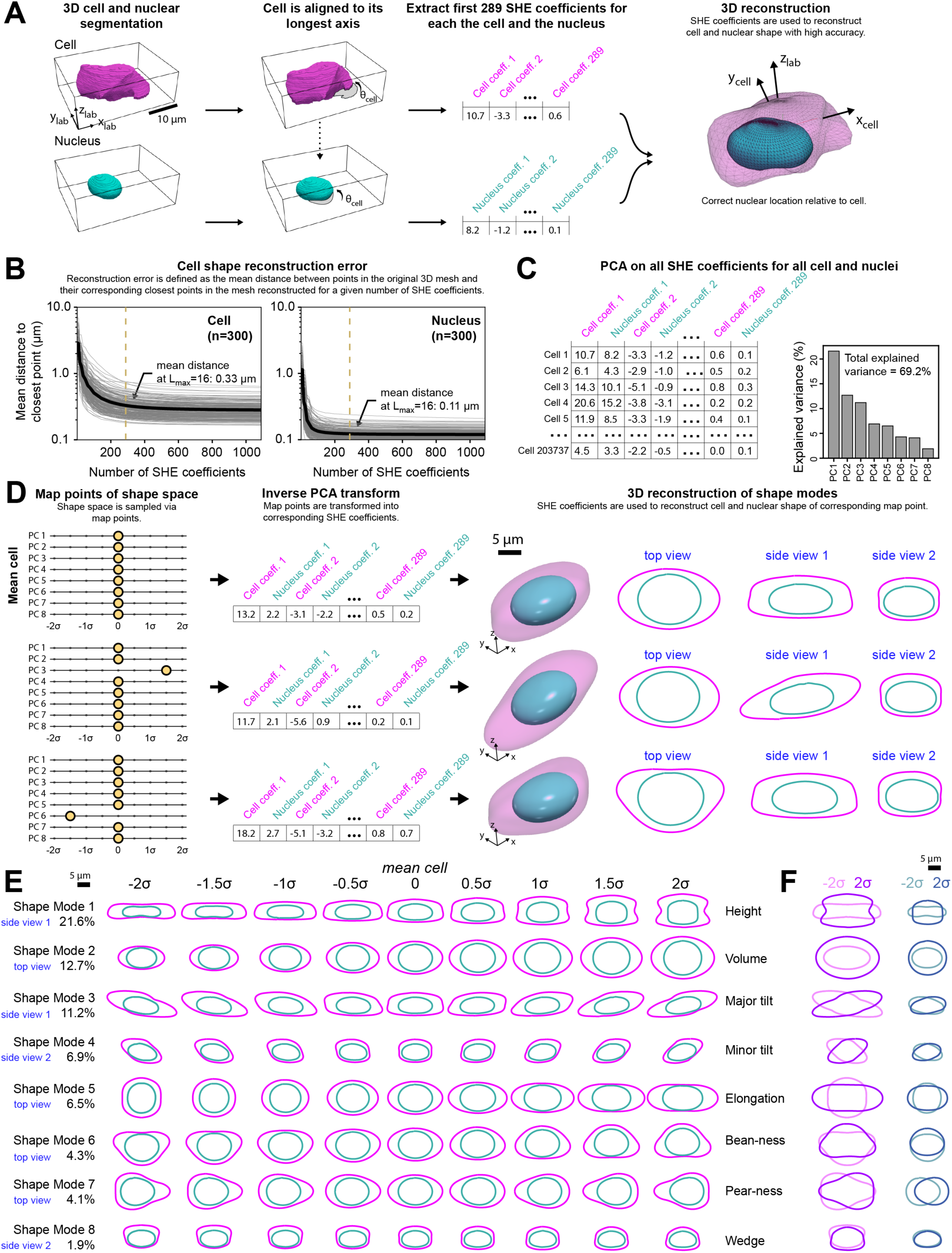
A Principal component analysis (PCA)-based cell and nuclear shape space reveals interpretable modes of shape variation in hiPSCs. **A)** Segmented 3D single-cell images of a cell and its nucleus are used as the input for a 2D alignment algorithm. The cell image is rotated in the xy-plane by θ_cell_ degrees around its centroid such that its longest axis becomes parallel to the x-axis. The same rotation angle θ_cell_ is applied to the segmented nuclear image. The resulting aligned images of the cell and nucleus are used as the input for spherical harmonics expansion (SHE) of degree L_max_ = 16 resulting in a total of 289 SHE coefficients for each the cell and the nucleus. These 578 coefficients, together, now can be used to reconstruct the cell and nuclear shape as two separate 3D meshes with high fidelity. After reconstruction, the nuclear mesh is translated to the correct position relative to the cell centroid. x_lab_, y_lab_ and z_lab_ denote the lab frame of reference and x_cell_ and y_cell_ represent the x and y coordinates in the rotated cell frame of reference. **B**) Mean distance between points in the original meshes of cell (left) and nucleus (right) to their corresponding closest points in the reconstructed meshes as the number of coefficients in the SHE increases. Each gray line is one cell (left; n=300 randomly selected samples) or nucleus (right; n=300 randomly selected samples). Black lines represent the mean. The dashed vertical lines indicate the number of coefficients for SHE degree L_max_ =16. **C)** SHE coefficients were calculated for all of the n=203,737 cells and nuclei in the analysis dataset. PCA was used to reduce the dimensionality from 2×289 SHE coefficients into the first eight principal components (PCs). **D)** Each PC was normalized into units of standard deviation, generating eight shape modes, which together are referred to as the cell and nuclear shape space. Each shape mode is sampled at nine map points. These map points are located at −2σ to 2σ in steps of 0.5σ (σ = standard deviation). Only one shape mode is permitted to vary at a time. The top example shows the mean cell and nuclear shape, represented by the map point (0,0,0,0,0,0,0,0). These nine map points for each of the eight shape modes are used as the input for an inverse PCA transform to obtain the corresponding SHE coefficients and their corresponding 3D reconstructions at these map points. The top view corresponds to an intersection between the 3D mesh of the cell and nucleus reconstructions and the xy-plane. Side views 1 and 2 correspond to an intersection between the 3D meshes and the xz- or yz-planes, respectively. **E**) 2D projections of 3D meshes obtained for each of the nine map point bins of each of the eight shape modes. The center bin in all modes is the identical mean cell shape. The most relevant of the three possible views is shown for each mode, as indicated on the far left. All three views for each shape mode and map point can be seen in **Movie S1**. Human-interpretable names for these shape modes are indicated on the right. **F**) Overlay of mesh projections of the cell (magenta) and nucleus (cyan) for the two most extremes map points (at −2σ, lighter shade and +2σ, darker shade) of each shape mode.

To build a human-interpretable understanding of the modes of shape variation in our population, we reconstructed cell and nuclear shapes at regular intervals separated by 0.5 standard deviation units along every axis of this shape space (Figure 2E, **Movie S1**). These idealized cells represent “map points” within the shape space, that can be used to identify and cluster individual real cells that are similar in shape to each idealized map point and to each other. Intuitively, these mathematically orthogonal modes of shape variation appear to describe expected variable cell shape features that are independent of one another. For example, Shape Mode 1, the mode representing the greatest amount of shape variation, appeared to largely reflect the height of the cell, and Shape Mode 2 appeared to largely reflect the overall volume of the cell (**Figure S2A**). The fact that these two biologically meaningful modes of shape variation correspond to the two top modes identified by the PCA indicates that, within this dataset, the total height of the cell is largely independent of its overall volume. Indeed, for the hiPSCs grown in self-organized colonies, the colony size and cell position within the colony appear to be the primary determinants of cell height (see “Statistical Analysis of …” section in Methods). The remaining Shape Modes 3 to 8 represented other systematic ways the shapes of these epithelial-like cells might be expected to vary, including tilting along the major or minor xy-axes (Shape Modes 3 and 4) or elongation along the major axis (Shape Mode 5). In Shape Modes 1, 2, and 5, nuclear shape changed concomitant with cell shape, while in the other shape modes, nuclear shape changed very little as the shape mode axis was traversed. Instead, for these modes it was the position and orientation of the nucleus within the cell that adjusted concomitant with cell shape (Figure 2E **and MovieS1**). For completeness, we also independently calculated shape spaces using only the overall cell shape and only the nuclear shape, which generally showed similar, biologically meaningful, modes of variation (**Figure S2B&C**).

### Building integrated average cells throughout the shape space via SHE coefficient-based parameterization and 3D morphing

The human-interpretable understanding of each of the shape modes in this cell and nuclear shape space now permits us to take advantage of the variation of cell and nuclear shape within this dataset in two significant ways. First, this standardized shape space permits clustering of similarly shaped cells, facilitating investigation of the location of cellular structures while keeping any chosen 3D spatial constraint constant. For example, we can measure how variable the locations of mitochondria are within cells of similar height. Second, the ability to analyze cellular structure location throughout this shape space permits us to consider each cell as its own “experiment” in intracellular organization, representing a particular point in the overall cell shape space comprising this normal population. Thus, we can ask how robust the location of a cellular structure may be when it is subjected to systematic variation in cell and nuclear shape. For example, we can compare differences in structure locations or their variations between flat and tall cells, small and large cells, or cells with shapes that are less or more polarized. In brief, this shape space creates an opportunity to investigate how the rules of cellular organization change in response to a set of naturally occurring shape perturbations compared to the “mean” cell shape.

To directly and quantitatively compare similarly shaped individual cells and their contents near a particular part of the shape space, we needed to map all of the possible locations of the contents of these cells into one identically bounded cell shape. Therefore, we developed a method to “morph” all of the locations of all of the points within a cell into the idealized reconstructed cell shape that best represents that cell shape (Figure 3). We took advantage of the SH expansion describing the outer cell boundary and the outer nuclear boundary and interpolated between the relevant SH coefficients. This generates a “c*ytoplasmic mapping*” of successive 3D concentric shells between the nuclear and cellular boundaries at a specified spacing. Similarly, we generated a “*nuclear mapping’*” from the nuclear centroid to the nuclear boundary. We then created a “*parameterized intensity representation”* of all of the intensity values at all of these mapped locations within the cell. This parameterized intensity representation can then be transformed back into any cell or nuclear shape (Figure 3A). As a proof of concept, we first performed this internal mapping and transformation of all of the fluorescent signal within an individual cell back into that cell’s own original shape, permitting us to measure how well spatial information is conserved using this approach. Since this internal mapping is discrete, the resultant reconstructed intensity image will have gaps, which were filled using nearest neighbor interpolation. We then calculated the voxel-wise Pearson correlation of the original and recreated images of the same cell in 3D for individual cells representing each of the 25 cellular structures. We found that for most structures this correlation was very high, above r=0.8 (**Figure S3A**). Only those structures that localized to separate discrete spots displayed slight reductions in these correlation values, likely due to the discrete nature of both the parameterized intensity representation and the structures themselves.

**Figure 3.**
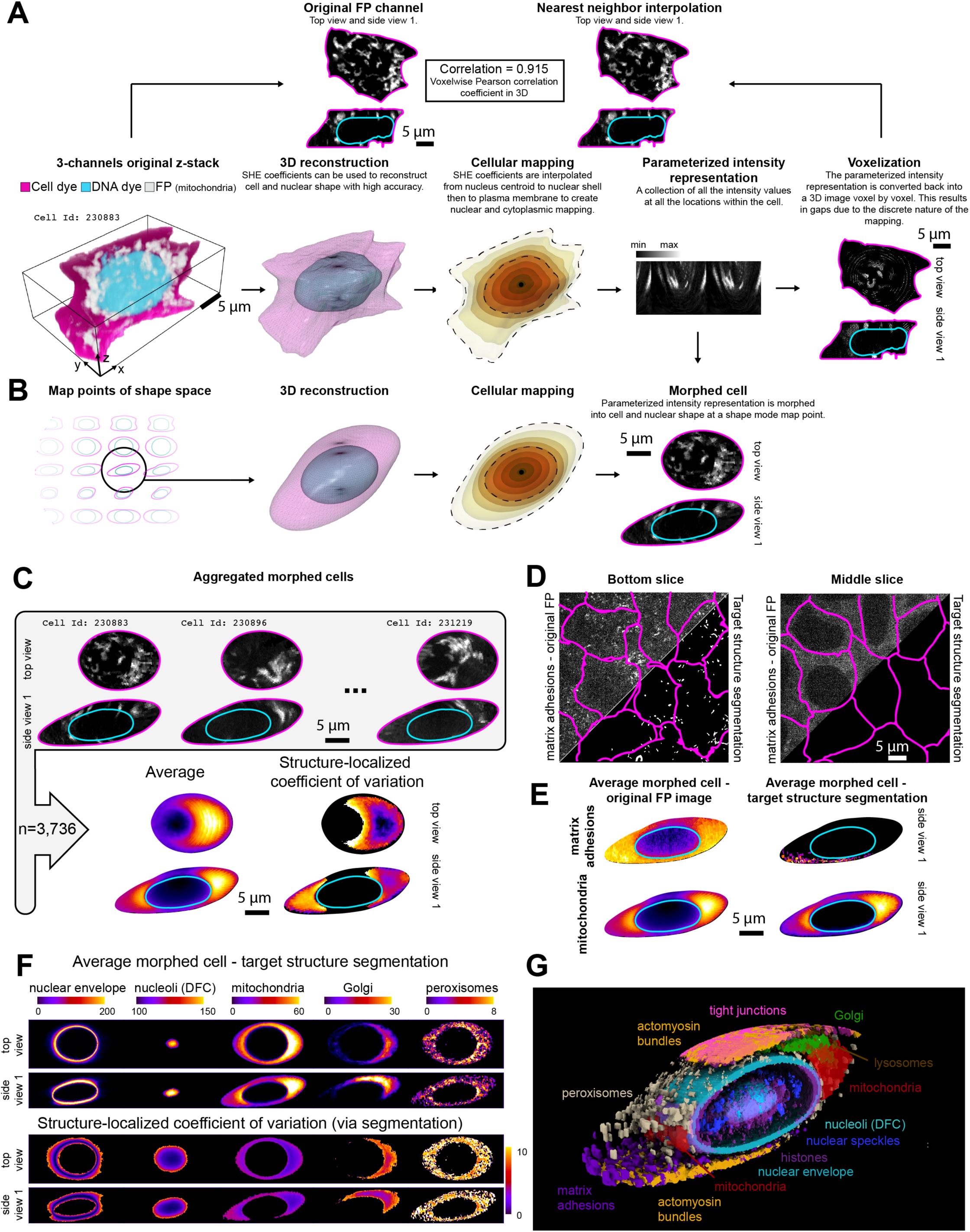
Building integrated average cells throughout the shape space via SHE coefficient-based parameterization and 3D morphing. **A)** Bottom left image, labeled as *3-channel original z-stack,* shows a 3D visualization of the original fluorescent protein (FP) intensities of tagged mitochondria (via Tom20, grayscale) in a single cell and nucleus, visualized via cell membrane dye (magenta) and DNA dye (cyan). Moving rightward along the bottom row are the steps to create the parametric intensity representation of the mitochondria via the FP signal in this cell. The second image, labeled *3D reconstruction*, shows the SHE-based 3D reconstruction meshes of the segmentations of this cell and nucleus. The third image, labeled *cellular mapping,* shows the result of interpolating the SHE coefficients to create a series of concentric mesh shells (indicated by different colors) from the centroid of the nucleus (black dot) to the nuclear boundary (inner dashed contour) to create the nuclear mapping and from that nuclear boundary to the cell boundary (outer dashed contour) to create the cytoplasmic mapping. The intensity values in the FP channel are recorded at each mesh vertex location, resulting in a *parameterized intensity representation* that is shown in a matrix format in the fourth image. This parameterized intensity representation is then converted back into a 3D image, voxel by voxel, into the same reconstructed cell and nuclear shape, shown in the fifth image, labeled *voxelization*. Here the top view and side view 1 are shown with the intensity image in the FP channel in grayscale and the cell and nuclear boundaries in magenta and cyan lines, respectively. The top left image, *labeled original FP channel*, is the top view and side view of the same cell as in the 3-channel original z-stack panel on the bottom row. The intensity image in the FP channel, in this case mitochondria), is shown along with the cell and nuclear segmentations (magenta and cyan lines, respectively). The top right image, labeled *nearest neighbor interpolation*, is the voxelized parameterized intensity representation, now with gaps filled using nearest neighbor interpolation. Voxel-wise Pearson correlation in 3D is used to compare the input image (original FP channel) with the image reconstructed via the parametric intensity representation (nearest neighbor interpolation). **B)** The same 3D reconstruction and cellular mapping procedure is now applied to a cell and nuclear shape at any map point in the shape space, shown here to the Shape Mode 3 map point (0,0,1.5σ,0,0,0,0,0). In the fourth image, labeled *morphed cell*, the parameterized intensity representation of the FP channel is morphed into this shape-space based cell and nuclear shape creating a morphed cell with morphed structure location. **C)** Top panel shows images of three different example cells with tagged mitochondria), each located near the Shape Mode 3 map point (0,0,1.5σ,0,0,0,0,0), morphed into the reconstructed cell and nuclear shape of that map point. These three and all other morphed cells with tagged mitochondria within that map point bin in the shape space can be aggregated voxel by voxel to create an average morphed cell representing the average mitochondria locations (image labeled *average*) in that part of the shape space. Morphed cells can also be aggregated by calculating the standard deviation at each voxel of the morphed cell shape (**Figure S3**). The average and standard deviation morphed cells can be combined to calculate the *structure-localized coefficient of variation*, representing a quantitative measure of how variable the location of a structure is at any given voxel. **D**) FOV images of multiple cells (cell membrane indicated by magenta lines) with labeled matrix adhesions (via paxillin) at two z positions in the z-stack. Top left triangles in each image show the original FP image. Matrix adhesions are visible near the bottom of the cells (left) but considerable FP-tagged paxillin signal is visible both at the bottom and center (right) of cells. Bottom right triangles in each image show the result of the matrix adhesion specific segmentation, in which only the matrix adhesions themselves remain near the bottom of the cells as the target of the segmentation. **E)** Average morphed cells at the Shape Mode 3 map point (0,0,1.5σ,0,0,0,0,0), representing matrix adhesions (top row) and mitochondria (bottom row) generated using either the original FP images (left column) or the target structure segmentations (right column). **F)** Top view and side view 1 of average and structure-localized coefficient of variation morphed cells at the Shape Mode 3 map point (0,0,1.5σ,0,0,0,0,0), based on the target structure segmentations, representing five distinct cellular structures. See **Figure S3** for examples for all 25 structures and **DataFile S1** for numbers of cells aggregated at each shape space bin. **G)** Eleven structures are rendered simultaneously to illustrate their relative spatial relationships in this 3D visualization (actomyosin bundles and mitochondria are labeled twice highlight their dual locations). The average morphed cells representing each of these structures at the Shape Mode 3 map point (0,0,1.5σ,0,0,0,0,0), based on the target structure segmentations were combined in this image. For each of these, the average structure image was segmented using the default *Surface* option found in the *Volume Viewer* window of ChimeraX. Thresholds for each channel were selected arbitrarily to clarify dominant localization patterns observed in the voxel intensities. See **Movie S2** for rotating image.

We next applied this approach to morph the parameterized intensity representation of each cell into the idealized cell and nuclear shape representing a nearby map point location in the shape space, creating a ‘morphed cell’ (Figure 3B). Now we can choose any map point within the shape space and identify a cluster of individual cells around that point, then create morphed versions of these cells and their contents to fit within the exact same idealized shape. In this way the location of the contents of each of these cells could be directly and quantitatively compared. The set of morphed cells within a chosen region in the shape space could also be aggregated via their parameterized intensity representations to generate an average of all of the intensities mapped within the cell and nuclear shape, or to quantify the variation in intensities via the coefficient of variation (Figure 3C).

This parametric intensity representation takes all intensities in the image into account, including any FP-tagged protein not localized to the target structure that the protein represents. For example, EGFP-tagged paxillin localized to matrix adhesions at the bottom of the cell but also throughout the cytoplasm. However, the segmentation target defined for this cell line included only the high intensity regions representing the matrix adhesions (Figure 3D). Applying this same cell morphing approach to the segmented versions of the cellular structure images (rather than to the FP images directly) permits the creation of average morphed cells containing the locations of the cellular structures that each tagged protein represents (Figure 3E). Our remaining analyses in this paper focus on the segmented structure images; but conceptually the same approach could also be applied to the raw intensity images.

We clustered all cells in the dataset arbitrarily into nine bins along each of the eight shape modes at the standard deviation intervals shown in Figure 2 (**Figure S3C**) to create a total of 65 cell shape map points (the center bin is the same in all modes), into which we morphed each of the 25 structures (Figure 3F and **Figure S3D**). By direct visual inspection, we found that the average morphed cells accurately represented the location patterns of these structures in individual, unmorphed cells (**Figure S3D**). We could then combine the average morphed locations of each of the 25 structures into the same cell shape (11 integrated structures visualized in Figure 3G and **Movie S2**), creating integrated average morphed cells, which we did for each of the 65 map point cell shapes.

### The location stereotypy of cellular structures depends on the structure but not the cell shape

To measure how variable the location of each individual cellular structure is within the cell, we used individual morphed cell images based on the structure segmentations for each structure at each map point bin in the cell and nuclear shape space. We calculated the 3D voxel-wise Pearson correlation between pairs of individual morphed cell images within a shape bin for each of the 25 cellular structures (Figure 4A) and averaged those correlation values to generate a measure of the “*location stereotypy*” of each structure (Figure 4B). Structures with a high stereotypy value have little cell-to-cell variation in their overall absolute positions for similarly shaped cells, while structures with a low stereotypy value may be found in distinct locations even for two cells whose shapes are very similar. Comparing the average stereotypy for each structure permitted us to rank structures that are most to least stereotyped in their location within the mean cell and nuclear shape. The most stereotyped structures were the nuclear envelope (lamin B1) and the plasma membrane (CAAX domain of K-Ras, “CAAX”). These observations are effectively positive controls, because these two structures should be very similar to the cell and nuclear boundary shapes that were used as fixed points in the SHE interpolation. In decreasing order of stereotypy, the next highest were two nucleolar compartments, the Dense Fibrillar Component (DFC, via fibrillarin and the Granular Component (GC, via nucleophosmin), followed by the ER (both Sec61 beta and SERCA). Structures with the least location stereotypy within the mean cell included those with a low number of discrete separated locations near the top or bottom of the cell such as centrioles (via centrin-2), desmosomes (desmoplakin), and matrix adhesions (paxillin). They were followed by slightly increased stereotypy for cohesins (SMC-1A), endosomes (Rab-5A) and peroxisomes (PMP34). To control for any effects of variable spacing within locations of each of the 25 cellular structures, we performed a systematic downsampling of the voxel size (**Figure S4A)** and found little change to the stereotypy order of the structures (**Figure S4B**).

**Figure 4.**
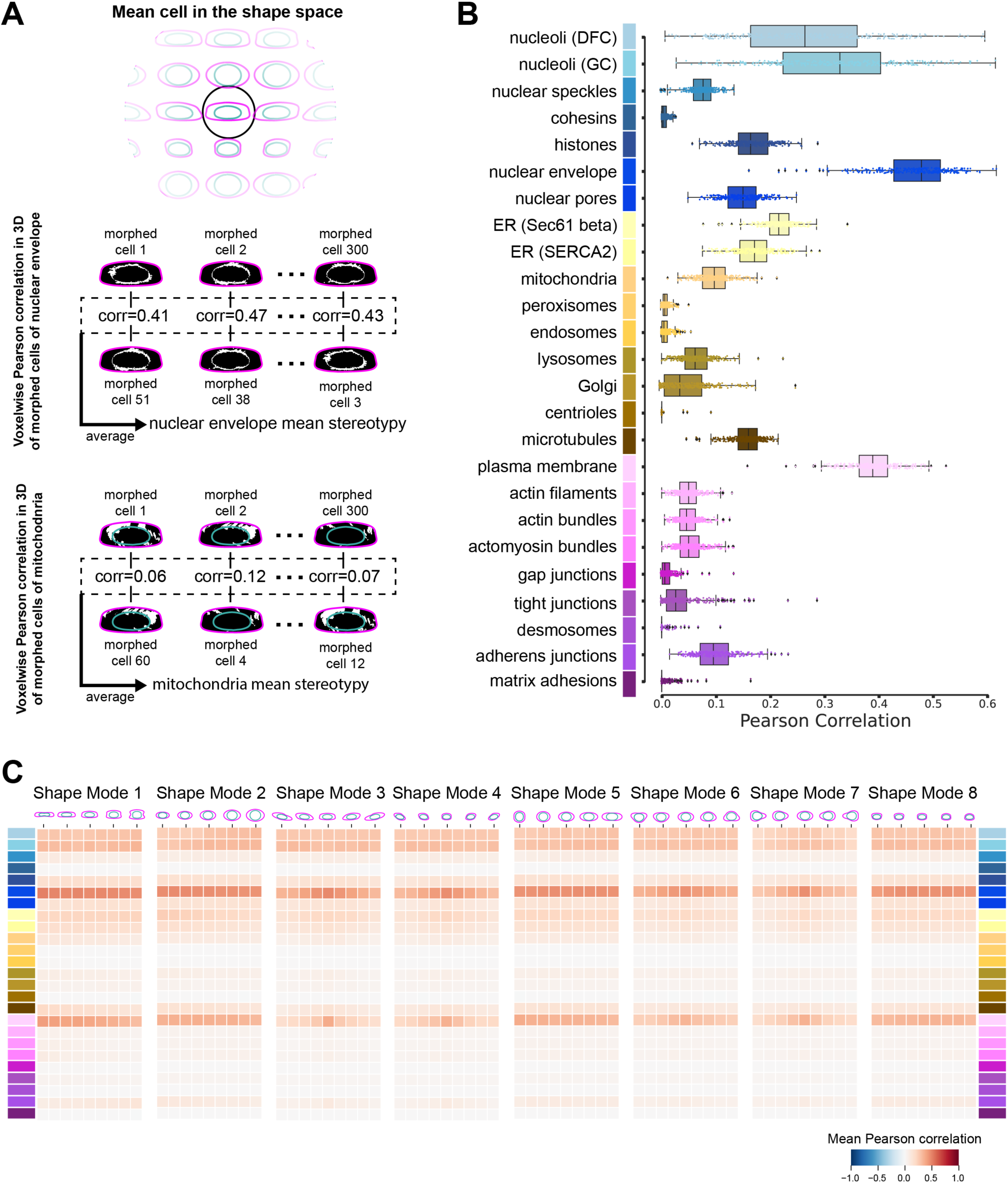
The location stereotypy of cellular structures depends on the structure but not the cell shape. **A)** Overview of the process to calculate the stereotypy of cellular structures within the mean cell shape, using the nuclear envelope (via lamin B1) and mitochondria (via Tom20) as examples. Segmented images of each cellular structure within 300 cells located in the mean cell bin in the shape space were each morphed into the mean cell shape, creating, for example, 300 nuclear envelope and 300 mitochondria morphed cells. The voxel-wise Pearson correlation was calculated for 300 unique pairs of morphed cells of same cellular structure and the results were organized as a correlation list. The mean value of the correlation list was defined as the mean location stereotypy for that structure. **B)** Box plots corresponding the values in the correlation list (see panel A) for each of the 25 cellular structures, represented by unique colors to the left of the y-axis. Dots represent the raw data (n=300), vertical black lines represent first and third quartile, boxes represent the interquartile range and the vertical black line inside the box is the mean. **C)** The process described in panel A was performed for cells in each of the 72 shape space bins to calculate the average location stereotypy for all 25 cellular structures throughout the shape space. Each heatmap value corresponds to the mean stereotypy of all 25 cellular structures for a given shape mode. Each row in the heatmap represents a different cellular structure, indicated by the same colors as in panel B. Columns in the heatmaps represent the nine binned map points along each shape mode (see **Figure S3C**). N = 300 morphed cells for each cellular structure and shape mode bin or the maximum number of cells available (see **DataFile S1** and Methods).

We next investigated how much the location stereotypy changed in response to the set of naturally occurring cell shape perturbations represented by the systematic changes in cell shape along each of the eight shape modes compared to the “mean” cell shape. Strikingly, we found very little change in the magnitude or rank order of the location stereotypy throughout the entire shape space, demonstrating that the stereotypy of all of these 25 structures was extremely robust to overall cell shape variation (Figure 4C **and Figure S4C&D**).

### The location concordance of all 25 cellular structures to each other suggests a robust, ordered compartmentalization of the cell

The analysis described above enables quantitative ranking of the cell-to-cell variation in localization of each tagged cellular structure relative to that same structure in a different cell. Also of interest is the relative similarity of the absolute localizations of each tagged structure as compared with every other structure. To measure the relationships of the locations of each of the 25 cellular structures relative to all the others, we calculated the 3D voxel-wise Pearson correlation between the average morphed cell images for all pairs of structures (pairwise structure location “concordance”) within the mean cell shape (Figure 5A). We then performed a hierarchical clustering analysis of the concordance values. This clustering is purely data-driven based on the images alone. Importantly, we found that the location concordance of these cellular structures clustered naturally into an ordered compartmentalization of the cell, from the center of the nucleus outward (Figure 5B and colors in Table 1). The four top-level clusters included structures localized to the nucleus, nuclear periphery, cytoplasm, and cell periphery, respectively. A priori, we expected to find strong concordance for several cellular structure sets, including the two nucleolar structures (DFC and GC), the two structures at the nuclear periphery (nuclear envelope and nuclear pores), the two ER tags (Sec61 beta and SERCA), and the three structures with primary localization to the apical cell-cell contacts (gap junctions (connexin-43), tight junctions (ZO-1), and desmosomes (desmoplakin)). The concordance hierarchy confirmed the expected strong concordances within each of these sets, validating this analysis approach.

**Figure 5.**
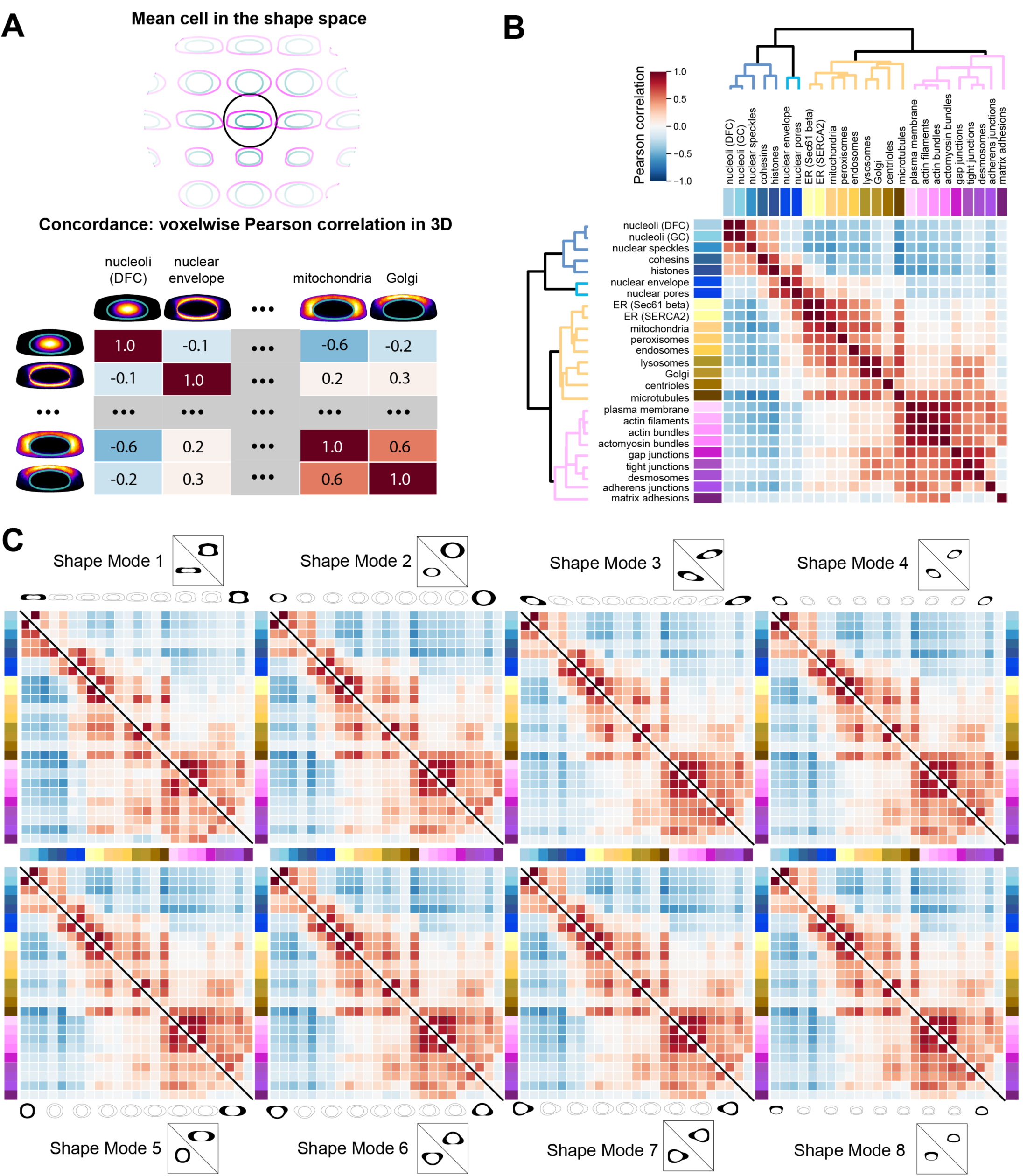
The location concordance of all 25 cellular structures to each other suggests a robust, ordered compartmentalization of the cell. **A)** Overview of the process to calculate the location concordance between all pairwise-combinations of the 25 cellular structures within the mean cell shape. The voxel-wise Pearson correlation was calculated between pairs of average morphed cells, based on the structure segmentations, morphed into the mean cell shape. This created a correlation matrix including each of the 25 cellular structures with elements of this matrix representing the location concordance between two cellular structures. **B)** The heatmap representing the location concordance for every pair of 25 cellular structures in the mean cell shape. Each heatmap value corresponds to the Pearson correlation value between the two indicated structures. The correlation matrix is used as input for a clustering algorithm to produce the dendrogram shown alongside the heatmap. Dendrogram branches are color coded according to major cell compartments (nucleus in blue, nuclear periphery in cyan, cytoplasm in orange and the cell periphery in magenta). Lengths of dendrogram branches represent the distance between clusters. **C)** The process to create the concordance heatmap for the mean cell shape in B was repeated for the reconstructed cell and nuclear shapes at the −2σ and 2σ shape space map points for each of the eight shape modes. Each heatmap represents one shape mode. The lower triangle represents shape space map point −2σ and the upper triangle represents shape space map point 2σ. For sake of clarity, diagonals are colored in white and black lines are used to separate the lower and upper triangles. The number of cells analyzed for each cellular structure and shape mode bin can be found in **DataFileS1**.

We also identified several other notably high relative concordances such as the tight concordance between lysosomes (LAMP-1) and Golgi (sialyltransferase 1), consistent with their enrichment in location in the cytoplasm near the top of the cells and the known role of Golgi in regulating lysosome localization (Hao et al., 2018; Wang and Hong, 2002). Mitochondria (Tom20) and peroxisomes (PMP34) were more tightly concordant with each other than either structure was with endosomes (Rab-5A), even though direct visual examination of individual peroxisome-tagged and endosome-tagged cells did not easily highlight this distinction. However, this observation is consistent with the known association between mitochondria and peroxisomes (reviewed in (Fransen et al., 2017).

Next, we investigated how much the concordance between all pairs of the 25 cellular structures changed in response to changes in cell shape, as described above for the stereotypy analysis. Overall, we found very little change in the hierarchical compartmentalization of these 25 structures throughout the shape space (Figure 5C, **Figure S5**). Some structures showed an overall decrease in the magnitude of concordance with other structures in the shape mode bins furthest from the mean. These structures also had greatly decreased numbers of cells available in these bins for this calculation, for example actin filaments (beta-actin) or cohesins (SMC-1A) in the furthest bins of Shape Mode 1, and so these decreases may not be biologically meaningful (**DataFile S1**).

### The impact of cell and nuclear size on the variation in cellular structure size is structure-dependent

Intracellular structures exhibit cell-to-cell variation not only in their locations but also in how much of a given structure is present in the cell. So far, we have found that neither the variability in each location of each structure in the cell nor their relative locations to each other changed much with cell volume (Shape Mode 2; Figure 5C). However, it has previously been shown that the volume of several cellular structures in the cell does correlate with overall cell volume, including the nucleus and mitochondria (reviewed in (Marshall, 2020)). We therefore used our large dataset to perform a systematic and comparative analysis of the relationship between cellular structure volume and five relevant size metrics (cell volume, cell surface area, nuclear volume, nuclear surface area, and cytoplasmic volume) for the 15 cellular structures in this dataset validated for structural volume analysis (Figure 6 and **Figure S1**). We used simple linear regression to fit the data and calculated the percent of the variation in cellular structure volumes that can be explained by each of the four cell and nuclear size metrics (“percent explained variance”; Figure 6A). The rolling average, a non-linear model fit, and analyses in which we considered the geometrical relationship between the volume and surface area of the roundest nuclei all showed similar results, validating the simple linear regression approach (Figure 6B**, Figure S6A&B,** and Methods). We found that the percent explained variance attributable to these overall cell size metrics was substantially greater for some structures, such as mitochondria (Tom20; 54%) than for other structures, such as endosomes (Rab-5A; 2%, Figure 6D&E). We also found that for nuclear structures like the nucleolar DFC (fibrillarin), more of the variance in their volumes could be explained by nuclear volume than by cell volume (77% vs. 68%, respectively; Figure 6F&G).

**Figure 6.**
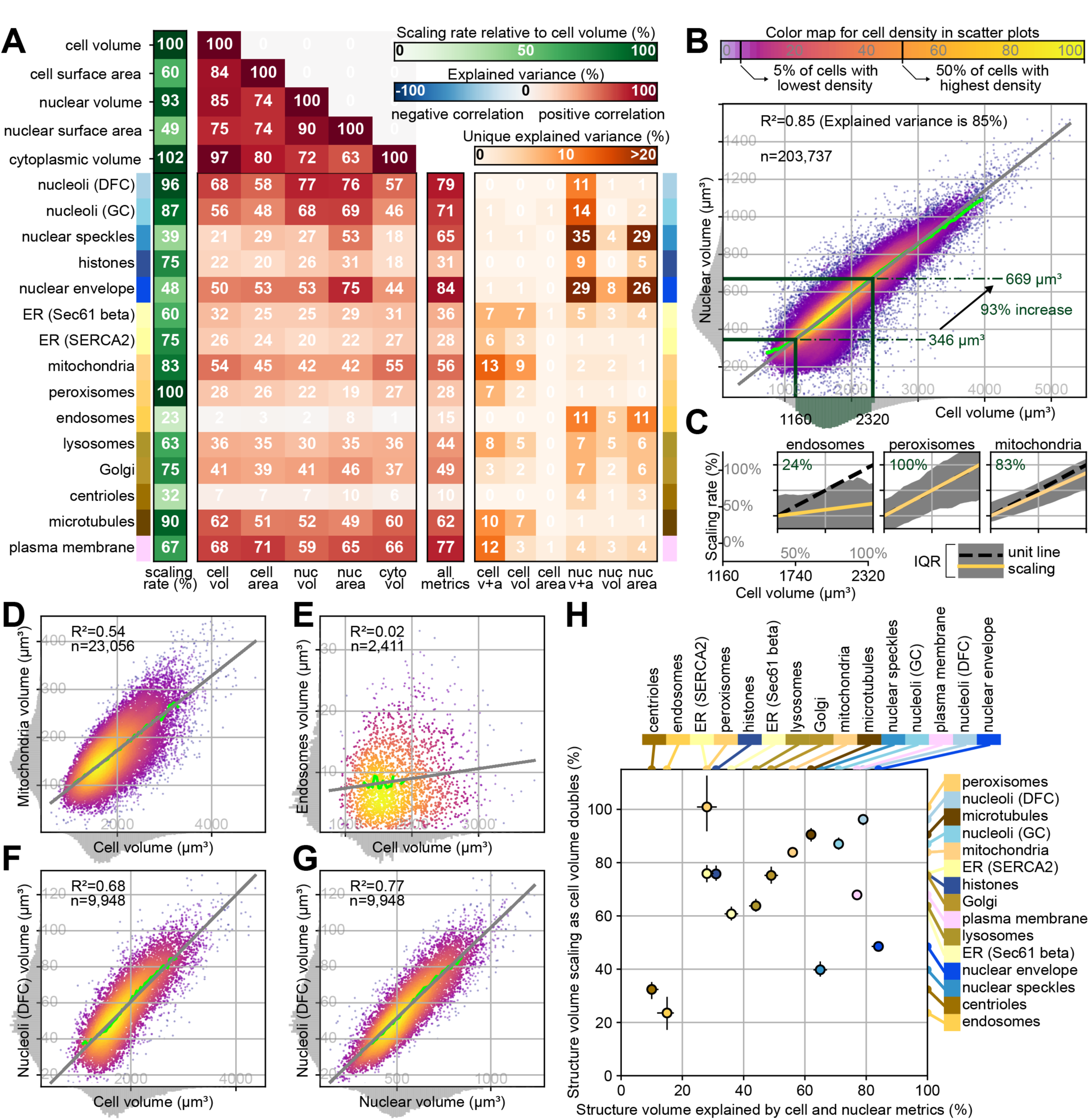
The impact of cell and nuclear size on the variation in cellular structure size is structure-dependent. **A)** Heatmap in four parts summarizing the results of a systematic, comparative analysis of the relationship between the volumes of 15 cellular structures and four cell and nuclear size metrics (cell and nuclear volume and surface area, also referred to as *Cell vol*, *Cell area*, *Nuc vol*, *Nuc area* in the heat map columns). The number of cells for each cellular structure can be found in **Table S1**. The very left of the heatmap shows the same compartmentalized cellular structure coloring scheme as in other figures. Heatmap part 1: the leftmost column, labeled *scaling rate* and colored in green indicates the percent scaling of cellular structure volumes relative to a doubling in cell volume. For example, a value of 83% for mitochondria indicates that mitochondrial volume is increased by 83% when the cell doubles in size (100%) (B and C). Heatmap part 2: the next five columns, each labeled with the four cell and nuclear size metrics plus cytoplasmic volume (the difference between cell and nuclear volume, labeled *Cyto vol*) are colored with the blue-red heatmap color range, labeled *explained variance*. These columns indicate the percent of variation in each cellular structure volume (and surface area for the cell and nucleus; rows) that can be statistically explained using the metrics indicated in each column. Negative values would represent a negative correlation relationship between the two variables (row and column), but are not present in this heatmap. These percent of explained variance values are a measure of the tightness of the coupling between cellular structure volume and specific cell and nuclear size metrics. Heatmap part 3: the center single column, labeled *all metrics*, uses a multivariate model that includes the four cell and nuclear size metrics. The values in this heat map column represent the total percent of variation in each cellular structure volume that can be statistically explained using a combination of all four metrics. Heatmap part 4: The last six columns colored with the orange heatmap color range, labeled *unique explained variance*, show the percentage of variation in each cellular structure volume that can be uniquely attributed to a single metric (each of the four cell and nuclear size metrics) or a pair of metrics (cell volume plus surface area as the cell size metrics – labeled *cell v+a*, nuclear volume plus surface area as the nuclear size metrics, labeled *nuc v+a*). This number is computed as the difference between the total explained variance (the all metrics column) and the variance explained by a model using all four cell and nuclear size metrics except for the metric (or pairs of metrics) indicated in that column. For example, the second orange heat map column, labeled *Cell vol*, indicates the percentage of explained variance that is lost when cell volume is removed from the multivariate model. Thus, this is the percent of explained variance that can be uniquely attributed to cell volume. **B)** Scatterplot comparing cell volume (x-axis) and nuclear volume (y-axis) across all cells (n=203,737). Cells are colored based on an empirical density estimate. The green line is a running average. The gray line depicts a linear regression model where variation in the nuclear volume (y-axis) is explained as a linear function of the cell volume (x-axis). The explained variance (R^2^) in nuclear volume is 85% as stated in the top-left of the plot. The linear regression model is also used to calculate the scaling rate, i.e. how much larger (in %) nuclear volume is when cell volume doubles. Specifically, the regression model is evaluated for the cell volume interval from 1,160 to 2,320 µm^3^ (where the cell volume doubles) to determine to scaling percentage for nuclear volume. **C)** Line plots showing the relative volume scaling rate for three cellular structures (endosomes, peroxisomes and mitochondria) over the same cell volume doubling range as in **B**, from 1,160 to 2,320 µm^3^. The yellow lines represent the scaling rate, also indicated by the numbers in the top left corner of each of these plots. The regions filled in gray represent the interquartile range (IQR) measured across cells that were binned in 10 cell volume bins (y-axis). The xy-axes to the far left are used to indicate the values of the tick marks in each of the three plots. **D-G)** Similar scatterplots as in (B), correlating the volumes of mitochondria (D), endosomes (E), nucleoli (DFC, F and G) with either cell or nuclear volume (x-axes) along with statistical measures. **H)** Scatterplot comparing the total percent explained variance (x-axis contains the values in the all metrics column of the heatmap in (**A**) and the relative volume scaling rate (y-axis contains the values in the scaling rate column in the green heatmap in (**A**) across all of the 15 cellular structures. The error bars depict the 5-95% confidence intervals using a bootstrap analysis. The markers along the top and right side of the plot indicate the ranked order of the structures for that metric. Compartmentalized cellular structure coloring scheme is used to help identify specific structures.

Each of the five cell and nuclear size metrics themselves, of course, also correlate with each other (Figure 6A), thus obscuring their potential to independently explain the variance in the volumes of the 15 cellular structures. To disentangle these correlations, we applied a multivariate model and calculated the total percentage of the variance explained for each of these structures by the combination of all four direct cell and nuclear size metrics (‘total explained variance”; see ‘all metrics” column in Figure 6A). For all but two of the cellular structures, the total explained variance was at least 28%; but this percentage varied widely depending on the structure (x-axis in Figure 6H). At the lowest end were the centrioles (centrin-2), which we expected to be very small as they are discrete structures that should not get bigger as cells grow, and thus invariant with all size metrics. At the highest end were the nuclear envelope (laminB1) and the plasma membrane (CAAX), which we expected would correlate well with nuclear and cell surface areas, respectively. Notably, the volumes of all three nuclear body structures (nucleolar DFC, GC, and speckles) were the next-most tightly correlated to the optimal linear combination of cell and nuclear size metrics.

We then used the multivariate model to calculate the *unique* contributions of both cell size metrics vs. both nuclear size metrics vs. the unique contributions of each of the four metrics individually (Figure 6A). For all five nucleus-related structures, the variance in structure volume was better explained by nuclear size metrics than by cellular size metrics. For the nuclear envelope, more of the variance was uniquely attributable to the nuclear surface area than nuclear volume; this anticipated result confirmed the validity of this approach. Unexpectedly, the variance in nuclear speckle (SON) volumes was most uniquely attributable to the nuclear surface area and not the nuclear volume, although speckles localize throughout the nucleoplasm.

Of the cytoplasmic structures, microtubules (alpha-tubulin, see **Figure S1** for target segmentation of microtubule bundles), which localize throughout the cytoplasm (**Figure S5D**), had the highest percent variance explained by the optimal combination of the four size metrics, followed next by mitochondria, Golgi and lysosomes (x-axis in Figure 6H). Endosomes (Rab-5A) had one of the lowest percent explained variance values, almost as low as centrioles, even though they are spread out throughout the cytoplasm. For the cytoplasmic structures, some variation in their structure volumes was uniquely attributable to either cell or nuclear metrics; but in all cases the unique contribution of cell surface area on its own was negligible. While nuclear structures seem to be most tightly coupled to nuclear size metrics, cellular structures range more widely in how well the variance in their volumes was uniquely attributable to cell versus nuclear size metrics. We explored whether cell and nuclear shape might explain some of the variation in cellular structure volumes but found contributions from other shape modes to be negligible (**Figure S6C**). Overall, these results show that how well cell and nuclear size metrics account for the variation in cytoplasmic structure volumes is structure-dependent, consistent with the wide range of cell functions that these structures regulate.

We also measured the relative volume “*scaling rates*” for each of these 15 cellular structures as the percentage increase in structure volume given one doubling in cell volume over a range that was well represented in our cell population, specifically from 1160 µm^3^ to 2320 µm^3^ (Figure 6A-C). For example, the volume of mitochondria increased by an average 83% (from 108 to 198 µm^3^) for this doubling in cell volume (an increase of 100%). The structures with the greatest relative scaling rates were the peroxisomes (via PMP34), followed closely by both nucleolar structures and then microtubules (y-axis in Figure 6H), all of which nearly doubled in structure volume with the doubling of cell volume. The structures with the lowest relative volume scaling rates were also the structures identified as having the lowest explained variance, that is the endosomes and centrioles. For most structures, however, we observed relative scaling rates of at least 60%, consistent with the simple expectation that larger cells typically would also have larger organelles. We observed lower scaling rates for the two structures whose volumes correlated most strongly with nuclear surface area, the nuclear envelope and nuclear speckles. This is consistent with surface area generally scaling less quickly than volume, for example, doubling the size of a perfect sphere leads to only a 59% increase in its surface area. The peroxisomes stood out as exhibiting an unusual pattern of both a high relative volume scaling rate and great variability in peroxisome volume from cell to cell. This systematic analysis of the relationship between cellular structure volume and cell and nuclear size metrics creates a rich set of quantitative constraints for modeling intracellular organization.

### Downsampling the hiPSC Single-Cell Image Dataset demonstrates generalizability of this multi-part analysis approach

The over 200,000 individual cells spanning 25 cellular structures within the hiPSC Single-Cell Image Dataset permitted us to develop this multi-part, data-driven, computational approach that generates a collection of quantitative rule-building constraints for the locations, amounts, and degree of variation of a set of cellular structures within a population of 3D cell images. The four outputs of this approach are 1) a 3D cell and nuclear shape space with human-interpretable orthogonal shape modes; 2) measurements of the location stereotypy throughout the shape space; 3) measurements of the location concordance between all pairs of cellular structures throughout the shape space; and 4) measurements of the variation in the volumes of the cellular structure relative to cell and nuclear size. With this approach we have created a fundamental benchmark for comparison of these analyses of healthy, normal, undifferentiated hiPSCs with future studies of other populations of cells in different cell states, including differentiated cells, or cells in pathological states generated by pharmacological or genetic perturbations. However, broader adoption of this quantitative approach will depend on how generalizable these analyses are to other kinds of data sets, particularly those that have a substantially smaller number of cells.

We therefore assessed the minimal dataset size required to maintain the scientific conclusions of each of these four analyses (Figure 7). For the cell and nuclear shape space, seven of the first eight shape modes were clearly recapitulated with just 300 randomly chosen segmented cells and nuclei (Figure 7A). The location stereotypy was greatly invariant to sample size (Figure 7B); the correct location stereotypy rank order of all 25 structures could be entirely recapitulated with just 35 pairs of cells per structure per shape space bin. Next, we calculated the cellular structure pairwise concordance based on a random sampling of 300 cells per structure within the mean cell bin and found that only three structures changed their location in the hierarchical clustering dendrogram (Figure 7C). For these two analyses, sufficient cells would need to be imaged to ensure the required number of cells per desired shape space bin. Similarly, we repeated the full set of statistical analysis of cellular structure volume variation with a systematic reduction in numbers of cells per structure and found that we could recapitulate all of the biological observations reported above with 300 cells per structure (Figure 7D). Overall, each of the four analyses could be successfully performed based on numbers of cells that could be reasonably collected by a single investigator imaging on a standard laboratory microscopy over the course of a few days.

**Figure 7.**
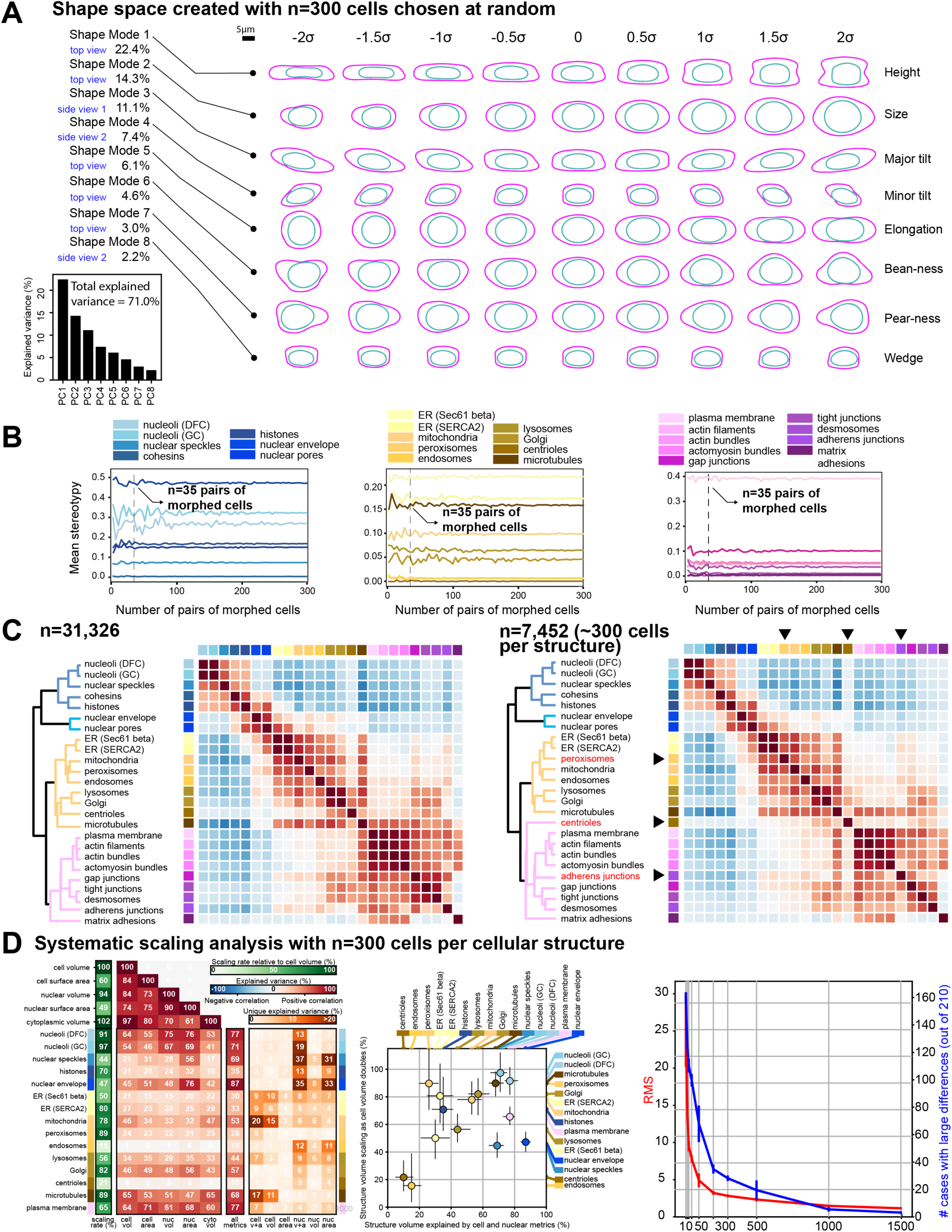
Downsampling the hiPSC Single-Cell Image Dataset demonstrates generalizability of this multi-part analysis approach. **A)** Cell and nuclear shape space generated with 300 randomly selected cells from the analysis dataset. The figure shows 2D projections of 3D meshes obtained for each of the nine map point bins of each of the eight shape modes. The center bin in all modes is the identical mean cell shape. The most relevant of the three possible views is shown for each mode, as indicated on the far left. Humanly interpretable names for these shape modes are indicated on the right. PCA was used to reduce the dimensionality from 2×289 SHE coefficients into the first 8 PCs. Total explained variance by each component is shown in the bottom left plot. **B)** Mean stereotypy as a function the number of pairs of morphed cells used to compute the voxel-wise Pearson correlations. Cellular structures are grouped into three clusters: nucleus and nuclear periphery, cytoplasm and cell periphery. Dashed vertical lines indicate the minimum number of pairs of cells necessary to recover the stereotypy ranking shown in **Figure S4**. **C)** The left heatmap represents the location concordance for every pair of 25 cellular structures in the mean cell shape as in Figure 5B computed on all n=31,326 cells within the mean cell shape bin. The right heatmap represents the location concordance for every pair of 25 cellular structures in the mean cell shape calculated on a dataset composed by 300 randomly chosen cells per structure (except n= 252 for nuclear speckles, see **DataFile S1**) for a total number of n=7,452 cells within this mean cell shape bin. Each heatmap value corresponds to the Pearson correlation value between the two indicated structures. The correlation matrix is used as input for a clustering algorithm to produce the dendrogram shown alongside the heatmap. Dendrogram branches are color coded according to major cell compartments (nucleus in blue, nuclear periphery in cyan, cytoplasm in orange and the cell periphery in magenta). Lengths of dendrogram branches represent the distance between clusters. Dendrogram on the left and right side only differs by the cluster assignment of three cellular structures, the peroxisomes, centrioles, and adherens junctions, highlighted in red and marked with arrowheads. **D)** Systematic scaling analysis with n=300 randomly chosen cells per cellular structure. The Heatmap on the left is the equivalent to FigureScalingA and the scatterplot in the center is the equivalent of Figure 6H, but with the downsampled number of cells per structures. The plot on the right shows the effect of downsampling the dataset on the complete set of statistical measurements calculated and shown in the heatmap on the left. Here, the original set of measurements were compared to a new set of measurements calculated on a series of downsampled versions of the dataset with n cells per structure randomly selected (n=10, 20, 30, 50, 100, 200, 300, 500, 1,000, 1,500; x-axis), repeated three times. The root-mean-square (RMS) difference between the two sets of measurements is shown in red and the number of cases, out of 300, where the absolute difference between the measurements was larger than 5% is shown in blue. A dataset size of 300 cells lies near the inflection point of both metrics.

## Discussion

A major goal of cell biology is to determine how a subset of expressed genes dictate cellular phenotypes. To address this enormous challenge, we must develop approaches that can reduce the amount and complexity of the information contained in cell behaviors. While many others are approaching this using genomics and proteomics, our strategy is to approach this question from the perspective of cellular organization, because it is both a key readout and driver of cell behavior. We choose a dimensionally-reduced approach by focusing on the level of the major cellular structures. To do this, we developed new ways to convert raw image data of cells and their structures into dimensionally reduced information in a form that both summarizes the raw data and embraces the vast cell-to-cell variability observed even within a population of putatively identical cells. Our work toward this goal has included creating a collection of isogenic cell lines with FP-tagged cellular structures (the Allen Cell Collection), building and standardizing an automated microscopy imaging pipeline, creating new deep learning image processing algorithms, generating a large high-quality dataset, inventing a new multi-part analysis approach that generates a collection of quantitative rule-building constraints (rules), and building tools to facilitate the democratization of these results such as an online 3D viewer to access the data. With this approach we determined where, how much and how variable the various cellular structures are in an integrated and holistic manner. The initial objective is to identify quantitative relationships that can become rules of organization, and then to use these data to eventually develop and improve biological models and ultimately laws for understanding and predicting cellular organization in a wide variety of biological contexts.

This grand plan requires high-quality image data of cells and their structures. We introduce such a dataset here: the hiPSC Single Cell Image Dataset with over 200,000 live cells in 3D and spanning 25 major cellular structures. The scale and quality of this dataset permitted us to create a multi-part, data-driven, generalizable approach that transforms the image data into a dimensionally reduced and human-interpretable set of quantitative rule-building constraints for the locations, amounts, and degree of variation of each of these 25 cellular structures. These constraints comprise a quantitative benchmark for comparison to other cell types and states, e.g., those observed during hiPSC differentiation as they transition from their epithelial-like state into more mesenchymal-like cells, with consequent differences in intracellular organization. Second, these constraints are inherently quantitative and therefore can be used to both produce and test simulations and models of cell organization. For example, models that predict the organization of sets of cellular structures based on conceptual or mechanistic constraints would need to recreate the location, variability, amount, and relative positional dependencies of these structures. Comparing these quantitative rule-building constraints with those of cells in other cell states and testing how underlying mechanisms can generate these rules via models and simulations, will permit us to move towards a deeper understanding of cell organization, aspiring to find general principles.

The analysis approaches presented here reveal some initial glimpses into new biologically interesting phenomena. The cell and nuclear shape space that represents the hiPSC Single-Cell Image Dataset revealed that we can reduce the vast complexity of 3D cell and nuclear shape into a human-interpretable understanding of the mean cell shape and the variation around it. It also creates a coordinate system by which we can now cluster groups of similar cells for further analysis and identify outliers. This shape space exposes a relationship between the behavior of the overall cell shape and the nuclear shape, which deserves deeper investigation. Next, the methods we developed to morph cells and their cellular structures together within clusters of similarly shaped cells permit both the analysis of cellular structure stereotypy and concordance. In principle, the overall concordance among structures can span a range. At one extreme, all structures could be coupled, e.g. every structure depending on every other structure; whereas at the other extreme, every structure could be independent from every other structure. We found that the location concordance of all the cellular structures clustered naturally into an ordered compartmentalization of the cell, from the center of the nucleus outward. Then we tested whether this result was valid for only a particular cell shape, e.g., the mean cell shape, or changed with systematic changes in that shape. We were surprised at how robust both the stereotypy and the concordance proved to be across all of the cell shape variation in our population.

Our systematic analysis of how cellular structure volume relates to cell and nuclear size also raises interesting questions. The apparent lack of correlation between endosome volumes and cell and nuclear size could arise in several ways. First, endosome size may just depend on the size of other cellular structures, highlighting a need for experimental data that includes the size of other structures, going beyond just the cell and nuclear size. Second, we use Rab-5A as the marker for endosomes; but it marks only a subset of endosomes, the early endosomes, and perhaps different subsets of endosomes relate differently to cell and nuclear size. Interesting questions also follow from our observation that the variation in the volumes of all nuclear structures was better explained by nuclear size than cell size metrics; and furthermore, the nuclear surface area was more tightly coupled to these structures than to nuclear volume. Nuclear speckles, for example, exhibited a surprisingly strong relationship with nuclear surface area. This is intriguing in light of the possible connection between transcript splicing (which occurs at nuclear speckles) and increased rates of nuclear export (Valencia et al., 2008). This is the first time that the relationship between cellular structure size and cell and nuclear size has been compared among so many different cellular structures all in the same consistent experimental system.

Transcriptomics and proteomics are having a great impact in understanding how the “building blocks” of the cell generate and regulate cell behavior and disease. Recent single-cell versions of these studies, particularly for gene expression, together with dimension reduction approaches to statistically identify and separate groups of similar individual cells, have permitted new insights into different cell types and states. Our studies add a new dimension to these analyses by incorporating the spatial organization of cell structures, that is, where and when these parts come together in space and time to drive that function. Our approach relies on the well-established tight linkage between 3D cell organization and cell function, aspiring to use this to identify cell types and states from images, and relate them to single cell gene expression profiles (Gerbin et al., 2020). These approaches were built with live cell imaging in mind and thus are poised to incorporate dynamics. Recently, studies combining quantitative measures of sarcomere organization with gene expression in the same individual cardiomyocytes demonstrates the importance of incorporating the spatial cell organization metrics for a more complete classification of cell states (Gerbin et al., 2020).

Other recent systematic image-based approaches have catalogued the location of human proteins in several cell types and used protein and structure locations within cells to identify differences in intracellular spatial patterns among cells in distinct states (Caicedo et al., 2017; Gut et al., 2018; Thul et al., 2017). Our work complements these approaches with its focus on analyses of 3D cell organization at the level of cellular structures, and on the generation of quantitative measurements in a human-interpretable manner. Taken together, these studies bring a critical missing dimension, i.e., the spatio-temporal component, to the single cell revolution (Aldridge and Teichmann, 2020). Our study furthers this community goal by adding critical tools, data, and analyses that show the importance of studying large populations of cells and embracing their variations to further our understanding of the underlying rules that organize cells. As part of our mission, we aspire to democratize this emerging area of research; the full image dataset and analysis algorithms introduced here, as well as all the reagents, methods, and tools needed to generate them, are shared in an easily accessible way (Allencell.org). This data is available to all for further biological analyses and as a benchmark for new development of tools and approaches moving towards a holistic understanding of cell behavior.

## Acknowledgments

We thank Joan Brugge, Gaudenz Danuser, Michael Elowitz, Tom Goddard, Quincey Justman, Jennifer Lippincott-Schwartz, Wallace Marshall, Sean Palacek, Zach Pincus, and Tom Pollard for helpful scientific discussions. The WTC line that we used to create our gene-edited cell lines was provided by the Bruce R. Conklin Laboratory at the Gladstone Institute and UCS. We wish to acknowledge the Allen Institute for Cell Science founder, Paul G. Allen, for his vision, encouragement and support.

## Author Contributions

J.M.B., J.A.C., J.C., T.P.D., M.A.F., N.G., K.A.G., B.W.G., R.N.G., A.H., M.C.H., C.H., A.R.H., G.T.J., J.J.L., A.N., A.M.N., L.P., S.M.R., M.M.R., B.R., L.M.S., M.S., J.S., J.E.S., J.A.T., D.J.T., D.M.T., A.P.T., V.V., M.P.V., W.W., C.Y. contributed to the development and/or design of the methods used in this paper. M.B., J.M.B., J.A.C., B.C., J.C., Z.J.C., S.D., S.D., T.P.D., R.M.D., T.J.F., G.F., T.G., L.J.H., H.C.H., E.J.I., G.T.J., G.R.J.,B.K., J.J.L., G.E.M., S.L.M., K.M., A.N., L.P., T.A.P., M.M.R., L.M.S., M.S., J.S., S.S., M.F.S., M.J.S., D.J.T., D.M.T., R.V., M.P.V., W.W., T.W., C.Y., R.Y. contributed to the software developed to create, assess, and analyze the dataset. J.E.A., A.B., J.A.C., J.C., M.E.C., S.Q.D., M.A.F., N.G., K.A.G., T.G., B.W.G., A.H., C.H., J.J.L., H.M., I.A.M., A.N., A.M.N., L.P., S.M.R., M.M.R., B.R., L.M.S., J.E.S., D.J.T., D.M.T., A.P.T., V.V., M.P.V., C.Y., R.J.Z. contributed to validation of the cell lines used, images taken and data that lies herein to ensure reproducibility. J.E.A., J.M.B., J.A.C., S.Q.D., R.M.D., M.A.F., K.A.G., T.G., R.N.G., A.H., G.R.J., T.A.K., J.J.L., H.M., L.P., S.M.R., M.M.R., B.R., L.M.S., J.S., Y.S., D.M.T., A.P.T., V.V., R.V., M.P.V., C.Y., R.J.Z. contributed to the formal analysis that is shown in main figures and supplementary materials. J.E.A., A.B., S.C., M.E.C., C.M.D., S.Q.D., M.A.F., N.G., J.L.G., K.A.G., T.G., B.W.G., A.H., M.C.H., C.H., W.W.L., S.A.L., H.M., I.A.M., A.N., A.M.N., L.P., S.M.R., B.R., J.E.S., W.J.T., J.A.T., D.J.T., A.P.T., V.V., M.P.V., C.Y., R.Y., R.J.Z. contributed to collecting the data, performing the experiments or analyzing the imaging data. B.B., J.M.B., J.A.C., J.D., M.A.F., B.W.G., A.H., A.N.L., R.M., T.L.M., M.M.R., B.R., J.S., J.E.S., D.M.T., W.W. contributed to providing the reagents, materials or compute resources needed to accomplish this study. J.E.A., M.B., B.B., A.B., J.M.B., S.C., J.C., M.E.C., C.M.D., S.Q.D., R.M.D., T.J.F., M.A.F., N.G., J.L.G., K.A.G., T.G., B.W.G., A.H., M.C.H., C.H., T.A.K., W.W.L., J.J.L., S.A.L., K.M., I.A.M., A.N., M.M.R., B.R., J.S., S.S., J.E.S., M.J.S., W.J.T., D.J.T., D.M.T., A.P.T., C.Y. contributed to management activities to annotate, scrub data, and maintain research data. J.C., M.E.C., T.P.D., N.G., C.H., A.R.H., G.T.J., T.A.K., A.N., S.M.R., J.A.T., R.V., M.P.V., C.Y. helped prepare, create and present the original draft of this work. A.B., J.C., K.R.C.M, T.P.D., C.L.F., N.G., C.H., A.R.H., G.T.J., T.A.K., I.A.M., S.M.R., J.A.T., R.V., M.P.V., C.Y. helped provide critical review and editing of this work. J.A.C., K.R.C.M, T.P.D., N.G., G.T.J., T.A.K., M.M.R., D.M.T., R.V., M.P.V. helped prepare the visualization of the work described, specifically via visual tools, resources, and data presentation. J.E.A., K.R.C.M, M.A.F., N.G., K.A.G., R.N.G., A.H., A.R.H., G.T.J., T.A.K., I.A.M., A.M.N., S.M.R., B.R., D.M.T., W.W. provided oversight and leadership, including planning and execution of the cross-team project management. K.R.C.M, R.N.G., A.H., G.T.J., I.A.M., S.M.R., S.S., D.M.T. provided management, coordination of the project and administration of resources.

## MATERIALS AND METHODS

### RESOURCE AVAILABILITY

#### Lead Contact

Further information and requests for resources and reagents should be directed to and will be fulfilled by the Lead Contact, Susanne Rafelski.

#### Material Availability

Using the Wild Type WTC-11 hiPSC line background (Kreitzer et al., 2013), we previously generated the Allen Cell Collection of hiPSC lines in which each gene-edited cell line harbors a fluorescent protein endogenously tagged to a protein representing a distinct cellular structure of the cell (Roberts et al., 2017b). Fifteen additional Allen Cell Collection lines were generated using the same methods in this study. The cell lines are described at https://www.allencell.org and are available through Coriell at https://www.coriell.org/1/AllenCellCollection. For all non-profit institutions, detailed MTAs for each cell line are listed on the Coriell website. Please contact Coriell regarding for-profit use of the cell lines as some commercial restrictions may apply.

#### Data and Code Availability

The Datasets generated during this study are available at Quilt as packages:

- Full dataset: https://open.quiltdata.com/b/allencell/packages/aics/hipsc_single_cell_image_dataset
- 12X colony dataset: https://open.quiltdata.com/b/allencell/packages/aics/hipsc_12x_overview_image_dataset
- Supplementary MYH10 repeat dataset: https://open.quiltdata.com/b/allencell/packages/aics/hipsc_single_cell_image_dataset_supp_myh10
- Tutorials and demo for how to access the data for different purposes: https://github.com/AllenCell/quilt-data-access-tutorials
- Original/source data for figures in the paper are available in Github: https://github.com/aics-int/cvapipe_figure_notebooks

The code supporting the current study has been deposited in Github repositories and the released code repositories and packages use the following packages in parts, including Numpy (Harris et al., 2020), Scipy (Virtanen, 2020), Napari (Nicholas Sofroniew et al., 2019), Seaborn https://seaborn.pydata.org/citing.html, Py Torch (Paszke et al., 2019), ITK (McCormick et al., 2014), pandas (McKinney, 2011), matplotlib (Hunter, 2007), and label free (Ounkomol et al., 2018):

- Image segmentation code, trained models, and demo Jupyter notebooks have been released at https://github.com/AllenCell/segmenter_model_zoo.
- Segmentation code used to reproduce structure segmentations from a set of algorithms to choose from, each with restricted numbers of parameters to tune are available at https://github.com/AllenCell/aics-segmentation.
- Mitotic image classifier code (Falcon and Cho, 2020; Paszke et al., 2019), (for both training and testing) and all trained models are available at https://github.com/AllenCell/image_classifier_3d.
- Code used to generate contact sheets for quality control single-cell visualizations of all segmented cells is available at https://github.com/AllenCellModeling/actk
- Code used for feature calculation:
  - aicsfeature (https://github.com/AllenCell/aicsfeature)
  - spherical harmonics parameterization (https://github.com/AllenCell/aics-shparam)
  - cytoplasmic parameterization (https://github.com/AllenCell/aics-cytoparam)
- Code used to perform organelle size-scaling analysis (https://github.com/AllenCell/stemcellorganellesizescaling)
- Code used to perform morphing, compute shape modes, and calculate multi-resolution Pearson correlation analysis on 3D single cell images (Rocklin, 2015) (https://github.com/AllenCell/cvapipe_analysis)
- Code to create 12X colony dataset and to perform cell height regression (https://github.com/AllenCell/colony-processing)
- Software will be shared under the Allen Institute Software License and Contribution Agreement, subject to any applicable third-party licensing restrictions.
- Datasets will be shared under the Allen Institute Terms of Use: https://alleninstitute.org/legal/terms-use/.

### METHOD DETAILS

#### Cell lines and cell culturing

The gene-edited cell lines used in this study were created using the parental WTC-11 hiPSC line, derived from a healthy, male donor (Kreitzer et al., 2013). Each gene-edited cell line harbors a fluorescent protein endogenously tagged to a protein representing a distinct cellular structure (Table 1). The complete list of cell lines can be found in the Resource Table. The CRISPR/Cas9-mediated genome editing methodology used to generate these cell lines was previously described in (Roberts et al., 2017b). The tagging strategy for AAVS1 safe harbor targeting was altered to additionally introduce a strong exogenous promoter for expression of CAAX-mTagRFP-T as described previously (Hockemeyer et al., 2009; Oceguera-Yanez et al., 2016). The identity of the unedited parental line was confirmed with short tandem repeat (STR) profiling testing (29 allelic polymorphisms across 15 STR loci compared to donor fibroblasts (https://www.coriell.org/1/AllenCellCollection). Since WTC-11 is the only cell line used by the Allen Institute for Cell Science, edited WTC-11 cells were not re-tested because they did not come into contact with any other cell lines.

The culture and handling protocols for all used hiPSC lines was internally approved by an oversight committee and all procedures performed in accordance with the National Institutes of Health, National Academy of Sciences, and Internal Society for Stem Cell Research guidelines. All cell lines were expanded and grown on an automated cell culture platform developed on a Hamilton Microlab STAR Liquid Handling System (Hamilton Company). This platform is summarized in part in Figure 1A. Three daily workflows were performed on this platform, (1) plate maintenance, (2) passaging, and (3) Matrigel coating of plates. Plate maintenance included the replacement of old media with fresh media for both 6- and 96-well plates. Cells were cultured in a Cytomat 24 (Thermo Fischer Scientific) at 37°C and 5% CO2 in mTeSR1 medium with and without phenol red (STEMCELL Technologies), supplemented with 1% penicillin-streptomycin (Thermo Fischer Scientific).

Cells were passaged every 4 days for up to 10 passages post-thaw. Cells were dissociated into a single cell suspension with 37°C StemPro Accutase cell dissociation reagent (Thermo Fisher Scientific) and counted with a Vi-CELL XR Series cell viability analyzer and associated Vi-CELL XR sample vials (Beckman CoulterA). Cells were re-plated in mTeSR1 medium supplemented with 1% penicillin-streptomycin (Thermo Fischer Scientific) and 10 mM Rho-associated protein kinase (ROCK) inhibitor (Stemolecule Y-27632, STEMCELL Technologies) for 24 hr. Cell culture plates used for cell expansion were clear-skirt, sterile, plastic 6-well plates with lid with condensation rings (Greiner Bio-One). For imaging, samples were plated on glass-bottom, black-skirt, 96-well plates with #1.5 optical grade cover glass (Cellvis). Cells were seeded at a density of 1.3×103 to 3.0×103 in 96-well plates and at 80×103 to 175×103 in 6-well plates.

For most of the dataset, cell culture plates were coated with growth factor reduced (GFR) Matrigel basement membrane matrix, phenol red-free (Lot # 5292003, Corning) diluted with Dulbeco’s modified eagle medium (DMEM)/F-12 (Thermo Fischer Scientific) for a final protein concentration of 0.337 mg/mL. Matrigel coating was performed at 4°C with 100 μl and 1,500 μl added to each 96-well and 6-well, respectively. Plates were incubated at room temperature (RT) for 2hr and Matrigel removed before cell seeding. For the last two cell lines (cohesins and nuclear speckles) imaged on the pipeline, the Matrigel coating protocol was adjusted for improved cell plating. These cells were also plated on a new lot of Matrigel basement membrane matrix, phenol red-free (Lot # 9021357) at a final protein concentration of 0.185 mg/mL. For these samples, glass bottom 96-well plates coated with Matrigel were incubated overnight at 4°C and for an additional 2 hr at 37°C before removing Matrigel at RT. Further details for cell culture reagents and consumables can be found in the Resource Table and standard protocols can be found at www.allencell.org.

#### Cell culture and imaging sample quality control

Rigorous and standardized quality control (QC) workflows for cell culture health were performed at each passage, before imaging the cells at high resolution, and following the completion of imaging a cell line. These QC workflows included cell and morphology assessment via microscopy, cell stemness marker expression with flow cytometry, and outsourced cytogenetic analysis.

hiPSC morphology was evaluated by expert scientists for both plastic 6-well and glass-bottom 96-well plates and was examined 4 days post-passaging. Individual wells of plastic 6-well plates were deemed ready for passage when cells reached ∼85% confluency and displayed typical morphologies associated with hiPSCs that have preserved the expression of stemness markers (Roberts et al., 2017a). Exclusion criteria included, but were not limited to, under- or over-confluency, presence of morphology associated with differentiating cells, and over 5% of cell death. The morphology of cells and colonies grown on glass bottom 96-well plates was also examined prior to 3D field of view (FOV) image acquisition. 12X well overview images were used to exclude wells that did not meet the following four morphology criteria requirements: less than three occurrences of 1) colonies presenting atypical crater-like morphology, 2) lifted colonies (ball-like morphology), 3) partially lifted colonies (edges lifting) and 4) morphology associated with differentiation (Roberts et al., 2017a).

Following the completion of all 3D FOV image acquisition for a given cell line, two types of QC were performed to ensure hiPSCs had retained stemness marker expression and normal G-band karyotyping throughout the imaging period as previously described (Coston et al., 2020; Roberts et al., 2017a). All cell lines imaged during the three years of data acquisition and included in the hiPSC Single-Cell Image Dataset passed theses QC requirements.

#### Image acquisition

The following methods are described in the order they were performed for a given image acquisition workflow on the imaging pipeline. The image acquisition workflow and experimental setup evolved over the three years of dataset collection and was versioned as such. Below is the list of all pipeline image acquisition workflows and a description of each update and modification. The list of pipeline workflow versions used to acquire each cell lines can be found in Table 1.

| Workflow version | Description of changes and updates |
| --- | --- |
| Pipeline 4.0 | Single camera system for interwoven acquisition of four channels. Original transmitted light = white light LED. Specimen exposed to dual peak emissions with highest at 460 nm and range of 400-700 nm. Collecting light with 525/50 bandpass filter, from 500-550 nm. Mode C acquisition performed without photoprotective cocktail. |
| Pipeline 4.1 | Single camera system for interwoven acquisition of four channels. Same original transmitted light source as 4.0. Added photoprotective cocktail with mode C acquisition. |
| Pipeline 4.2 | Single camera system for interwoven acquisition of four channels. Same original transmitted light source as 4.0. Photoprotective cocktail used with mode C acquisition. New emission filter added for acquisition of mTagRFP-T with a 600/50 nm band pass filter. |
| Pipeline 4.4 | Second camera added to all systems. Using dual camera system to image four channels (bright-405-488-638 nm) with interwoven acquisition of 2X two channels simultaneously. New Transmitted Light: Red 740 nm LED Transmitted Light. Specimen exposed to peak emission 740 with narrow range. Collecting light with 706/95 bandpass filter, so from 660-750 nm. Using piezo z stage for fast movement in z. |

##### Microscopy

Imaging was performed on Zeiss spinning-disk confocal microscopes with 10X/0.45 NA Plan-Apochromat or 100X/1.25 W C-Apochromat Korr UV Vis IR objectives (Zeiss) and Zen 2.3 software (blue edition; Zeiss) unless otherwise specified. The spinning-disk confocal microscopes were equipped with a 1.2X tube lens adapter for a final magnification of 12X or 120X, respectively, a CSU-X1 spinning-disk scan head (Yokogawa) and two Orca Flash 4.0 cameras (Hamamatsu). Standard laser lines were used at the following laser powers measured with the 10X the objective; 405 nm at 0.28 mW, 488 nm at 2.3 mW, 561 nm at 2.4 mW and 640 nm at 2.4 mW unless otherwise specified. The following Band Pass (BP) filter sets (Chroma) were used to collect emission from the specified fluorophore; 450/50 nm for detection of DNA dye, 525/50 nm for detection of mEGFP tag, 600/50 nm for detection of mTagRFP-T tag and 706/95 nm for detection of cell membrane dye. Images were acquired with 200 ms exposure time unless otherwise specified. The microscope setup allowed us to collect either all channels with a single camera (Pipeline 4.0-4.2, see description above) or two channels simultaneously, either the bright field and mEGFP or the cell membrane and DNA dyes (Pipeline 4.4, see description above). Cells were imaged in phenol red-free mTeSR1 media on the stage of microscopes outfitted with a humidified environmental chamber to maintain cells at 37°C with 5% CO2 during imaging. Transmitted light (bright field) images were acquired using a white LED light source with broad emission spectrum (pipeline 4.0-4.2) or a red LED light source with peak emission of 740 nm with narrow range and a BP filter 706/95 nm for bright field light collection (Pipeline 4.4 only). A Prior NanoScan Z 100 mm piezo z stage (Zeiss) was used for fast acquisition in z (Pipeline 4.4 only).

##### Well overview and manual well position selection

Typical imaging sessions started with a bright field overview image acquisition of wells from selected rows of a 96-well plate as 2D, 12X tiled images before cell membrane and DNA dye staining. These well overview images were used for final evaluation of cell morphology (see above) and manual or automated position selection for 3D FOV acquisition at 120X in wells satisfying QC criteria requirements (see description above). Manual selection of positions to be imaged at 120X was performed using the 12X overview images and stage function in Zen software. Manual position adjustments were also made at 120X using streaming bright field imaging to satisfy the requirement of each mode of imaging.

##### Imaging modes

Colony position selection was performed manually using the stage function in Zen software or as described below using an automated method (for the last six cell lines imaged) for imaging mode A. One position per colony was selected approximately half-way between the colony edge and center such that the imaged FOV did not fall at the edge nor at the center of the colony. In mode B, positions were also selected as per mode A followed by manual adjustment of the FOV position using transmitted light and streaming bright field imaging to navigate to a region enriched in mitotic cells. This mode was used to substantially increase the number of mitotic cells imaged in an FOV by 3-fold. Operators were trained on how to identify mitotic morphology from just bright field images using merged DNA dye and bright field images (Figure 1A, mode B). In mode C, three positions per colony were selected; a mid-center position area (as in mode A), a position right at the colony edge and a position just inward from the edge in an area referred as the ridge due to the tendency of these cells to grow taller until they flatten into the center area of the colony (Figure 1A; **Figure S7**). Due to the increased photosensitivity of the cells located at the edge of the colony, a photoprotective cocktail (see “Dye Staining” section below) was used when imaging in mode C to prevent premature cell retraction, blebbing and death. Mode C positions were selected manually for all cell lines.

##### Automated well position selection

We developed an automated method that segments the colonies from a 12X well overview image and automatically suggests positions for 3D FOV acquisition based on the distance from the edge of a colony satisfying mode A criteria (see description above). Tiles from the well overview images were acquired with a 10% overlap and stitched using a processing function in Zen software. The automated position selection method segmented the colonies in the image with the following image processing steps developed in Python; 1) rescale intensity to increase contrast of colony edges from background, 2) apply Sobel filter (Scikit-Image) to identify colony edges and fill the holes to segment entire colony, 3) correct for segmentation artifacts with erosion, dilation on segmentation and removal of small objects, 4) identify centers of individual colonies with a distance map on binary segmentation, 5) separate connecting colonies using the center coordinates of colonies and binary segmentation of colony areas with watershed method, and finally 6) compute a distance map for each colony. Our criteria for position selection were as follows, 1) one position was selected per colony, 2) no more than 10 positions were selected per well, 3) the position had to be from a colony greater than 34992 μm2 (corresponding to colony size with a uniform flattened and well-packed central area), 4) the position had to be imaged approximately half-way between the edge and center of the colony (mid-center). To fulfill these criteria, the algorithm first filtered colonies based on their size, and selected mid-center colony positions (x-y coordinates). If more than 10 positions per well were automatically identified, the method gave preference to positions selected in larger colonies and in colonies closer to the center of the well. A graphical user interface was developed to assist users in viewing and confirming the proposed algorithm-generated positions for 3D FOV imaging. The user had the flexibility of moving, adding or deleting positions to finalize the list of FOV to be imaged at higher magnification for that imaging session. The list of positions was then saved as a text file with the stage coordinates and position number in the Zen software readable format (.czsh) and integrated into the experiment xml file for 3D FOV imaging at 120X.

##### DNA and cell membrane dye staining

Following well overview acquisition, the cell membrane and DNA of cells from selected wells were stained with fluorescent dyes. Wells were first incubated at 37°C and 5% CO2 for 20 min with a DNA dye, NucBlue Live (Thermo Fisher Scientific, 1:16.66) diluted in phenol red-free mTeSR1 medium. A cell membrane dye, CellMask Deep Red (CMDR, Thermo Fisher Scientific) was then added to the well (in the continued presence of NucBlue Live) at a final concentration of 5X (earlier lot) or 3X (last 2 lots, adjusted to provide equivalent contrast to noise ratio within a 2.5 hr imaging session) and the 96-well plate was incubated for an additional 10 min at 37°C and 5% CO2. Each well was washed once with phenol red-free mTeSR1 medium before a final 200 μl of phenol red-free mTeSR1 medium was added per well and the plate returned to the stage of the microscope. In mode C acquisition, a photoprotective cocktail (1mM ascorbic acid, 0.3 U/ml OxyFluor and 10 mM lactate) was mixed into the phenol red-free mTeSR1 media before it was added to the well. For consistency, we limited the cell staining to a single row, or 10 wells, per plate at a time and imaged for a maximum of 2.5 hr post completion of the staining protocol. We limited the imaging time to 2.5 hours since we saw no adverse effects of the dyes on cell cycle (evaluated as % mitotic cells in cell colonies) or cell viability (evaluated as increased presence of dead cells on top of colonies) within that time frame. We imaged halfway between the edge and center of a colony to avoid imaging FOVs with reduced dye penetration at the center of large, tightly packed colonies.

##### 3D FOV image acquisition

After the final wash with phenol red-free mTeSR1 media, plates were returned to the stage of the spinning-disk confocal microscope and all 3D FOVs, at pre-selected positions, were acquired with a 100X/1.25 NA objective at a final magnification of 120X. Four channels were acquired at each z-step (interwoven channels) in the following order: bright field, mEGFP or mTagRFP-T, CMDR and NucBlue Live with laser powers and exposure times as stated above. For the single camera system acquisition only, empty channels were acquired between each channel with 0.3 ms exposure time to reduce noise introduced during the filter position change. This was necessary due to the long travel range of the filter wheel moving between four different positions at each z-step. Pipeline 4.4 3D FOV acquisition was performed with two cameras using two interwoven sets of simultaneous acquisitions. In this case, bright field and CMDR channels were acquired on the back camera and all other channels acquired on the left camera. Resulting images from either single or dual camera systems were of 16 bits and 924 x 624 pixel2 in x-y dimension after 2×2 binning. Images from the dual-camera system required channel alignment (see below). Following channel alignment, the final images were cropped into a final size of 900 x 600 pixels2 in the x-y dimension. FOVs from both single and dual camera systems had a final x-y pixel size of 0.108 µm and z-stacks composed of 50–75 z-slices (to encompass the full height of the cells within an FOV) acquired at a z interval of 0.29 µm.

#### Post-acquisition FOV image processing

##### Channel alignment for dual camera acquired images (Pipeline 4.4 only)

Optical control images were acquired at the start of each data acquisition day to monitor microscope performance. Optical control images of TetraSpeck microsphere beads or the “field of ring” pattern on the Argolight HM slide (Argolight) were used to register and align the appropriate channel images of an FOV acquired with two cameras. A z-stack of 10 to 30 z-slices of these patterns was acquired at 120X with all four fluorescent channels. Channel images from the 638 nm laser line and 706/95 nm BP filter (back camera) and 488 nm laser line and 525/50 nm BP filter (left camera) were used to generate an affine transformation matrix identifying the shift in x-y, rotation and scaling factor between the 638 nm (from the back camera) and 488 nm (from the left camera) wavelength channels. We used the z-slice with maximum focus along the z-axis. The two channel images were pre-processed separately by normalizing the intensities and applying gaussian smoothing prior to segmenting the objects such as individual beads or rings with intensity thresholding. Due to the nature of the sample preparation of TetraSpeck beads, which randomly adhere to the glass, we excluded some beads based on the following criteria: 1) overlapping beads, 2) beads that are outside of the range of an expected bead size and intensity, and 3) beads that have inconsistent centroid location (mass versus peak intensity). Centroid locations of segmented objects (beads or rings) from both 638 nm and 488 nm channels were compared and only objects in close proximity (within 5 pixels) between the two channels were kept. The exclusion steps were not necessary with the stable and consistent field of ring pattern of the Argolight HM slide. Using the two sets of centroid locations of objects, the method estimated a similarity transform matrix with the “estimate_transform” function in scikit-image that transforms the image with translation, rotation and scaling. The values from this matrix were also used to identify any deviations from the normal trend indicating potential system performance issues over time. The affine transformation matrix was applied on every z-slice of the channel acquired on the back camera (bright field and 638 nm) and as such aligned to the reference channel images acquired on the left camera (405 nm, 488 nm and 561 nm) with a Warp function (scikit-image). FOVs were then cropped in x-y for a final dimension of 900 by 600 pixels2 to remove empty pixels introduced in the bright field and 638nm channel images by the alignment.

##### 3D FOV image quality control

All 3D FOV images were visually inspected by experts for obvious issues related to the experimental settings. Typical exclusion criteria were related to microscope acquisition system failures (laser, exposure time, z-slice positioning in relation to cell height, empty or out of order channels), or any other issues that would cause downstream processing to fail or analysis steps to identify outliers. Some of these QC steps were also automated with a series of Python scripts to ensure a more systematic and standardized way to catch problematic FOVs and exclude any outliers. To do so, intensity metrics were extracted from each channel of each FOV and trends and averages were used to determine exclusion thresholds or cutoff values. Overall, three main automated exclusion FOV QC steps were applied to the hiPSC Single-Cell Image Dataset; channel intensity out of range, z-stacks with incomplete cell height, and z-slice empty or out of order.

###### FOV channel intensity quality control

The FOV channel intensity QC script calculated the minimum, maximum, median, 99.5th and 0.5th percentile pixel intensity value in each channel for each FOV. FOVs were flagged if the median intensity of one channel was outside a predetermined range (low and high cutoffs, see values below). These cutoff values were based on offset, noise and maximum intensity values of the microscopes, fluorescent tags and dyes imaged.

###### Cutoffs for median intensity

| Channels | Low cutoff<br>(AU) | High cutoff<br>(AU) |
| --- | --- | --- |
| bright<br>field | 0 | 50000 |
| 405 nm | 400 | 430 |
| 488 nm | 400 | 1600 |
| 561 nm | 400 | 700 |
| 638 nm | 400 | 8000 |

Low cutoffs for the maximum intensity in the 405 nm (DNA) and 638 nm (cell membrane) channels were also applied to ensure the minimum required contrast in the images for successful single cell segmentation.

###### Cutoffs for max intensity

| Channels | Low cutoff<br>(AU) |
| --- | --- |
| 405 nm | 500 |
| 638 nm | 635 |

Given a normal distribution of FOV intensities, we also excluded from the dataset all individual FOVs with a channel median intensity within the bottom 0.5th percentile of the whole dataset. We calculated a z-score for each channel of each FOV and excluded all FOVs that had a channel intensity with z-score of 2.58 below the mean.

###### Automated detection of z-stack with incomplete cell height quality control

We automated the detection of z-stacks with incomplete cell height in a FOV due to mis-positioning or mis-sampling of a z-stack acquisition. The cell membrane channel (638 nm) was used to determine whether the top and/or bottom of the cell were included in the z-stack image. The intensity of the cell membrane channel image was first normalized to the maximum intensity of the cell membrane channel image. Next, the median intensity and contrast (maximum intensity-background intensity)/maximum intensity) for each z-slice were calculated to generate an intensity and contrast profile along the z-axis. Local maxima of the intensity profile were detected with a “peak detection” method described in SciPy-image ((Virtanen, 2020); scipy.org), where the lower peak corresponds to the bottom of a z-stack and the higher peak corresponds to the top of a z-stack. In the scenario where more than 2 peaks were detected, the method used the top-most peak and the bottom-most peak and the contrast profile to refine the measured range of the z-stack.

Thresholds of contrast values for bottom (0.2) and top (0.19) of a z-stack were estimated from data trends of the entire dataset. Using these threshold values, the method iterated from the top/bottom peaks detected to the full range of the z-stack and reported the closest z-slice to reach the thresholds as the detected top/bottom of the cells for this z-stack. We also measured the rate of change in contrast in the detected top and bottom 5 slices of each z-stack and flagged z-stacks as incomplete cell height if the rate of change in the top 5 slices were smaller than −0.015 and the bottom 5 slices were smaller than −0.01 (in contrast units, see contrast definition above). To ensure cell height completeness any FOV with detected top/bottom z-slices within 5 slices of the first and last slice of the z-stack were flagged as either an “incomplete -missing top” or a “incomplete -missing bottom” and excluded these FOVs from the datasets.

###### Out of order z-stacks in FOV quality control

Out of order z-stacks were also observed. We generated an algorithm capable of identifying if the z-stack first z-slice had the highest median intensity, indicating that the z-stacks were placed in improper order by the acquisition software. We excluded any FOV with a first z-slice registering the maximum intensity and flagged the FOV as “z-stacks out of order”.

#### 3D segmentation

##### Cell and nuclear segmentation

To segment each individual cell and its corresponding DNA from the membrane dye and DNA dye channels of each 3D z-stack, we used the deep learning-based cell and nuclear instance segmentation algorithm developed as part of Allen Cell and Structure Segmenter (Chen et al., 2018). We combined the Segmenter’s Iterative Deep Learning workflow and the Training Assay approach to ensure accurate and robust segmentation for downstream quantitative analysis. Complete step-by-step details of this algorithm are described in (Chen et al., 2018).The code, trained models, and demo Jupyter notebooks have been released at https://github.com/AllenCell/segmenter_model_zoo. In the Training Assay approach, a secondary experimental assay that is more amenable to accurate segmentation is linked to the primary assay for the purpose of training segmentation models. The secondary assay is used to generate accurate segmentations, which are then imposed as the target for training the model to segment the images of the primary assay. As a result, the final segmentation model can achieve better accuracy and robustness even when running on the poorer-quality primary assay images. We applied two training assays to develop the cell and nuclear segmentation algorithm.

The first training assay (**Figure S1E**) addressed the challenge that the membrane dye images suffered from very weak signal near the top of cells due to both dye labeling of a very thin membrane and photobleaching even during a single z-stack acquisition via 3D spinning-disk confocal microscopy. The secondary assay in this training assay used the CAAX cell line containing the membrane-targeting domain of K-Ras tagged with mTagRFP-T, which made it possible to accurately delineate cell boundaries, even near the top of cells. This training assay is described in detail in (Chen et al., 2018). In brief, the first step of this training assay is to obtain the initial semantic (whole FOV) segmentation of tagged CAAX signal on ten sample images using a semi-automatic algorithm based on a seeded watershed. Seven images were sorted as having good segmentation and used to train a CAAX segmentation model. We then applied this CAAX segmentation model on 312 CAAX images to create a CAAX-based cell segmentation ground truth set, which we then used together with the membrane dye images to train a membrane dye-based segmentation model. This model robustly segmented cells including their dimly visible top boundaries, from the membrane dye images in all 18,186 FOV’s in the dataset.

The second training assay (**Figure S1F**) was to use images of mEGFP-tagged lamin B1 cells for segmenting interphase nuclei and mEGFP-tagged H2B cells for segmenting mitotic DNA during mitosis (representing the “nucleus” during nuclear envelope break-down). Lamin B1 and H2B both provided more biologically accurate detection of the nuclear boundary. The shell of intensity around the nucleus in tagged lamin B1 cells was more directly detectable in 3D than the DNA dye images and both endogenously tagged structure cell lines had better signal to noise compared to both dyes. This training assay is described in detail in (Chen et al., 2018). Briefly, we began with classic image segmentation results for lamin B1 where the “shell” of lamin B1 is filled to represent the nucleus. We sorted eight out of 80 images and used these to train a deep learning model to segment “nuclei” (i.e. filled shells) from lamin B1 images. We then applied this model on 1,017 lamin B1 images to create a lamin B1-based nuclear segmentation ground truth set, which we then used together with the DNA dye images to train a DNA dye-based segmentation model for interphase nuclei. Regions containing mitotic cells in these images were automatically identified and excluded from training, see (Chen et al., 2018) for details. In parallel we used H2B images and a classic segmentation workflow to generate a cleaner segmentation target for training a mitotic DNA segmentation model. We generated a set of 28 merged segmentation targets (mitotic DNA segmentation from H2B images and interphase nuclei segmentation by applying the first interphase model on DNA dye images) to train an overall DNA dye-based nuclear segmentation model. This is the model we applied to all 18,186 FOVs in the dataset.

To convert the cell and nuclear segmentation model outputs into individual cells (i.e., instance segmentation), we had to train two additional models: a “cell seeding” model and a “mitotic pair detection” model. We took advantage of the DNA dye-based nuclear segmentation model to create a deep learning based “cell seeding” model. This used a subset of 600 images from the same training images as for the interphase DNA dye segmentation model, but with a modified segmentation target obtained by shrinking the mask for interphase nuclei (and very early and very late mitosis DNA), and generating a convex hull for the mask for other mitotic DNA. The binarized membrane segmentation model output was used to cut the potentially falsely merged seeds from tightly touching nuclei and the resultant seeds were applied back on the cell membrane segmentation output for use in a seeded watershed to identify individual cells. We also trained a FasterRCNN-based mitotic pair detection model, which permitted us to identify mitotic cells that were in anaphase and telophase/cytokinesis and make sure they were segmented as one cell. Several other steps were performed to enhance the robustness of the cell and nuclear segmentation for application at scale to the 18,186 FOVs in the hiPSC Single-Cell Image Dataset. These are described in detail in (Chen et al., 2018) and included training and applying a label-free segmentation model of nuclei and cell membrane to boost the robustness when the signals in the DNA dye or membrane dye channel were extremely dim, as well as several minor steps such as morphological refinement on the segmented nuclei and refinement of the bottom of the cell. The very bottom surface of the cells protrudes out into the tightly packed neighboring cells and the z-resolution does not permit proper disentangling of the overlapping parts. We therefore automatically identified a z-slice with a reasonable cell segmentation near the bottom and propagated it downward through all other z-slices to the bottom of the cell.

To validate the performance of the cell and nuclear segmentation results, we selected and inspected a representative set of images (576 images from 22 different cell lines) at the single-cell instance level. From this validation, we estimated the percentage of well-segmented cells and the percentage of FOVs for which the segmentation of all cells and nuclei were successful without obvious errors along the segmented boundaries. We developed an in-house scoring interface in Python using Napari that allows for overlaying the segmentations on the original images and inspecting them slice by slice in 3D. Each image was manually scored by at least two human experts. We found that over 98% of individual cells were well-segmented and over 80% of images generated successful cell and nuclear segmentations for all cells in the entire FOV. Based on these validation results, we decided the cell and nuclear instance segmentation algorithm was sufficiently reliable to be applied to all of the FOVs in the dataset. For quality control purposes, single-cell visualizations of all segmented cells were generated using https://github.com/AllenCellModeling/actk as a set of contact sheets and all cells in the final dataset were manually reviewed for basic quality criteria such as only one nucleus per cell except later in mitosis, no obviously chopped nuclei, and no especially aberrant cell shapes due to segmentation errors.

##### Cellular structure segmentation

We applied a collection of modular segmentation workflows from the Classic Segmentation component of the Segmenter, each optimized for the particular morphological features of the target cellular structures (Chen et al., 2018). Representative examples for each of the 25 FP-tagged cellular structures are shown in **Figure S1**. For each structure, results of the segmentation workflow were evaluated on sets of images representing the variation observed across imaged cells (e.g. different regions of colonies) to ensure consistent segmentation quality across all images for each structure. The algorithms for all but two of the 25 cellular structures were classic image segmentation workflows. The exceptions were the plasma membrane (via CAAX) and the nuclear envelope (via lamin B1). All Classic Segmentation workflows contain three parts: pre-processing, core segmentation algorithms, and post-processing. For each part, there exists a set of algorithms to choose from, each with restricted numbers of parameters to tune. All workflows are accessible at https://github.com/AllenCell/aics-segmentation. For the plasma membrane, a deep learning-based segmentation model was developed as part of the training assay for cell segmentation described above. For the nuclear envelope, we developed an algorithm that combines multiple deep learning models including (1) the lamin B1 filled segmentation model we developed for nuclear segmentation training assay, (2) an overall lamin B1 segmentation model, (3) a lamin B1 seeding model, and (4) the plasma membrane segmentation model developed for cell and nuclear segmentation model. Briefly, we used the plasma membrane segmentation model output to cut the lamin B1 seeding model outputs to generate one seed per interphase nucleus. Then, the seeds were applied on the overall lamin B1 segmentation model via seeded watershed to obtain a one-voxel thick “shell” for each interphase nucleus. The “shells” were merged with the overall lamin B1 segmentation as the final lamin B1 segmentation result, which contained both complete nuclear envelope and properly segment invaginations and lamin B1 during mitosis. More details can be found in (Chen et al., 2018). Both models and code for CAAX and lamin B1 can be accessed via https://github.com/AllenCell/segmenter_model_zoo.

We performed an additional validation step to determine whether a given target structure segmentation was sufficient for interpretation in the cellular structure volume analysis (Figure 6). We identified ten structures for which there were obvious caveats to the ability to use their target structure segmentation for biological interpretations of how much of the target structure was present in each cell and thus these ten structures were excluded from the structure volume analysis (**Figure S1B-D**). These three types of caveats included: (1) The cell boundary segmentation may have potential segmentation errors in the very top slices of the cell. This type of error has a minor effect on the overall segmentation of the cell but for structures localizing to the cell periphery at the very top of cells, this caveat can cause structures to be miss-assigned to neighboring cells (including tight junctions (ZO-1), gap junctions (connexin-43), desmosomes (desmoplakin; **Figure S1B**), adherens junctions (beta-catenin), actin filaments (beta-actin), actin bundles (alpha-actinin-1), and actomyosin bundles (non-muscle myosin IIB)). Therefore, these seven structures were not validated for cellular structure volume analyses. (2) Structures localizing or partially localizing to a thin 3D surface (such as the cell or nuclear periphery), especially when that surface is slanted, may suffer from non-uniform accuracy between the middle and the top/bottom of that structure due to the anisotropic resolution of the images. The accuracy of the nuclear pores target segmentation was sufficient to identify the general location of nuclear pores in the cell for the location-based analyses but not sufficient to be validated for use in the cellular structure volume analysis and thus this structure was excluded (**Figure S1C**). This nuclear periphery caveat was also observed for perinuclear ER (both Sec61 beta and SERCA2) and the nuclear lamina enriched localization of histones (H2B). However, these structures were still well segmented for the cytoplasmic ER localized throughout the cell and for histones localized throughout the nucleoplasm, each of which contributed more to overall structure volume. Therefore, those structures were not excluded from the cellular structure volume analysis. This caveat was also observed for structures that localize to the cell periphery (listed in the first caveat), which were excluded from the structure volume analysis. (3) The segmentation result for cohesins (via SMC1A) can depend on how far along a cell is in interphase and works well for most, but not all, of interphase (**Figure S1D**). Therefore, this structure was excluded from the structure volume analysis. Matrix adhesions (paxillin) localized to the very bottom of the cells where the membrane dye signal does not permit accurate identification of cell boundaries (see “Cell and nuclear segmentation” in Methods). Therefore, due to high likelihood of misassigment of matrix adhesions to neighboring cells, they were excluded from the structure volume analysis.

#### Single cell dataset generation

##### Single cell image generation

To build the single-cell version of the image dataset for downstream analysis, we extracted all complete individual cells in each FOV automatically from the cell segmentation results of the image, ignoring any cells that were not at least 4 pixels away from the image border in the xy-plane (∼12 complete cells per FOV, on average). All images were rescaled to isotropic voxel sizes by interpolating along the z dimension to upscale the voxel size from 0.108333 µm x 0.108333 µm x 0.29 µm to 0.108333 µm x 0.108333 µm x 0.10833 µm. For each cell, a cropping region of interest (ROI) was calculated by extending the 3D bounding box of the cell by 40 voxels in each direction in both x and y and by 10 voxels in each direction in z.

This same cropping ROI was applied to the original intensity z-stacks to extract the DNA, membrane and tagged structure for each cell. Similarly, the cropping ROI was used to extract the cell, nuclear and structure segmentations for each cell within this ROI. These extracted segmentations were then each masked by the cell segmentation result such that all voxels outside of the segmented boundary of the cell was set to zero. A roof-augmented version of the cell segmentation was also calculated for each cell to ensure proper inclusion of structures within the cell due to limited resolution and accuracy near the top of the cells (see “Single cell basic feature extraction” section). The roof-augmented cell segmentation is created by applying a morphological dilation (voxels only along the z-axis) at the top 25% of the cell segmentation mask. Each individual cell is thus associated with five segmentations: DNA segmentation, cell segmentation, roof-augmented cell segmentation, structure segmentation, and roof-augmented structure segmentation, which is masked by the roof-augmented cell segmentation after ROI cropping.

##### FOV-based feature extraction

FOV-based features were calculated for each cell. Specifically, we calculated (1) the Euclidean distance from the nucleus of each cell to the nucleus of each complete neighboring cell within the FOV, (2) the lowest and highest z position of all cells in this FOV, and (3) whether a cell is located on edge of a colony, for those cells within colony edge FOVs (mode C edge; see image acquisition methods). All details are released via https://github.com/AllenCell/cvapipe.

##### Colony-based feature extraction

In addition to FOV-based and single-cell-based features, we extracted colony-based features. For each 12X overview image and 120X FOV image taken, we extracted the name of the well in the 96-well plate and the stage coordinates at which the image was taken from the file metadata. To obtain colony segmentations from the 12X overview images, we applied the same segmentation method used for automated position selection (see “Automated well position selection” section above). We then associated each segmented colony with a set of colony or well features including the confluency of the well, the size of the colony, the centroid location of the colony in the overview image, and whether the colony was touching the boundary of the overview image. We mapped the position where the 120X FOV image was taken relative to the 12X overview image by using the microscope stage coordinates, identified the colony in which that 120X FOV image was taken and added colony features to this 120X FOV image. We also calculated the Euclidean distance between the center of the FOV image and the nearest edge location of the colony. We added QC methods to ensure data accuracy and usability by flagging 120X FOVs with: (1) poor colony segmentations, detected as well confluency less than 10%, (2) 120X FOV images that were taken outside of the 12X overview image FOV and (3) 120X FOV images that were in a colony touching the edge of the 12X overview image. These colony-based features were not only linked to each 120X FOV but also to all of the individual cells associated with that FOV.

##### Deep learning based single cell annotation

Each cell in the hiPSC Single-Cell Image dataset was automatically annotated by a deep learning-based classifier into one of the following 7 annotation categories: interphase, prophase, early prometaphase, prometaphase/metaphase, anaphase/telophase (unpaired cell), anaphase/telophase (paired cell) or other (e.g. failed segmentations, dead cell segments, or dye blobs). Note: unpaired cells in anaphase/telophase refer to cells where it was impossible to find the other member of the pair (e.g. the other pair member is outside of the FOV). The automated classifier is a combination of a rule-based classifier and an ensemble of three 9-class 3D ResNet50 models. First, a cell is annotated as category anaphase/telophase (pair) if the nuclear segmentation satisfies the following three criteria: (1) contains at least two connected components, (2) the ratio of the sizes of the largest two connected components is greater than 0.64, an empirically determined value, (3) the distance between the centroid of the largest two connected components is greater than 85 voxels. Otherwise, the 9-class ResNet50 models are used. To train the ResNet50 models, we created a training set consisting of 5,664 cells from the main dataset and through expert-annotation assigned these into 9 classes: 1-interphase, 2-prophase, 3-early prometaphase, 4-prometaphase/metaphase, 5-anaphase/telophase unpaired, 6-anaphase/telophase paired (but not necessarily satisfying all three criteria), 7-failed segmentation, 8-dead cell segments and 9-dye blobs. Class 1 (interphase) accounted for 43.5% of the data to ensure a balanced training set, while the total of classes 7, 8, and 9 accounted for 2.9%. Three ResNet50 models were trained with different training/validation splits. An ensemble of these three models was used to make the final class predictions. These ResNet50 models were validated by testing on 100 cells that were held out from the training set. The model generated eight incorrect predictions, but all were either incorrectly predicting mitotic stages (3/100) or incorrectly predicting a cell in interphase to be in mitosis (5/100). The recall rate for interphase cells was 100%. Cells that were predicted to be of classes 7, 8 or 9, or that generated prediction of low confidence, were annotated as belonging to the “other” category and removed from the hiPSC Single-Cell Image dataset. The confidence score of a prediction was approximated as the highest probability among all 9 classes. Confidence scores lower than 0.677 were considered low confidence and these cells removed. The final automated 3D image classifier code (for both training and testing) and all trained models are available at https://github.com/AllenCell/image_classifier_3d.

##### Single cell basic feature extraction

Methods described here are implemented in several repositories for different parts of the analysis and begin with the hiPSC Single-Cell Image Dataset (see Data and code availability above). The data table downloaded from Quilt contains 216,062 rows and 47 columns, where each row corresponds to a single cell uniquely identified by its ID that is specified by the column CellId. The columns contain both the necessary information about each cell (e.g., path to segmentations, path to images, and important meta data) to calculate single cell features, as well as the calculated features. These features are included ready to download for ease of use. Demos in https://github.com/AllenCell/cvapipe_figure_notebooks can be used to produce the main figures from these released features, while all these features can be reproduced with the released code (see Data and Code availability above, (McKinney, 2011; Nicholas Sofroniew et al., 2019; Paszke et al., 2019; Pedregosa et al.; Walt et al., 2014)). The cell segmentation, the DNA segmentation and the roof-augmented structure segmentation were used to extract basic cell, nuclear and cellular structure features, respectively. The analysis dataset (see just below) contains only interphase cells, so the DNA segmentation represents the nucleus. The cell or nuclear segmentation for each cell is used as the input for calculating the following basic cell and nuclear features, respectively: (i) cell or nuclear volume as the number of non-zero voxels in the input image. The single-voxel volume (0.108 µm)^3^ was used to rescale this feature for further analysis. (ii) cell or surface area as the number of voxel sides facing the background in the input image. According to this metric, an isolated voxel has all its 6 sides facing the background and therefore a surface area equal to 6. The single-voxel-side area of (0.108 µm)^2^ was used to rescale this feature for further analysis (iii) cell or nuclear height as the distance in voxels along the z-axis between the bottom-most and top-most voxels in the input image. The single-voxel height of 0.108 µm was used to rescale this feature for further analysis. To calculate the volume of each cellular structure within a cell, the roof-augmented structure segmentation of that cell was used as the input. This ensures proper inclusion of structures within the cell due to limited resolution and accuracy near the top of the cells (see “Single cell image generation” section). The volume of the cellular structure in the cell is calculated as the number of non-zero voxels in the input image. We used the single-voxel volume (0.108 µm)^3^ to scale this feature for further analysis. These single cell basic features were merged into the hiPSC Single-Cell Image Dataset as additional columns and used in subsequent quantification and analysis.

### QUANTIFICATION AND STATISTICAL ANALYSIS

#### Analysis dataset generation

##### Mitotic cells removal

The first operation performed on the full dataset to create the analysis dataset was the removal of all of the 11,238 mitotic cells. This is done by removing all rows of the data table for which the column cell_stage is different from M0, the value used to flag interphase cells, resulting in a table with 204,824 rows (cells).

##### Outlier detection

In total, 1,087 (∼0.5%) cells were identified and removed from the dataset, resulting in a table with 203,737 rows that we refer to as the analysis dataset throughout the paper. These outliers fall into two classes. First, there were 670 cells for which the structure volume was zero. Cells with an empty structure segmentation could be real outliers (e.g., no FP signals within that specific cell) or could indicate errors in either structure segmentation or cell and nuclear segmentation (see caveats in the “Structure Segmentation” section). Since cells with zero structure segmentation only account for ∼0.3% of the whole population, we considered all such cells with potential segmentation errors, even minor, in cell and/or nuclear shapes as outliers. Second, we identified 417 cells, which were identified as outliers by an automated bi-variate outlier detection algorithm. Here, “bi-variate” refers to the notion that we looked at pairs of two variables to detect outliers and not at a single variable. As an example, the outlier detection procedure identified cells as outliers that have a very large cell volume (first variable) but very small nuclei (second variable), and clearly fall outside of the typical distribution of cell and nuclear volume. The outlier detection algorithm uses Gaussian kernel density estimation on the 2D space spanned by two variables, thereby assigning a probability to each of the cells. We use density estimation in the same way for visualization of bi-variate associations in scatter plots (see “Visualization of bi-variate association” section below and Figure 6). Cells with an extremely low probability were identified as outliers. We applied this outlier detection to the 21 pairs of variables that can be made of the seven main cellular and nuclear metrics: cell volume (µm^3^), cell surface area (µm^2^), cell height (µm), nuclear volume (µm^3^), nuclear surface area (µm^2^), nuclear height (µm), cytoplasmic volume (µm^3^). Cells with resultant probabilities smaller than 1e-20 were identified as outliers (n=177). This outlier analysis was also applied to pairs of variables for the four following cell and nuclear metrics (cell volume and surface area, nuclear volume and surface area) each with cellular structure volume for the 15 structures validated for structural volume analysis, totaling 15×4=60 scatter plots. Cells with a probability smaller than 1e-10 in any of these 60 scenarios were identified as outliers (n=240). The thresholds mentioned above were identified manually after inspection of the scatter plots and visual inspection of many cells identified as outliers. The majority of the inspected cells clearly showed imaging or segmentation artifacts.

#### Statistical analysis for quality control of the hiPSC Single-Cell Image Dataset

To be able to map cells from the 25 cell lines into the same shape space and cluster similar cells to integrate the location of their separately imaged structures we must first ensure that the cell lines themselves are not an experimental source of cell and nuclear shape variation. Further, this extensive dataset was acquired over a period of three years, including changes in the extent of pipeline automation, necessary adjustments to the microscopes, the lots of Matrigel, and other such experimental factors over the course of the imaging pipeline timeline (see “Imaging workflows” section). Therefore, we performed an extensive analysis to identify and account for any potential experimental contributions to cell shape variation (**Figure S7**). An analysis of how each of the Shape Modes varied with respect to the timeline of the imaging pipeline revealed that only Shape Modes 1 and 2, representative of cell height and cell volume, showed any signs of possible systematic experimental variation (**Figure S7A**). For cell height, we observed variation between cell lines throughout the pipeline timeline, while for cell volume we only observed a possible systematic difference between Pipeline 4.4 and the rest of the pipeline workflows (**Figure S7B&C**). The greatest systematic effect on cell height over the pipeline timeline was visible in the sequential imaging of the last two structures (nuclear speckles via SON and cohesins via SMC1A), which both contained flatter cells. These differences were attributable to a change in both the lot of Matrigel and an adjustment to the glass bottom well-plate Matrigel coating protocol as described above. This can be seen in a control experiment comparing the tagged actomyosin bundles (via non-muscle myosin IIB) cell line before and after this protocol change (**Figure S7B**). We separated the pipeline timeline into three periods, the period before Pipeline 4.4, and then within Pipeline 4.4, the period before and after the change in Matrigel coating protocol and compared both cell height and cell volume between these periods. We found that while the adjusted Matrigel coating protocol decreased cell height significantly, it did not affect cell volume. However, both cell height and cell volume were slightly and consistently decreased during the entire Pipeline 4.4. Further investigation into possible causes revealed a systematic inaccuracy in z spacing due to the use of a piezo z stage, which leads to an approximate 10% reduction in the z-step size and thus also in the overall height of the cell. When we corrected the Pipeline 4.4 z-step size by this approximate amount, we found this could account for the cell height difference. Cell volumes cannot be directly corrected by one single factor adjustment due to the varied cell shapes. However, the slight, yet significant and consistent decrease in average volumes of all cell lines imaged during Pipeline 4.4 can be accounted for by the same piezo-dependent problem. Unfortunately, we could not retroactively determine the exact adjustment to the z-step size for each independent image acquisition that was performed during Pipeline 4.4 and thus did not correct the data for this issue. However, the magnitude of the effect was much smaller than the variation of cell volumes and heights within the cell line datasets.

In addition to these two systematic experimental sources of variation during Pipeline 4.4, we observed variation in average cell height throughout the entire pipeline timeline. This suggested additional possible sources of variation. We had experimentally observed that cell height seemed to vary both with colony area and the location of cells within a colony (Figure 1A), suggesting that cell height variation might be part of normal changes to cell packing behavior within a growing colony. To test this observation quantitatively, we measured the cell area of a subset of colonies with accurate colony segmentations as well as both the distance from the center of the FOV to the edge of the colony and the average height of all the cells within that FOV. We transformed colonies and the locations of FOVs within them into circular representations and compared the location patterns, cell heights, and colony areas (**Figure S7D**). We found that smaller colonies tended to contain taller cells while in larger colonies, cells closer to the colony periphery were taller than those towards the center of colonies. Other than Shape Mode 1, representing cell height, none of the other shape modes showed any colony-specific patterns within the dataset (**Figure S7E**).

We next investigated how much of the variation in cell height (median height of the cells in an FOV) was explained by a set of eleven experimental variables including the distance of an FOV to the colony edge representing the position of cells in a colony, the colony area, the cell line identity, and several imaging pipeline settings (**Figure S7F**). We performed a Random Forest regression analysis (Liaw and Wiener, 2002) and found we could predict cell height with moderate accuracy (R^2^ = 0.52) based on this combination of eleven variables. When we removed cell line identity as a variable within this regression analysis, the accuracy of cell height prediction barely change (R^2^ = 0.51). The feature “FOV to colony edge distance” had the largest feature importance. Importantly, we found that cell line identity was statistically correlated with several imaging pipeline settings that varied throughout the imaging pipeline timeline.. All of the results above together confirm that cell line identity can contribute to cell height variation due to the fact that each cell line was imaged under a particular set of imaging conditions which varied throughout the imaging pipeline timeline, but that cell line identity itself does not greatly contribute to the variation in cell height observed in the hiPSC Single-Cell Image Dataset.

##### Circular colony mapping

We took advantage of the fact that many cells (n=104,269) of our hiPSC Single-Cell Image Dataset could be associated with information relative to the colony where they came from (see “Colony-based feature extraction” section), to visualize radially dependent spatial patterns of our cells. This is achieved by mapping the location of cells in a colony into a unit circle, as illustrated in **Figure S7D**. First, the distance from the center of the FOV to the closest edge point (d) is normalized by the effective radius of the colony (R_eff_) to determine the relative distance ℓ=d/R_eff_. Then, all cells in the FOV are mapped into a unit circle at radial distance ℓ from the edge of the circle. Each cell is assigned to an angular location drawn from a uniform distribution of angles in the range [0,2π].

##### Random forest regression model to predict cell height from experimental features

A multivariate Random Forest regression model was trained to predict the median cell height of all cells in an FOV from experimental, assay-dependent variables, including (1) cell growth information from the confluency of cells in the well, Matrigel-coating protocol, the FP-tagged protein name, and two cell passaging numbers, (2) colony features from the size of the colony the FOV was imaged at and the distance between the FOV and the nearest colony edge, and (3) instrument hardware configurations including the pipeline workflow information, the ID of the microscope which the FOV was taken with and the piezo configuration of the microscope. We first calculated the median cell height of an FOV from the single-cell segmentation that provides the height of each cell in the FOV. We then pre-processed the continuous variables (FOV to colony edge distance, confluency, colony area, total passages and passages post-thaw) with z-normalization, and labeled categorical variables (cell line via its FP-tagged protein name, imaging mode, workflow ID, Matrigel protocol, piezo setting, microscope ID) in R Studio. We added a control variable by randomly generating a number that ranges from −3 to 3 for each FOV.

This analysis was based on the 20 cell lines that contain >100 FOVs each, with a total of 7,914 FOVs. The cell lines with the following tagged proteins (Table 1) were included: alpha-actinin-1, alpha-tubulin, beta-actin, CAAX, centrin-2, connexin-43, desmoplakin, fibrillarin, H2B, lamin B1, LAMP-1, non-muscle myosin IIB, Nup153, paxillin, Sec61 beta, sialyltransferase 1, SMC-1A, SON, Tom20, and ZO-1. For each cell line, we randomly selected 90 FOVs for training, resulting in a training dataset of 90*20 = 1,800 FOVs and used the remainder of the FOVs (n = 6,114) to evaluate the model. We trained a Random Forest model with using all variables in R Studio with the RandomForest package (Liaw and Wiener, 2002) with 500 trees. We also trained another Random Forest model with all variables except cell line identify, again using 500 trees. We evaluated the model by calculating the Coefficient of Determination (R^2^) on the test set (n = 6,114 FOVs). Feature importance scores were calculated as the difference in mean squared error (MSE) between a model including the feature in question and a model where the values of that feature were randomly permuted across the samples. We repeated the sampling and model training 100 times to obtain confidence intervals of model performance and feature importance as shown in **FigureS7F** (left).

#### Spherical harmonics expansion (SHE) of cell and nuclear shapes

In addition to the basic features described above, we also used SHE coefficients as shape descriptors for cell and nuclear shape (Ruan and Murphy, 2019; Shen et al., 2009). We created a publicly available open-source Python package, aics-shparam (see “Data and Code Availability” section) to extract SHE coefficients from segmented images of cells and nuclei.

##### Cell and/or nuclear alignment

SHE coefficients are sensitive to the orientation of the shape they are extracted from. Therefore, a given set of cells and nuclei can be used to create different versions of a shape space, depending on how they are pre-aligned. To create the cell and nuclear joint shape space (Figure 2) we wanted to preserve the apical basal axis of the cell, which is the z-axis in the lab frame of reference. Therefore, we only aligned cells by rotation in the xy-plane. Cells and nuclei were rotated such that the longest cell axis falls along the x-axis. The cell segmentation was used to estimate the longest axis of the cell through a principal component analysis of the x and y coordinates of foreground voxels. The longest axis was defined as the direction of the first principal component and the alignment angle defined as the smallest angle between the longest axis and the x-axis. That cell was then rotated by the alignment angle such that the longest axis was aligned with the x-axis. Cells were rotated by using the function rotate from Python package scikit-image (Walt et al., 2014) with zero order interpolation. The input image was also resized as necessary to fit the whole rotated cell. The alignment procedure was implemented by the function align_image_2d in aics-shparam using default parameters. This function returns the final alignment angle, which is then used to align other images related to that cell, in this case the segmented images of the nucleus and the particular cellular structure in the cell as well as the three channels of the z-stack containing the original images of the membrane dye, DNA dye and FP-tagged structure. This was done using the function apply_image_alignment_2d available in the same Python package.

##### From segmented, aligned images to SHE coefficients and 3D meshes

Once a segmented image of a cell and nucleus is aligned, it is used as input for the function get_shcoeffs from aics-shparam. This function first converts the input binary image into a 3D triangular mesh using a traditional marching cubes algorithm from VTK Python library (Schroeder et al., 2018). To improve the quality of the output mesh, the binary input image is convolved with a Gaussian kernel with size σx=σy=σz=2, which is enough to smooth the image while retaining the overall cell and nuclear shape. Next, the mesh is translated to the origin and the coordinates of the mesh points are converted from cartesian to geographic coordinates (latitude, longitude and altitude). Altitude coordinates are then interpolated, using nearest neighbor, over a (lat,lon) spherical grid where each cell has a resolution of π/128. At this point, aics-shparam uses the Python package pyshtools (Wieczorek and Meschede, 2018) to expand, up to degree Lmax, the equally spaced grid into spherical harmonics coefficients using Driscoll and Healy’s sampling theorem (Driscoll et al., 1994). We used Lmax=16 as the SHE degree expansion to parameterize both cell and nuclear segmentation images. This was enough to guarantee a high fidelity mesh reconstruction, which can be quantified by the average distance between closest points in the original and reconstructed 3D meshes. We observed average distances of 0.33 µm +/− 0.1 µm for cells (n=300 randomly selected samples) and 0.12 µm +/ −0.02 µm for nucleus (n=300 randomly selected samples). Compared to the voxel size of our images (0.108 µm), we can say that Lmax=16 yields single pixel level precision for the nucleus and about three voxels precision for the cell, in average. This degree of expansion results in 289 coefficients for each input. Therefore, the shape of each cell in our dataset can be represented by a total of 578 coefficients (Figure 2A).

We can also recreate the 3D mesh representation of a particular set of SHE coefficients with aics-shparam. The Driscoll and Healy’s sampling theorem allows one to obtain a spherical grid from pre-computed SHE coefficients. These points on the spherical grid can be radially translated to their actual values in the grid to give rise to a 3D non-spherical shape.

#### Building the cell and nuclear shape space

##### Principal component analysis for dimensionality reduction

We used principal component analysis (PCA) to reduce the dimensionality of our joint vectors for all cells (578 SHE coefficients) down to eight principal components. We used the PCA implementation from the Python library scikit-learn (Pedregosa et al.) with default parameters (Figure 2B). Since the sign of a given principal component (PC) is arbitrary, we flipped the sign to ensure that the average volume of cells with negative PC values was less than that of cells with positive PC values. This was done independently for each PC.

##### Identifying the primary modes of shape variation

To prevent cells with extreme shapes from affecting the interpretation of the PCs, we excluded all cells that fell into the range 0th to 1st or 99th to 100th percentiles of each PC from subsequent analysis. These percentile ranges are shown by the vertical red lines in **Figure S3C**. The total number of cells left in the dataset was 175,935. We z-scored all PCs independently by dividing the PC values by the standard deviation (σ) of that PC. The probability distribution of each z-scored PC is shown in **Figure S3C**. The z-scored principal components are referred to as “shape modes” and the combination of the first 8 shape modes creates the 8-dimensional generative shape space used throughout this paper. We used the inverse of the PCA transform generated above to map shapes from the shape space back into SHE coefficients, which in turn, can be used to reconstruct the corresponding 3D shape. For example, the 8-components vector (0,0,0,0,0,0,0,0) represents the origin of the shape space and its corresponding 3D shape is called the mean cell shape throughout the paper (Figure 2C).

To systematically explore the shape space along each of the eight orthogonal axes, we let the elements of the 8-component array vary, one at the time, over discrete map points with values −2σ, - 1.5σ, −1.0σ, −0.5σ, 0, 0.5σ, 1.0σ, 1.5σ and 2.0σ. The combination of all eight shape modes and nine map points generates a grid of 8×9 3D shapes. Three different 2D views are used to visualize the 3D shapes. Top views represent the intersection of the 3D reconstructed mesh with the xy-plane, the equivalent of a single xy-slice through the center of the cell. In the same way, side views 1 and 2 represent the intersection of the 3D reconstructed shape with the xz- and yz-plane. To easily assign real cells to map points in the shape space, each shape mode is binned into nine bins of width 0.5σ, each centered around one map point, as represented by the black vertical lines in **Figure S3C**.

Cell and nuclear 3D mesh reconstructions using the inverse PCA transform are centered at the origin. Therefore, a few extra steps are required to translate the nuclear mesh back to its correct location relative to the center of the cell. We average all of the nuclear locations relative to their cell center for all the real cells within particular shape mode bin (**Figure S3C**). For example, to correct the nuclear location of the 3D mesh corresponding to the 8-components vector (0,0,0,0,0,-1.5σ,0,0) of Shape Mode 6 (Figure 2C), one would use the average location of all real cells that fall into the bin highlighted in blue in **Figure S3C**. Both the cell meshes and nuclear meshes with corrected locations for all shape modes are saved in VTK polydata format (Schroeder et al., 2018) for further analysis.

##### Alternative versions of the shape space

In addition to the joint cell and nuclear shape space, we also generated independent cell-only and nucleus-only shape spaces. For the cell-only shape space, the PCA was applied only on the cell SHE coefficients to reduce the data dimensionality from 289 to 8. For the nucleus-only shape space, images of DNA segmentation were aligned independently from any cell information. Nuclei were rotated such that the longest nuclear axis fell along the x-axis. The DNA segmentation was used to estimate the longest axis of a nucleus through a principal component analysis of the x and y coordinates of foreground voxels. The longest axis was defined as the direction of the first principal component and the alignment angle defined as the smallest angle between the longest axis and the x-axis. That nucleus was then rotated by the alignment angle such that the longest axis was aligned with the x-axis. Aligned images of nuclei were used as input for SHE coefficients calculation. PCA was applied only on the nuclear SHE coefficients to reduce the data dimensionality from 289 down to 8. After dimensionality reduction through PCA, these two alternative shape spaces were analyzed identically to the joint cell and nuclear shape space to identify the main modes of shape variation shown in **Figure SB&C**.

#### SHE coefficient-based parameterization and 3D morphing to build integrated average cells

##### Cytoplasmic and nuclear mapping

Pre-computed SHE coefficients were interpolated to morph the nuclear centroid mesh into the nuclear surface mesh and the nuclear surface mesh into the cell surface mesh. First, the nuclear centroid of each cell is described by the SHE coefficients representing a one-pixel radius (0.108 µm) 3D spherical mesh. These SHE coefficients representing the nuclear centroid and the pre-computed cell and nuclear SHE coefficients are concatenated and computationally described by a 3×289 matrix. This matrix is linearly interpolated to generate a 64×289 matrix. The interpolation is done by the function interp1d form scikit-learn in such a way that it guarantees that 1st, 3-th and 64th rows of the output matrix correspond exactly to SHE coefficients of centroid, nuclear and cell. SHE coefficients of each row of the interpolated matrix can be used to reconstruct corresponding 3D meshes. Meshes corresponding to rows 32 to 64 in the interpolated matrix are translated to a location that corresponds to a linear interpolation between nucleus and cell centroid. The visualization of subsequent 3D meshes (subsequent rows) causes the effect of mesh interpolation, as shown in Figure 3A, where we show only eight out of the 64 possible meshes (differently colored regions), including centroid (black dot) nuclear and cell meshes (represented by dashed lines).

##### Parameterized Intensity representation

Each of the 3D meshes is composed of points with xyz-coordinates. As the meshes are being generated from the interpolated matrix point by point, we can visit the corresponding xyz location in the aligned images that were used to generate the cell and nuclear SHE coefficients in the first place and associate the intensity value of that location with the mesh xyz coordinate. We can record either the original intensity values or the segmented intensity values since both types of images were aligned. The results can be organized as a matrix as shown in Figure 3A for the original FP signal. This matrix encodes a parameterized intensity representation of the cell.

This parameterized intensity representation can be used to reconstruct the aligned image that was used as the original input. We start with an empty image. We assign the value of each element of the parameterized intensity matrix to its closest xyz location in the empty image. We call this procedure voxelization and it produces a sparse representation of the original aligned image as shown in Figure 3A. The gaps in this image are due to the fact that our parameterized intensity representation samples only as many voxels of the original image as we have points in the 3D mesh. The gaps can be filled in by a nearest neighbor interpolation to produce an image that looks very similar to the original aligned image, as shown at the top of Figure 3A. We used the function NearestNDInterpolator from scikit-learn to perform the multidimensional nearest neighbor interpolation. We used the voxel-wise Pearson correlation coefficient in 3D to evaluate the similarity between reconstructed and original aligned images. We also performed an analysis between reconstructed and original aligned images on 32 randomly selected cells of all 25 cellular structures when the parameterized intensity representation is used to encode either original FP or segmented intensities (**Figure S3A&B**).

##### Generating morphed cells

The cellular mapping procedure described above only requires cell and nuclear SHE coefficients. Therefore, it can be applied to cell and nuclear shapes obtained for all map points of all shape modes. This is illustrated in Figure 3B for map point (0,0,1.5σ,0,0,0,0,0) of Shape Mode 3. The parameterized intensity representation of any given real cell can now be voxelized into any map point shape that underwent cellular mapping, to generate a morphed version of the real cell into that shape. This is illustrated in Figure 3 by morphing the FP signal from the real cell shown in panel (A) into the shape of map point (0,0,1.5σ,0,0,0,0,0) of Shape Mode 3 shown in panel (B). To prevent morphed cells from containing overly distorted signal intensity locations compared to the real cells, for instance by morphing a very flat real cell into a very tall shape (e.g. map point (2σ,0,0,0,0,0,0,0) of Shape Mode 1), we restrict our ourselves to apply the morphing only when the real cell shape and the map point shape are similar. This is achieved throughout the paper by allowing only cells of a given map point bin (**Figure S3C**) to be morphed into the corresponding map point shape. For example, only cells that fall into the bin highlighted in blue in **Figure S3C** are allowed to be morphed into the shape corresponding to map point (0,0,0,0,0,-1.5σ,0,0) of Shape Mode 6. The number of cells per structure available in each bin of each shape mode is shown in **Table S1**. We selected 300 randomly chosen cells (or the maximum number of cells available) per cellular structure and per shape mode and morphed these cells into their corresponding map point shapes. These morphed cells are stored as multichannel TIFF files and were used further for stereotypy analysis as described below.

##### Aggregating morphed cells

We compute the average and standard deviation of parameterized original FP and segmented intensity representations for all cells of each structure across map points of all shape modes of the shape space. This computation produces average and standard deviation parameterized intensity representations that could also be morphed into map point shapes of shape modes as described above. Results of these average and standard deviation images for all 25 cellular structures are shown as the first three columns of **Figure S3C** for map point (0,0,1.5σ,0,0,0,0,0) of Shape Mode 3. To quantify the location variation of each cellular structure, we normalized the standard deviation images by the average images to create coefficient of variation images. To prevent areas with very low average values (effectively very low original FP or segmented intensities) from greatly impacting the coefficient of variation, we defined the structure-localized coefficient of variation. The structure-localized coefficient of variation is computed as the coefficient of variation limited to a set of voxels containing intensities above a set threshold. The threshold was chosen to be the median of all non-zero voxels in the average image. Structure-localized coefficients of variation for all 25 cellular structures are shown as the 4th column of **Figure S3C** for map point (0,0,1.5σ,0,0,0,0,0) of Shape Mode 3. Aggregated cells are saved as 5D hyperstacks and used for visualization and concordance analysis, as described below.

##### Visualizing integrated average morphed cells

Average images of cellular structures morphed into the same map point shape are rendered simultaneously to illustrate the spatial relationships of different structures based on their average location in cells of a particular shape. Each volumetric channel of the 5D hyperstacks generated in the previous section for Shape Mode 3 was segmented using the default Surface option found in the Volume Viewer window of ChimeraX (Pettersen et al., 2020). Thresholds for each channel were selected manually to clarify dominant localization patterns observed in the voxel intensities.

#### Stereotypy calculation from morphed cells

We used morphed cell images to quantify the location stereotypy of a given cellular structure across different cells with similar shape. All 300 morphed cells available for each shape mode map point for each cellular structure were used to generate unique pairs of images. We calculated the voxel-wise Pearson correlation between all pairs of images, as illustrated in Figure 4A for lamin B1 (top) and mitochondria (bottom). The values of the resulting correlation coefficients represent a distribution of stereotypy values for each set of 300 cells. The distributions of stereotypy values for all 25 cellular structures for the mean cell shape are represented by the box plots in Figure 4B. The mean of the distribution of stereotypy values is called the mean stereotypy. The mean stereotypy values calculated for all 25 cellular structures across map points of all shape modes are shown as heatmaps in Figure 4C. To highlight the difference between mean stereotypy values relative to the mean cell shape, we created difference heatmaps as shown in **Figure S4C**, where the mean stereotypy of the mean cell bin is subtracted from the mean stereotypy of other map points.

#### Concordance calculation from average morphed cells

We used the 5D hyperstacks of average morphed images generated as described above to quantify the location concordance of all 25 cellular structures. The average morphed image of each structure for a given map point of a particular shape mode was used to build a voxel-wise correlation matrix as shown in Figure 5A. The element (i,j) of this matrix gives the concordance between structures i and j. The 25×25 correlation matrix is used as input for a hierarchical clustering algorithm to cluster all 25 cellular structures according to their relative concordance. We used the function cluster.hierarchy.linkage of type “average” from the Python package scipy (Virtanen, 2020) to produce the clustering represented by the dendrogram in Figure 5B calculated for the mean cell.

Concordance matrices were also calculated across map points for all shape modes. These matrices are represented by heatmaps across shape modes in Figure 5C, where the lower and upper triangle of each heatmap represent extreme opposite map points (see figure legend). To highlight the difference between concordance values relative to the mean cell, we create difference heatmaps as shown in **Figure S5B**.

#### Multiscale stereotypy and concordance analysis

Both stereotypy and concordance analysis were also performed across different spatial scales to investigate whether cellular structures display non-trivial behavior compared to what was observed in our initial analysis. Images for this analysis at different spatial scales were created by effectively downsampling the original images in all three dimensions by factors of 2 (**see Figure S4A**). The initial voxel-size of the morphed cell images was 0.108 µm. The downsampling process was repeated seven times to reach a voxel-size of ∼13.82 µm.

#### Cellular structure size scaling analysis

Statistical associations between volumes and areas of cells, nuclei and 15 cellular structures show how strongly these metrics are coupled to each and how they scale with respect to each other.

##### Description of data used for cellular structure size scaling analysis

This statistical analysis uses six metrics: The cell volume (µm^3^) and surface area (µm^2^), the nuclear volume (µm^3^) and surface area (µm^2^), the cytoplasmic volume (µm^3^), calculated by subtracting nuclear volume from cell volume, and the cellular structure volume (µm^3^). The cell and nuclear metrics are available for all cells (n=203,737) and calculated based on the segmentation of the cell and nucleus, respectively. The cellular structure volume is based on the segmentation of the FP-tagged structure in the cell and is applied to the 15 cellular structures validated for structure volume analysis. (see “Structure segmentation” section; **Table S1**). If multiple pieces (connected components) of the structure are present in this cell, structure volume gives the total volume of all connected components.

##### Linear regression model to compute statistical coupling between metrics

We employed a simple linear regression model (y = ax + b) to compute the amount of explained variance in the dependent variable y by the independent variable x. Linear regression models were calculated with x as one of the five cell and nuclear metrics (cell volume and area, nuclear volume and area, cytoplasmic volume) and y as one of all six metrics (including cellular structure volume). In the case of structure volume, the model was computed for each structure separately, using only those cells that correspond to the structure in question. The explained variance in y due to x, or the R2 statistic (coefficient of determination), was computed for all models and is shown in Figure 6B. We used a bootstrap analysis (n=100 bootstraps) to calculate the 5-95% confidence interval, visualized as horizontal error bars in Figure 6H.

##### Linear regression model to compute cellular structure scaling rates

Using the same simple linear regression model (y = ax + b), we calculated the “scaling rate” of each cellular structure relative to cell volume. The scaling rate gives the increase in volume (or area) of a cellular structure as cell size is doubled. In this case x is cell volume and y is one of the other five metrics. Using a histogram density estimation of cell volume, we determined the interval with the most cells where the cell volume doubles. This interval is from x0 = 1160 µm^3^ to x1 = 2320 µm^3^. These x values are then evaluated with the learned regression model to get the corresponding y values, termed y0 and y1. The scaling rate is computed as (y1-y0)/y0 * 100%. Figure 6B depicts this process to compute the scaling rate for nuclear volume. In this case y0 is 346 µm^3^ and y1 is 669 µm^3^, giving a scaling rate of 93%. The scaling rates across all metrics is given in Figure 6A. We used a bootstrap analysis (n=100 bootstraps) to calculate the 5-95% confidence interval, visualized as vertical error bars in Figure 6H.

##### Multivariate regression model to isolate the effect of cell and nuclear metrics in explaining structure volumes

Cell and nuclear metrics show a large degree of collinearity, which makes it non-trivial to isolate the effect of one particular cell or nuclear metric on structure volume. We used multivariate regression models to isolate the effect of cell and nuclear metrics. In contrast to univariate regression models (y = ax + b, where is x a vector and a is scalar), multivariate models have multiple dependent variables (y = aX + b, where X is a matrix with p columns and a is a vector with p entries). We first computed the explained variance in cellular structure volume using cell volume, cell surface area, nuclear volume and nuclear surface area as independent variables. Note that cytoplasmic volume is a linear combination of cell and nuclear volumes and does not need to be added to the model. Then, we remove a single metric or a pair of metrics from the independent variables and recalculate the model. The “unique explained variance” ascribed to the metric or pair of metrics is calculated as the difference in explained variance between the full model, i.e. containing the four metrics and the model where the metric or pair of metrics was left out. Specifically, the metrics (pairs) for which this unique explained variance was computed were cell volume and cell surface area (cell v+a), cell volume, cell surface area, nuclear volume and nuclear surface area (nuc v+a), nuclear volume, and nuclear surface area. The total explained variance (using all four metrics) as well as the unique explained variance portions are depicted in Figure 6A.

##### Non-linear regression models to compute statistical coupling between metrics

For each of the linear regression models described above, we also computed a more complex, non-linear model. Specifically, given a linear regression model y = aXl + b, where the design matrix Xl contains either a single vector or multiple columns, we expanded the design matrix Xl using two steps: 1) for all pairs of columns in Xl, we computed the pointwise product and added these new columns to the design matrix; and 2) for each column in the design matrix we added four copies and raised the values of these new columns to the following four powers: 1/3 (cube root), 1/2 (square root), 2 (square), 3 (cube). The resulting design matrix, Xc, was then used in the linear regression model y = aXc + b to compute the explained variances. A visualization of the explained variances using simple regression models compared with the non-linear models with interaction effects is shown in **Figure S6B**.

##### Visualization of bi-variate association using scatter plots

Associations between pairs of metrics were visualized in scatter plots, where each cell is plotted as a point in the two-dimensional space spanned by the two metrics, x (on the x-axis) and y (on the y-axis). The number of cells is stated in the upper left corner. The regression model is depicted as a gray straight line (y = ax + b) and the explained variance in y due to x (the R2 statistic) is also stated in the upper left corner. There are two additional graphical aspects to improve the interpretation of these bi-variate associations: 1) A green line is shown that depicts the running average. Briefly, the values of metric x are binned in 100 equally spaced bins. For each of these bins, the mean value for metric y is computed from all cells in that bin, i.e. unless the number of cells in the bin is below 50 in which case no value is recorded. The green line is the running average of metric y as a function of the bin centers. 2) Cells are colored according to a density estimate. Briefly, a kernel density estimate is performed in the two-dimensional space. Based on this estimation, each cell is assigned a probability. The probabilities are transformed to cumulative probabilities and normalized, such that the cell with the highest probability, i.e. the one within the highest density region, gets a value of 1. By aligning the probabilities with a colormap, cells are colored to convey the density. The use of cumulative probabilities ensures that the colors have the same interpretation across different plots, i.e. different metrics. See Figure 6.

#### Generalizability analysis

To test the generalizability of our multi-part analysis approach of the locations, amounts, and their variation of the 25 cellular structures we ran the main analyses shown in this paper on subsets of the analysis dataset where we selected (by downsampling) a much smaller number of cells per structure.

##### Shape space generalizability

Using the main analysis dataset (n = 203,737 cells) including the 578 SHE coefficients, we randomly selected 300 cells and we applied PCA on the 300×578 table to reduce its dimensionality down to eight principal components. The resulting shape space is analyzed identically to the main shape space.

##### Stereotypy and concordance generalizability

We calculate the mean stereotypy of all 25 cellular structures morphed into the mean cell for different numbers of pairs of morphed cells. We varied the number of pairs of morphed cells used to average the Pearson correlation scores from 2 to 300 with a step size of one (Figure 7B). By visual inspection we determined the minimum number of pairs of cells required to recover the ranking of cellular structure mean stereotypy from 300 pairs of cells to be 35 pairs. The morphed cells used here were randomly sampled from the 300 morphed cells available per cellular structure per map point of each shape mode generated as described in the “Generating morphed cells” section above.

For location concordance, we selected a set of 300 cells chosen at random for each cellular structure within the mean cell shape bin (except n= 252 for nuclear speckles, see **DataFile S1**). We used the 5D hyperstacks of average morphed cell images from this downsampled dataset as described above to calculate the location concordance.

##### Downsampling cellular structure size scaling analysis

We created downsampled versions of the dataset with n cells per structure randomly selected (n=10, 20, 30, 50, 100, 200, 300, 500, 1000, 1500), each with three repeats. The regression models to compute explained variances and scaling rates were recalculated on these downsampled versions of dataset. Figure 7D shows these statistics for a single repeat of n=300. This figure also shows how the recalculated numbers differ from the original numbers as a function of the number of cells per structure.

### ADDITIONAL RESOURCES

The Allen Cell Collection, the hiPSC Single-Cell Image Dataset, protocols, the Allen Cell Discussion Forum and additional information can be found here: (https://www.allencell.org/)

### RESOURCES TABLE

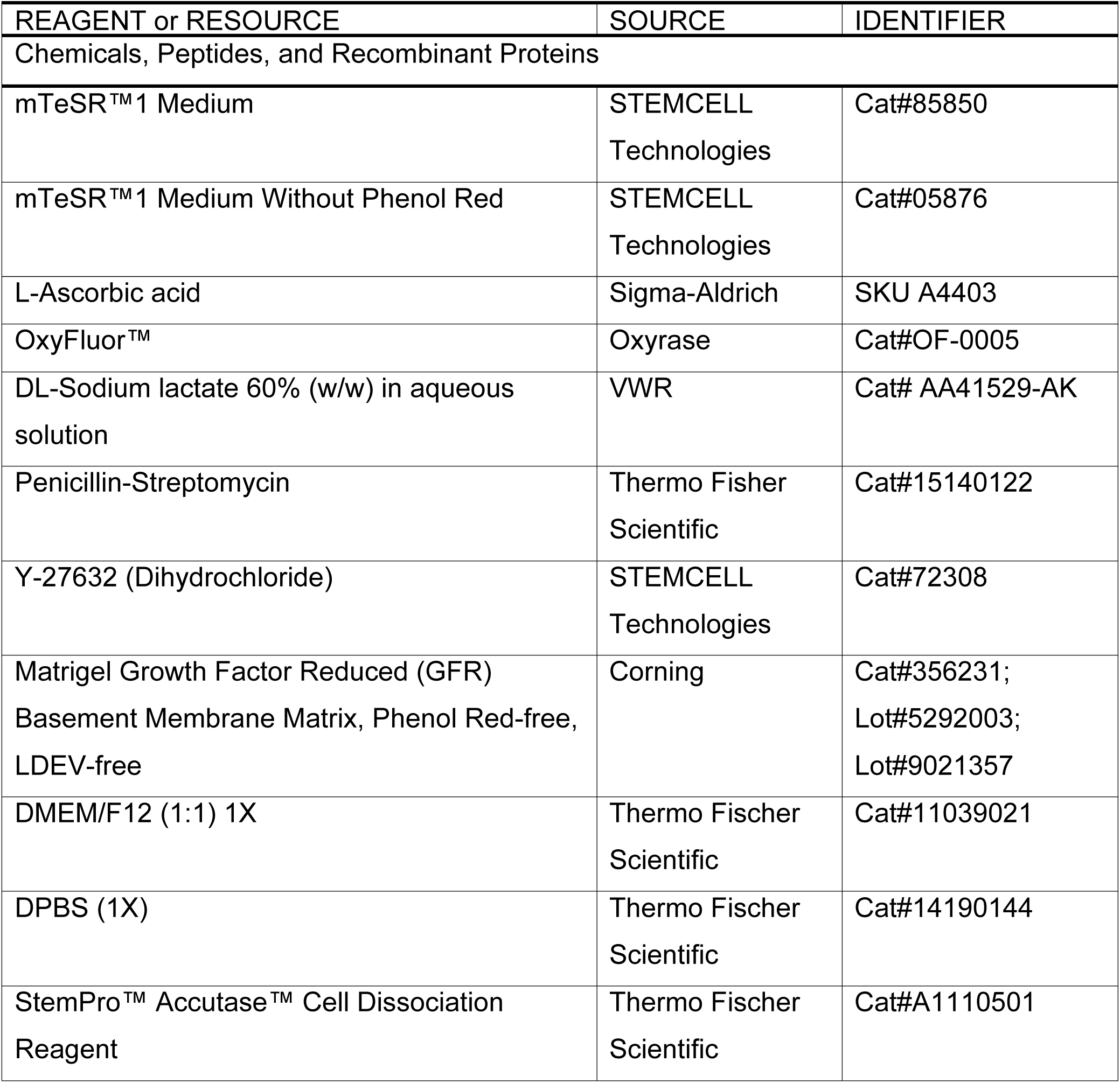

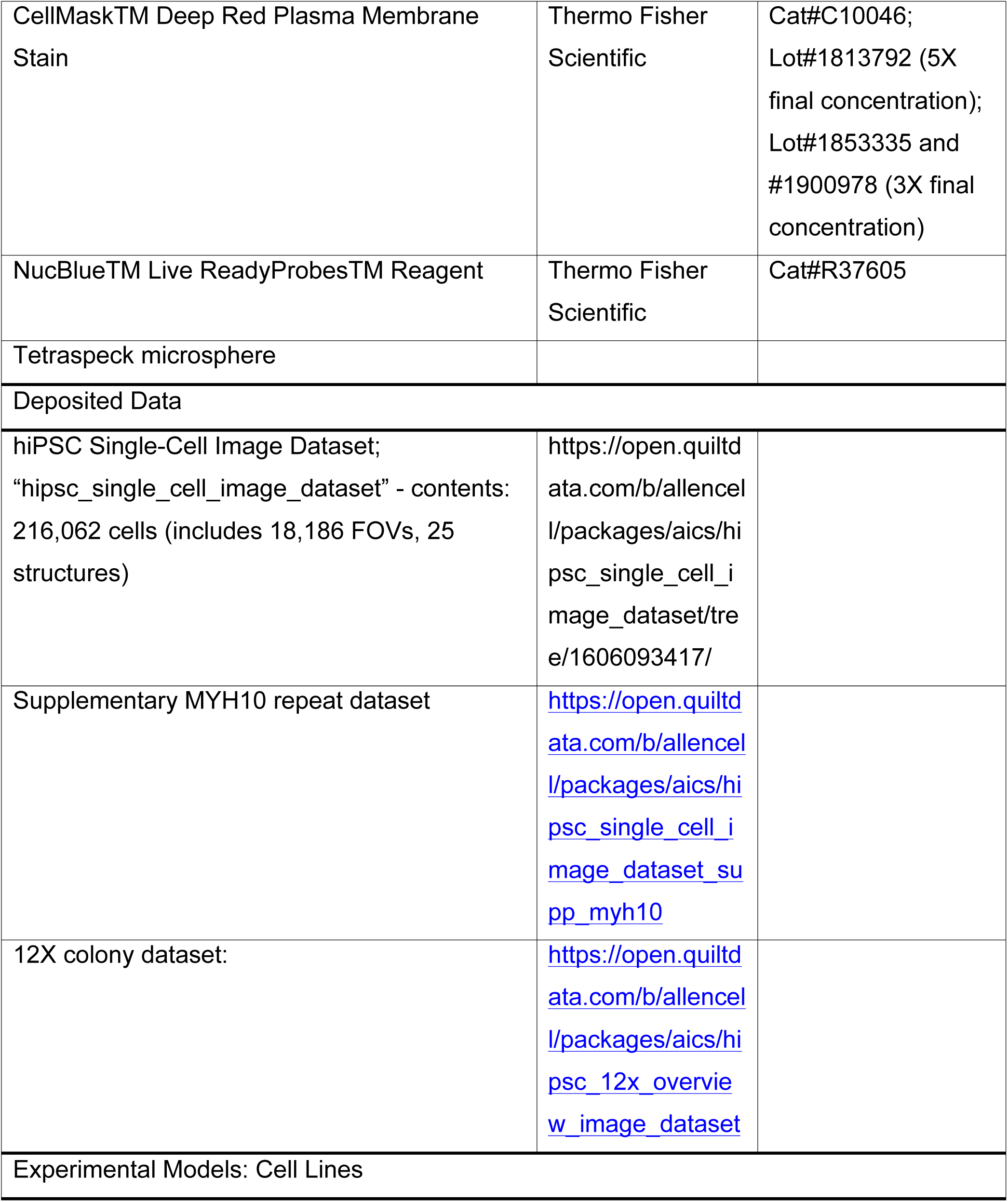

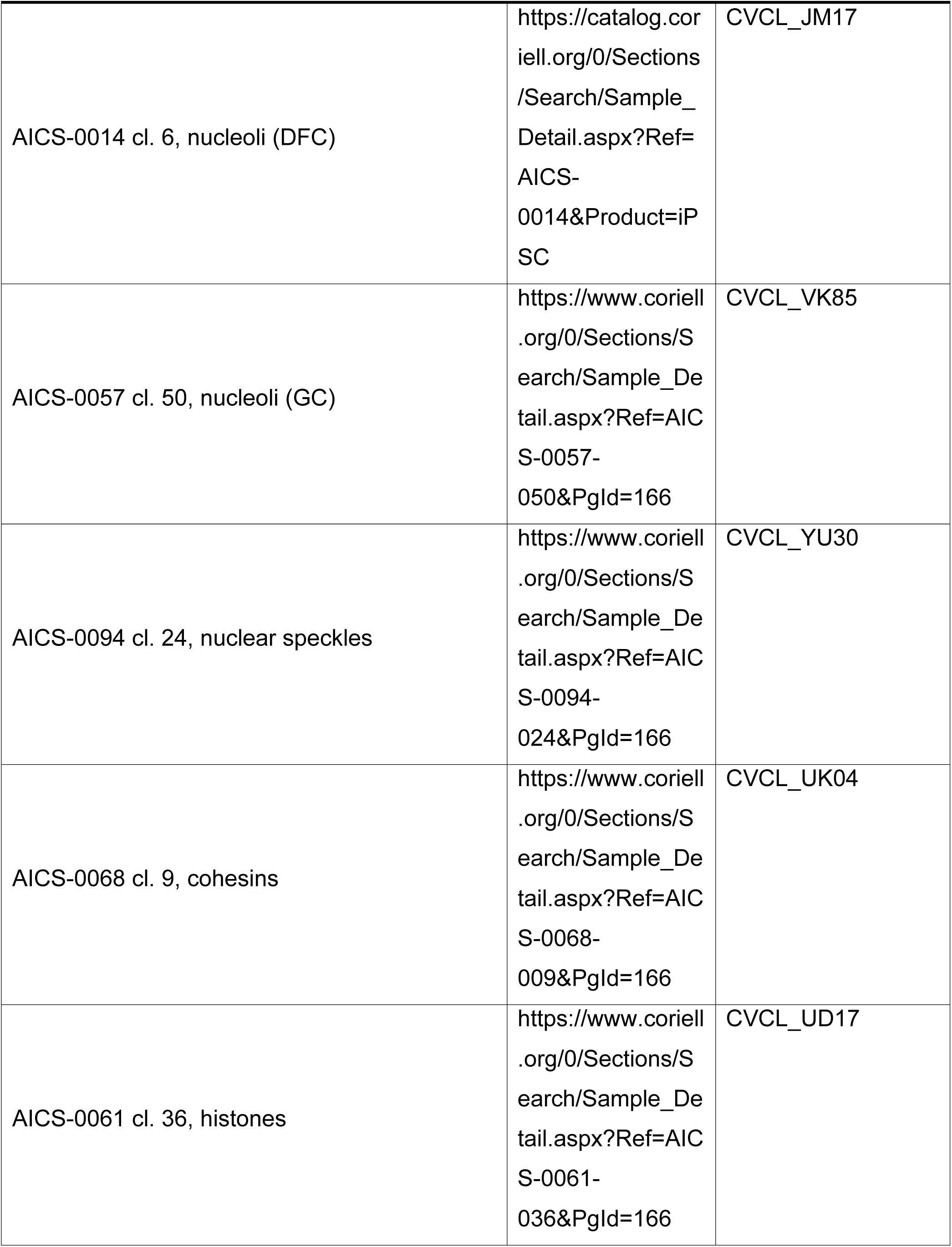

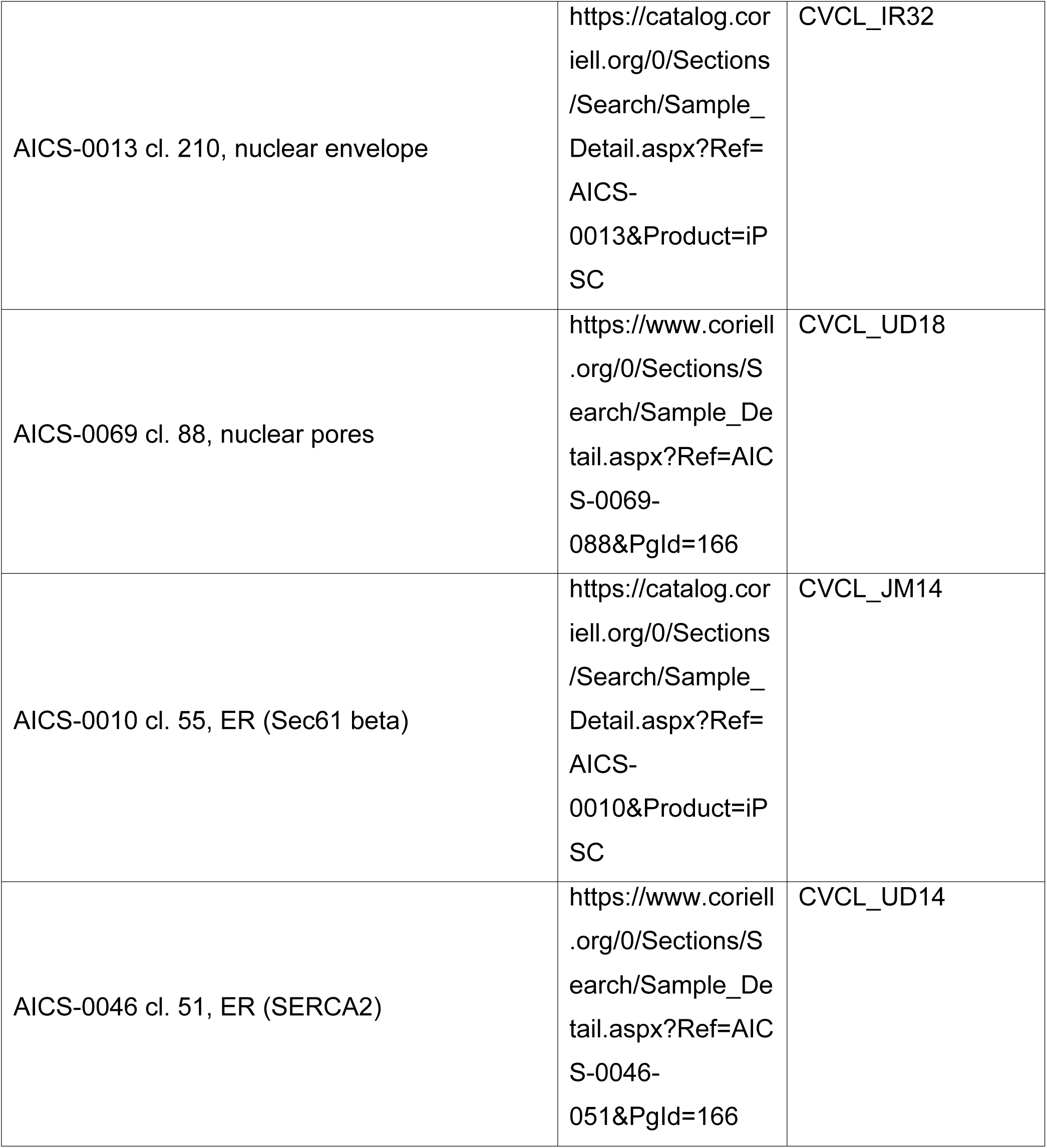

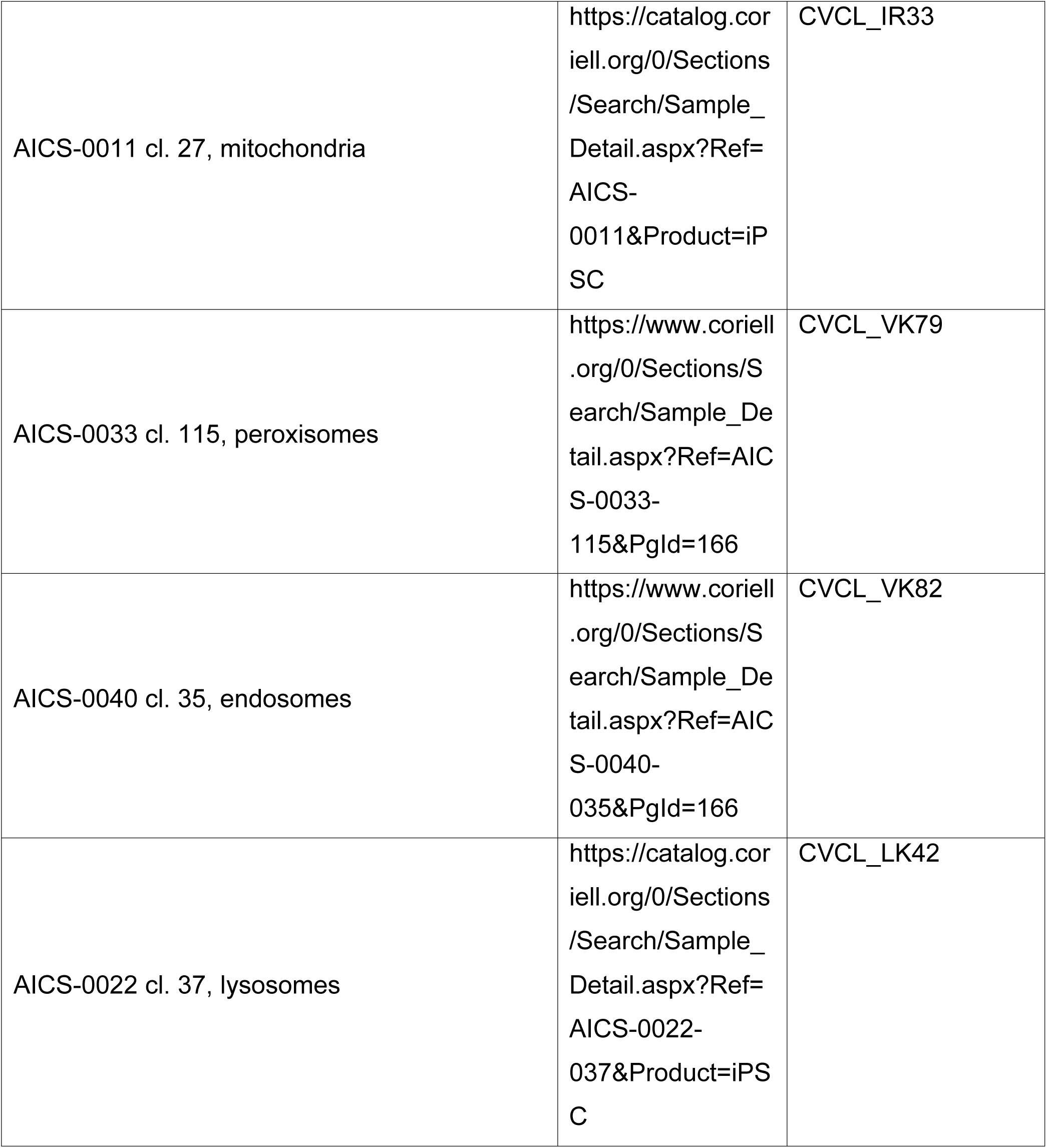

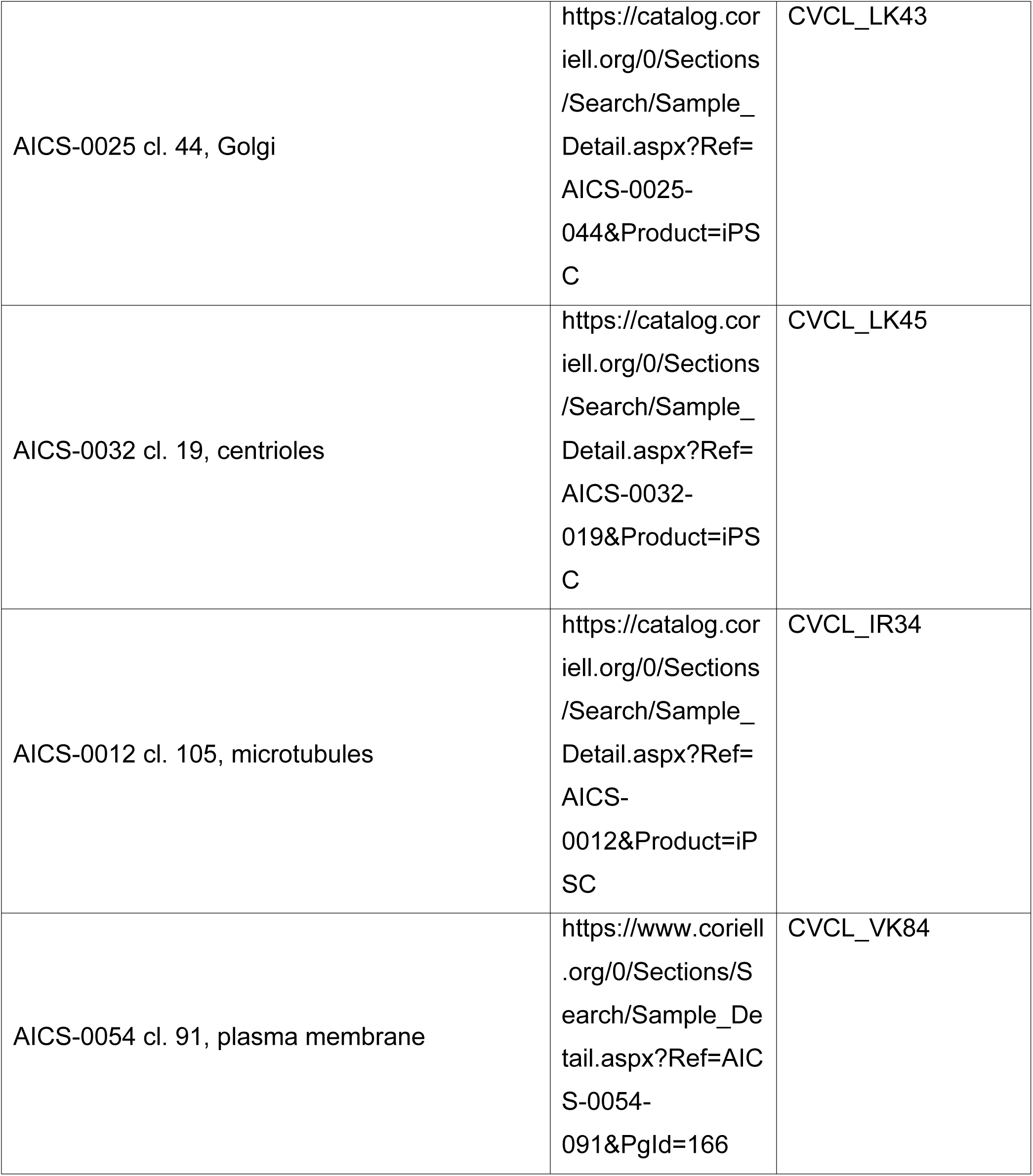

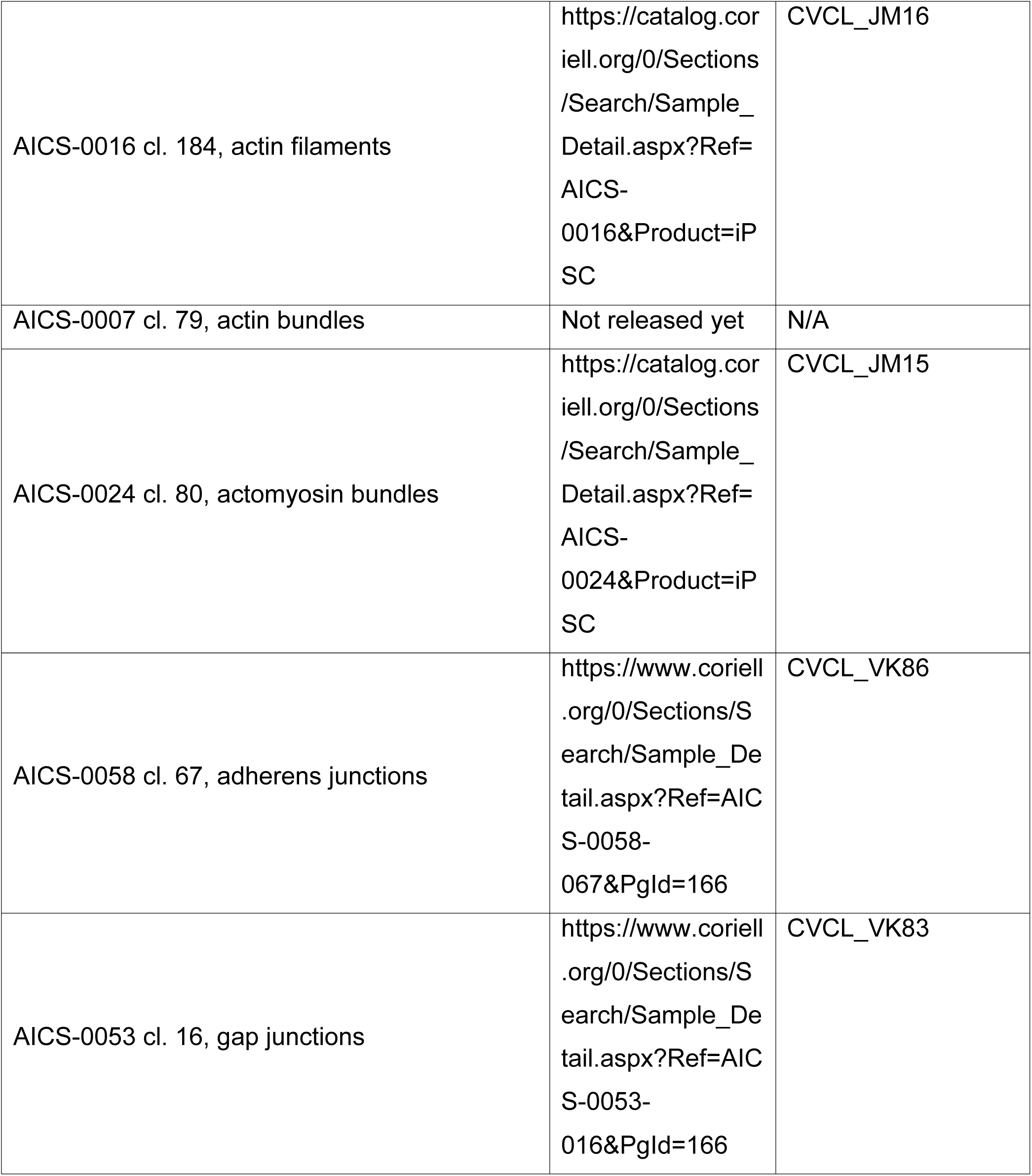

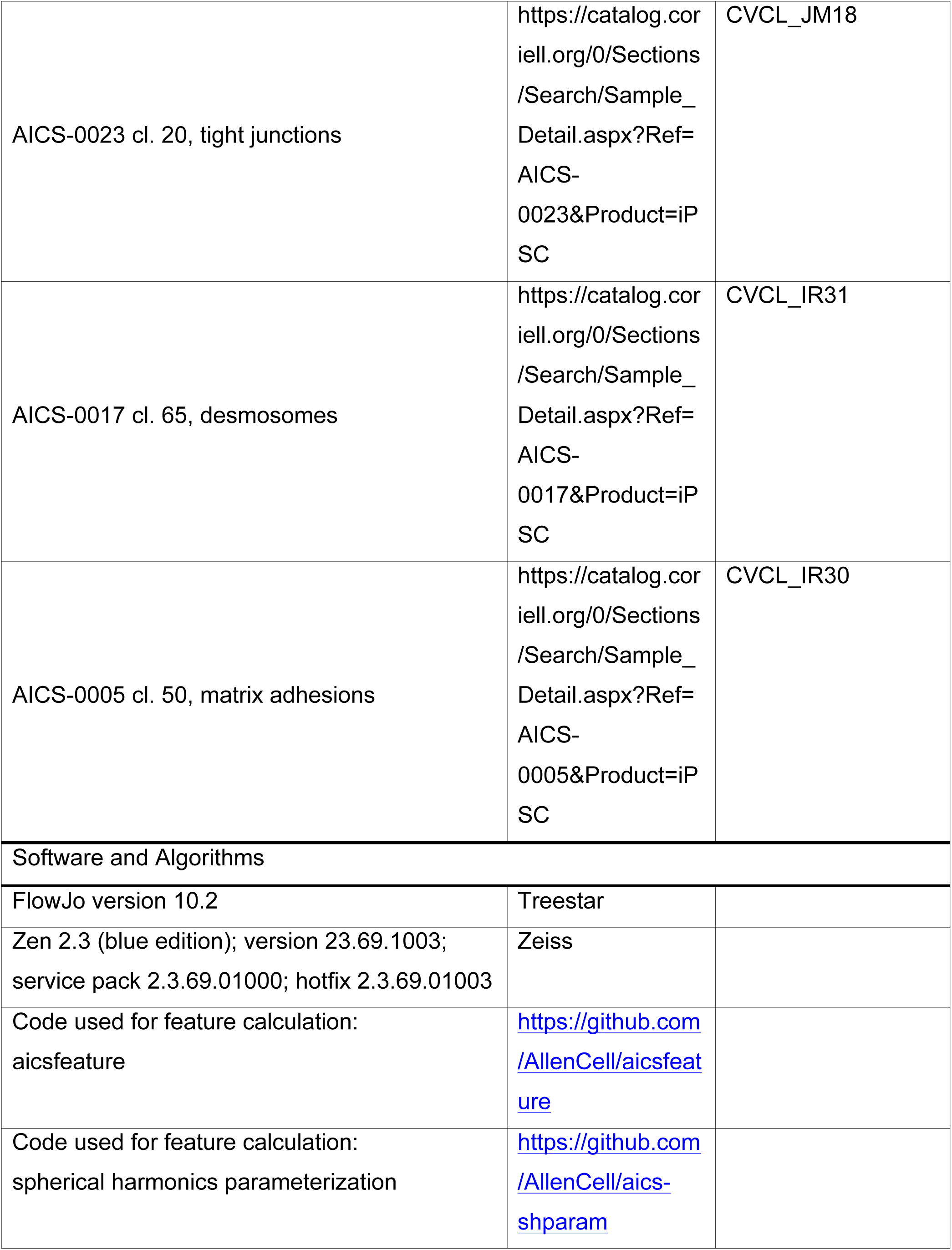

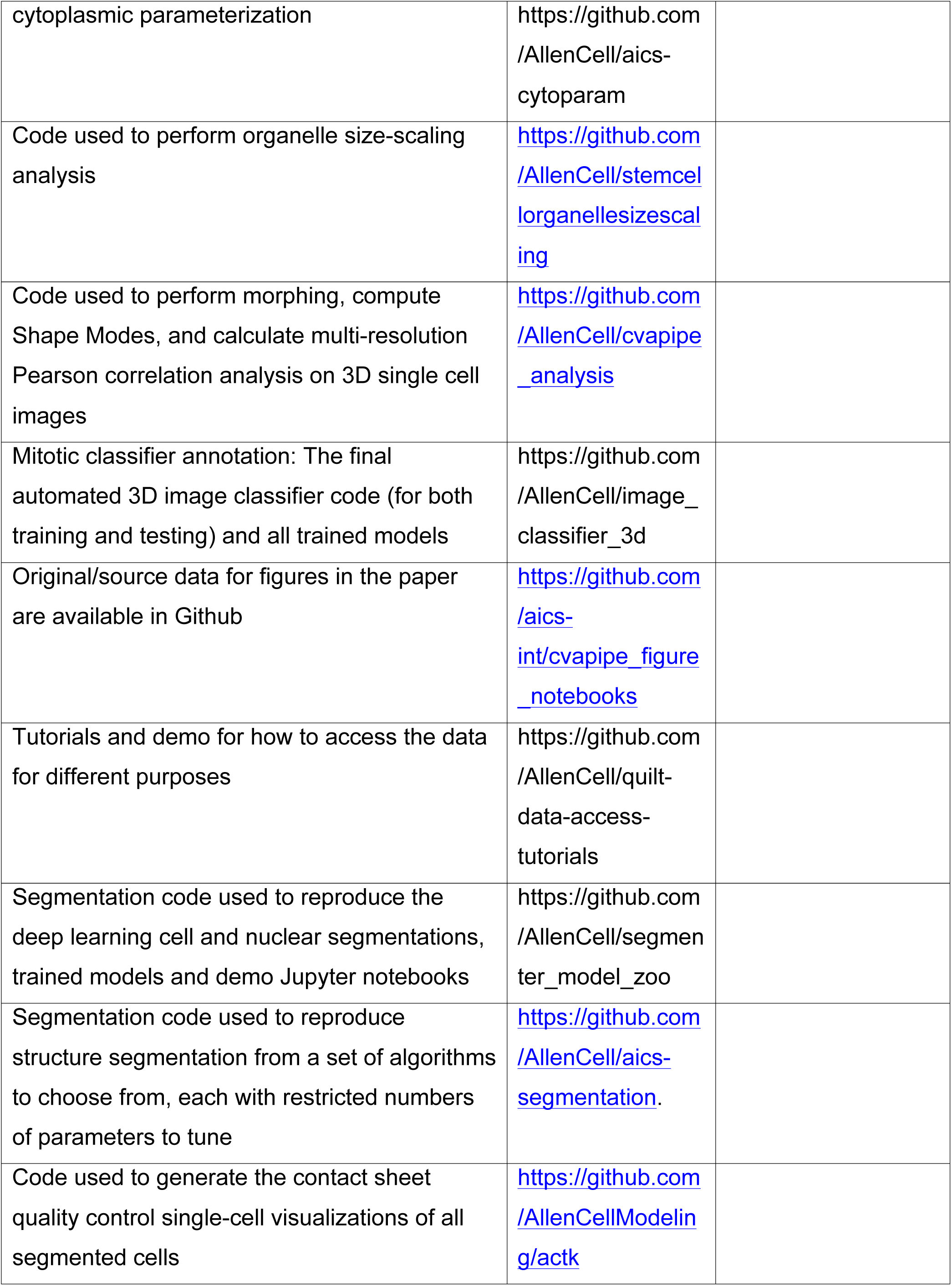

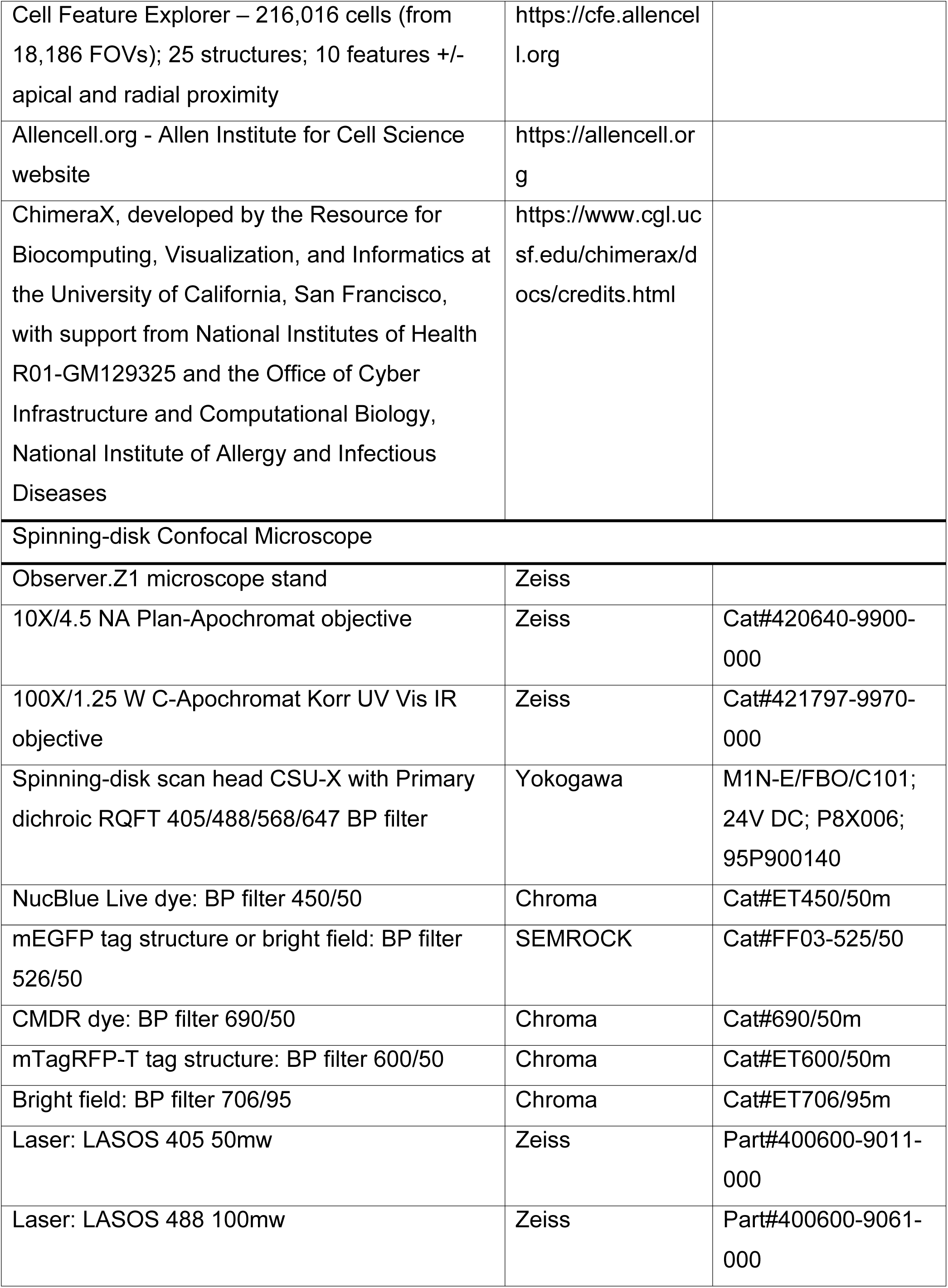

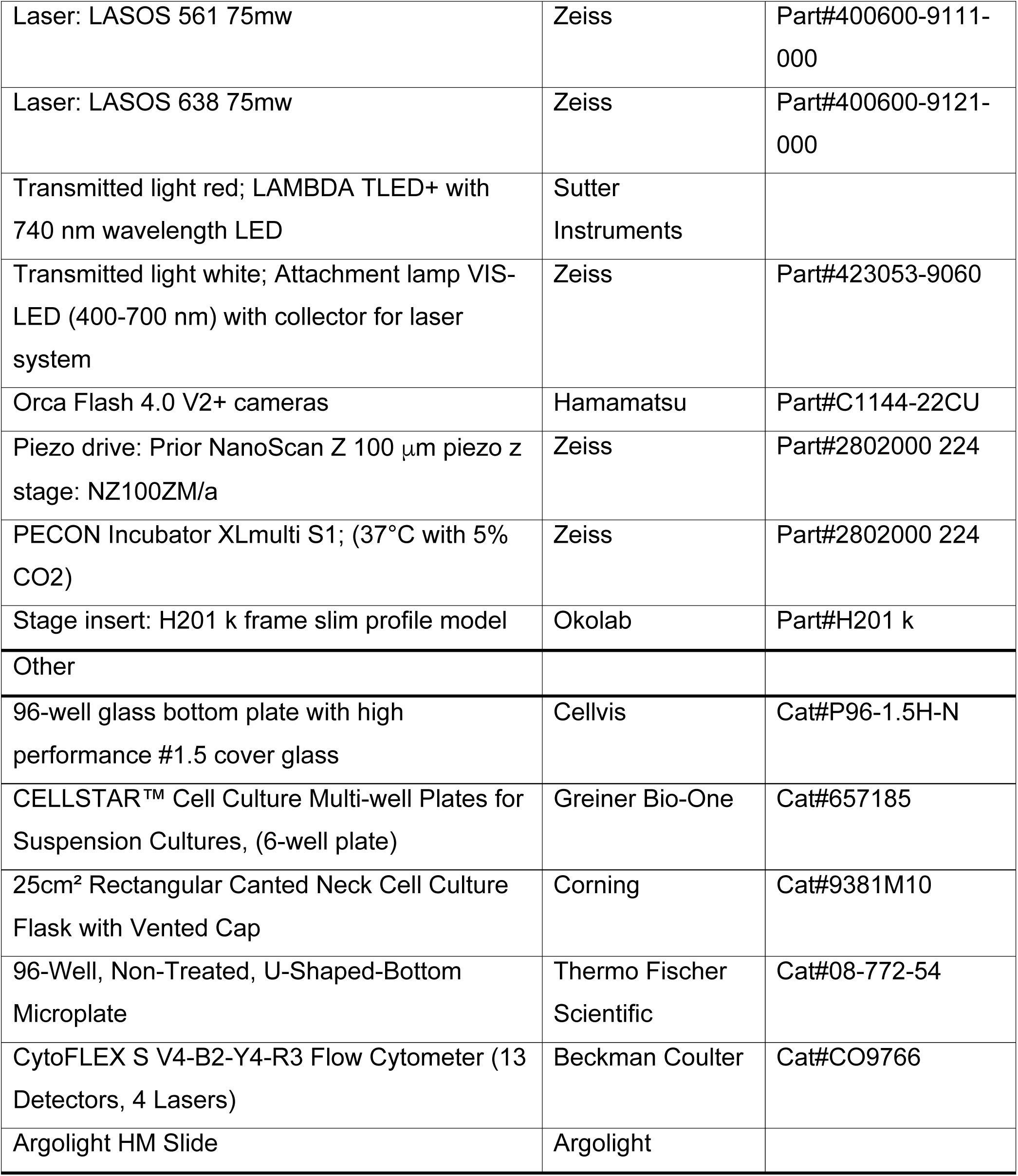

## References

Aldridge, S., and Teichmann, S.A. (2020). Single cell transcriptomics comes of age. Nature Communications 11, 4307.

Baghbaderani, A.A., Tian, X., Neo, B.H., Burkall, A., Dimezzo, T., Sierra, G., Zeng, X., Warren, K., Kovarcik, D.P., Fellner, T., et al. (2015). cGMP-Manufactured Human Induced Pluripotent Stem Cells Are Available for Pre-clinical and Clinical Applications. Stem Cell Reports 5, 647–659.

Caicedo, J.C., Cooper, S., Heigwer, F., Warchal, S., Qiu, P., Molnar, C., Vasilevich, A.S., Barry, J.D., Bansal, H.S., Kraus, O., et al. (2017). Data-analysis strategies for image-based cell profiling. Nat Methods 14, 849–863.

Chen, J., Ding, L., Viana, M.P., Hendershott, M.C., Yang, R., Mueller, I.A., and Rafelski, S.M. (2018). The Allen Cell Structure Segmenter: a new open source toolkit for segmenting 3D intracellular structures in fluorescence microscopy images. BioRxiv 491035.

Coston, M.E., Gregor, B.W., Arakaki, J., Borensztejn, A., Do, T.P., Fuqua, M.A., Haupt, A., Hendershott, M.C., Leung, W., Mueller, I.A., et al. (2020). Automated hiPSC culture and sample preparation for 3D live cell microscopy. BioRxiv 2020.12.18.423371.

Drubin, D.G., and Hyman, A.A. (2017). Stem cells: the new “model organism.” Mol Biol Cell 28, 1409– 1411.

Falcon, W., and Cho, K. (2020). A Framework For Contrastive Self-Supervised Learning And Designing A New Approach. ArXiv:2009.00104 [Cs].

Fransen, M., Lismont, C., and Walton, P. (2017). The Peroxisome-Mitochondria Connection: How and Why? Int J Mol Sci 18.

Gerbin, K.A., Grancharova, T., Donovan-Maiye, R., Hendershott, M.C., Brown, J., Dinh, S.Q., Gehring, J.L., Hirano, M., Johnson, G.R., Nath, A., et al. (2020). Cell states beyond transcriptomics: integrating structural organization and gene expression in hiPSC-derived cardiomyocytes. BioRxiv 2020.05.26.081083.

Gut, G., Herrmann, M.D., and Pelkmans, L. (2018). Multiplexed protein maps link subcellular organization to cellular states. Science 361.

Hao, F., Kondo, K., Itoh, T., Ikari, S., Nada, S., Okada, M., and Noda, T. (2018). Rheb localized on the Golgi membrane activates lysosome-localized mTORC1 at the Golgi–lysosome contact site. J Cell Sci 131.

Harris, C.R., Millman, K.J., van der Walt, S.J., Gommers, R., Virtanen, P., Cournapeau, D., Wieser, E., Taylor, J., Berg, S., Smith, N.J., et al. (2020). Array programming with NumPy. Nature 585, 357– 362.

Hockemeyer, D., Soldner, F., Beard, C., Gao, Q., Mitalipova, M., DeKelver, R.C., Katibah, G.E., Amora, R., Boydston, E.A., Zeitler, B., et al. (2009). Efficient targeting of expressed and silent genes in human ESCs and iPSCs using zinc-finger nucleases. Nature Biotechnology 27, 851–857.

Hunter, J.D. (2007). Matplotlib: A 2D Graphics Environment. Computing in Science Engineering 9, 90–95.

Johnson, G.T., Autin, L., Al-Alusi, M., Goodsell, D.S., Sanner, M.F., and Olson, A.J. (2015). cellPACK: a virtual mesoscope to model and visualize structural systems biology. Nature Methods 12, 85–91.

Kreitzer, F.R., Salomonis, N., Sheehan, A., Huang, M., Park, J.S., Spindler, M.J., Lizarraga, P., Weiss, W.A., So, P.-L., and Conklin, B.R. (2013). A robust method to derive functional neural crest cells from human pluripotent stem cells. Am J Stem Cells 2, 119–131.

Lauffenburger, D.A., and Horwitz, A.F. (1996). Cell migration: a physically integrated molecular process. Cell 84, 359–369.

Liaw, A., and Wiener, M. (2002). Classification and Regression by randomForest. 2, 5.

Macklin, D.N., Ahn-Horst, T.A., Choi, H., Ruggero, N.A., Carrera, J., Mason, J.C., Sun, G., Agmon, E., DeFelice, M.M., Maayan, I., et al. (2020). Simultaneous cross-evaluation of heterogeneous E. coli datasets via mechanistic simulation. Science 369.

Marshall, W.F. (2020). Scaling of Subcellular Structures. Annu. Rev. Cell Dev. Biol. 36, 219–236.

Marshall, W.F., Dernburg, A.F., Harmon, B., Agard, D.A., and Sedat, J.W. (1996). Specific interactions of chromatin with the nuclear envelope: positional determination within the nucleus in Drosophila melanogaster. Mol Biol Cell 7, 825–842.

McCormick, M.M., Liu, X., Ibanez, L., Jomier, J., and Marion, C. (2014). ITK: enabling reproducible research and open science. Front. Neuroinform. 8.

McKinney, W. (2011). “pandas: a foundational Pythonlibrary for data analysis and statistics”. In In:Pythonfor High Performance and Scientific Computing, p. 14.9.

Nicholas Sofroniew, Kira Evans, Juan Nunez-Iglesias, Ahmet Can Solak, Talley Lambert, kevinyamauchi, Jeremy Freeman, Loic Royer, Shannon Axelrod, Peter Boone, et al. (2019). napari/napari: 0.2.8 (Zenodo).

Oceguera-Yanez, F., Kim, S.-I., Matsumoto, T., Tan, G.W., Xiang, L., Hatani, T., Kondo, T., Ikeya, M., Yoshida, Y., Inoue, H., et al. (2016). Engineering the AAVS1 locus for consistent and scalable transgene expression in human iPSCs and their differentiated derivatives. Methods 101, 43–55.

Ounkomol, C., Seshamani, S., Maleckar, M.M., Collman, F., and Johnson, G.R. (2018). Label-free prediction of three-dimensional fluorescence images from transmitted-light microscopy. Nature Methods 15, 917–920.

Paszke, A., Gross, S., Massa, F., Lerer, A., Bradbury, J., Chanan, G., Killeen, T., Lin, Z., Gimelshein, N., Antiga, L., et al. (2019). PyTorch: An Imperative Style, High-Performance Deep Learning Library. Advances in Neural Information Processing Systems 32, 8026–8037.

Pedregosa, F., Varoquaux, G., Gramfort, A., Michel, V., Thirion, B., Grisel, O., Blondel, M., Prettenhofer, P., Weiss, R., Dubourg, V., et al. Scikit-learn: Machine Learning in Python. MACHINE LEARNING IN PYTHON 6.

Pettersen, E.F., Goddard, T.D., Huang, C.C., Meng, E.C., Couch, G.S., Croll, T.I., Morris, J.H., and Ferrin, T.E. (2020). UCSF ChimeraX: Structure visualization for researchers, educators, and developers. Protein Sci.

Pincus, Z., and Theriot, J.A. (2007). Comparison of quantitative methods for cell-shape analysis. J Microsc 227, 140–156.

Roberts, B., Haupt, A., Tucker, A., Grancharova, T., Arakaki, J., Fuqua, M.A., Nelson, A., Hookway, C., Ludmann, S.A., Mueller, I.A., et al. (2017a). Systematic gene tagging using CRISPR/Cas9 in human stem cells to illuminate cell organization. Mol Biol Cell 28, 2854–2874.

Roberts, B., Haupt, A., Tucker, A., Grancharova, T., Arakaki, J., Fuqua, M.A., Nelson, A., Hookway, C., Ludmann, S.A., Mueller, I.A., et al. (2017b). Systematic gene tagging using CRISPR/Cas9 in human stem cells to illuminate cell organization. Mol Biol Cell 28, 2854–2874.

Rocklin, M. (2015). Dask: Parallel Computation with Blocked algorithms and Task Scheduling. (Austin, Texas), pp. 126–132.

Roggiani, M., and Goulian, M. (2015). Oxygen-Dependent Cell-to-Cell Variability in the Output of the Escherichia coli Tor Phosphorelay. Journal of Bacteriology 197, 1976–1987.

Ruan, X., and Murphy, R.F. (2019). Evaluation of methods for generative modeling of cell and nuclear shape. Bioinformatics 35, 2475–2485.

Schroeder, W., Martin, K., and Lorensen, B. (2018). The Visualization ToolkitAn Object-Oriented Approach To 3D Graphics.

Shen, L., Farid, H., and McPeek, M. (2009). Modeling three-dimensional morphological structures using spherical harmonics. Evolution 63, 1003–1016.

Thul, P.J., Åkesson, L., Wiking, M., Mahdessian, D., Geladaki, A., Ait Blal, H., Alm, T., Asplund, A., Björk, L., Breckels, L.M., et al. (2017). A subcellular map of the human proteome. Science 356.

Valencia, P., Dias, A.P., and Reed, R. (2008). Splicing promotes rapid and efficient mRNA export in mammalian cells. Proc Natl Acad Sci U S A 105, 3386–3391.

Virtanen, P. (2020). SciPy 1.0: fundamental algorithms for scientific computing in Python / Nature Methods.

Walt, S. van der, Schönberger, J.L., Nunez-Iglesias, J., Boulogne, F., Warner, J.D., Yager, N., Gouillart, E., and Yu, T. (2014). scikit-image: image processing in Python. PeerJ 2, e453.

Wang, T., and Hong, W. (2002). Interorganellar Regulation of Lysosome Positioning by the Golgi Apparatus through Rab34 Interaction with Rab-interacting Lysosomal Protein. Mol Biol Cell 13, 4317– 4332.

Wieczorek, M.A., and Meschede, M. (2018). SHTools: Tools for Working with Spherical Harmonics. Geochemistry, Geophysics, Geosystems 19, 2574–2592.

