## Supplemental Information and figures for "Robust integrated intracellular organization of the human iPS cell: where, how much, and how variable"

**Supplemental information in this PDF includes:**

Figures S1 - S7

Table S1

Captions for Movies S1- S4

**Other Supplementary Materials for this manuscript include:**

Movies S1 - S4

Data File S1

Figure S1

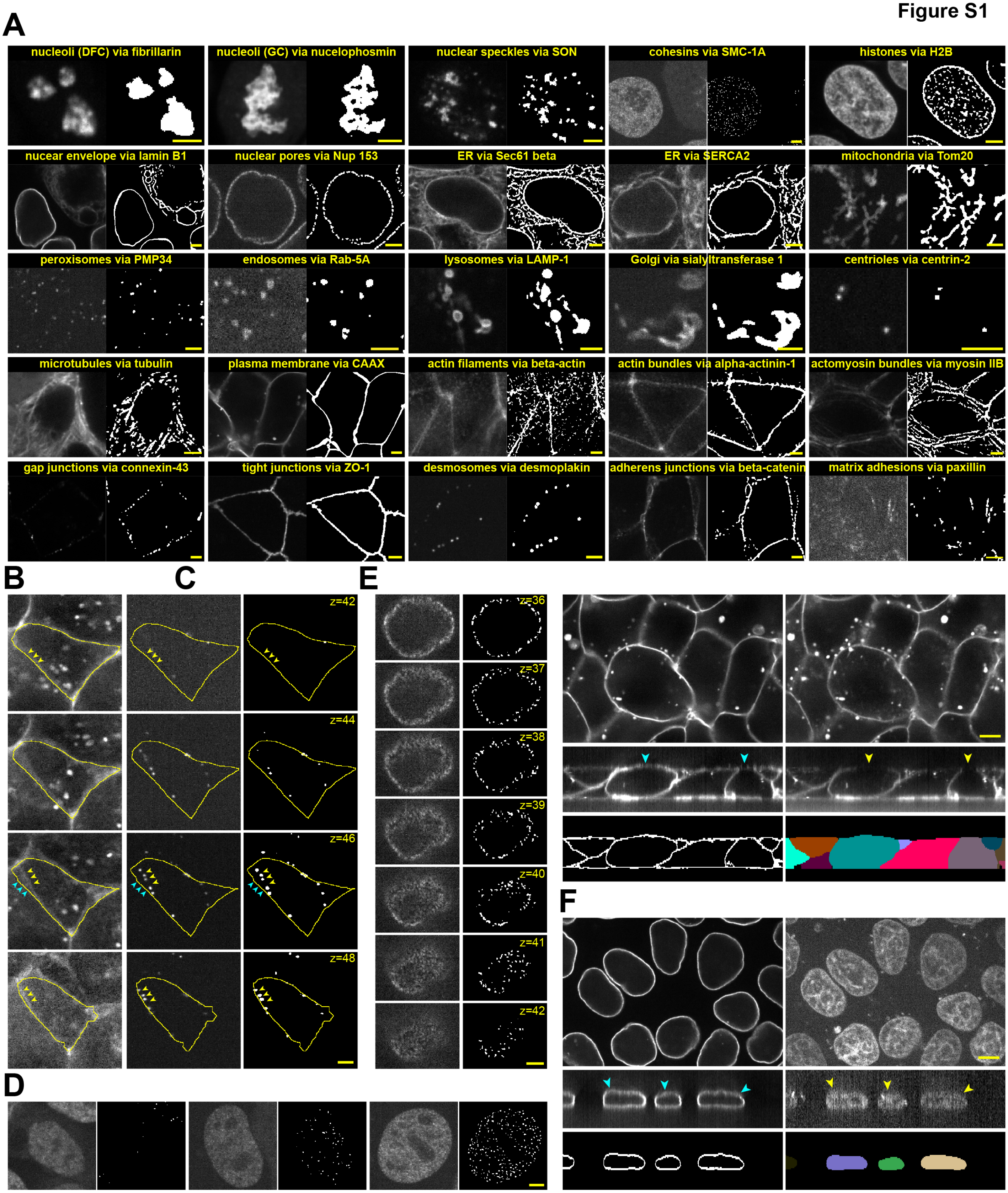
A
nucleoli (DFC) via fibrillarin
nucleoli (GC) via nucelophosmin
nuclear speckles via SON
cohesins via SMC-1A
histones via H2B
nucear envelope via lamin B1
nuclear pores via Nup 153
ER via Sec61 beta
ER via SERCA2
mitochondria via Tom20
peroxisomes via PMP34
endosomes via Rab-5A
lysosomes via LAMP-1
Golgi via sialyltransferase 1
centrioles via centrin-2
microtubules via tubulin
plasma membrane via CAAX
actin filaments via beta-actin
actin bundles via alpha-actinin-1
actomyosin bundles via myosin IIB
gap junctions via connexin-43
tight junctions via ZO-1
desmosomes via desmoplakin
adherens junctions via beta-catenin
matrix adhesions via paxillin
B
C
z=42
z=44
z=46
z=48
E
z=36
z=37
z=38
z=39
z=40
z=41
z=42
F
D

**Figure S1:** Supplemental figure for **Figure 1**. **A)** Overview panel showing example images of each of the 25 cellular structures in the hiPSC Single-Cell Image Dataset. For each pair of images, the left image shows a representative single z-slice of the FP-tagged protein and the right image shows the target structure segmentation. Scale bars are 3 μm. These images demonstrate the validity and degree of spatial accuracy of the structure segmentations used for analysis. Several of these structure segmentations have specific types of caveats that may affect interpretation of downstream analysis in an analysis-specific manner (see Methods). These caveats are further described in (B-D). **B**) The limits of the cell boundary segmentation algorithm include potential errors of boundary detection for the very top slices of the cell. Desmosomes (via desmoplakin), which localize to the cell periphery at the top of the cell, demonstrate this caveat well. This can be seen in four images of sequential z-slices moving upwards towards the top of the cell membrane dye signal (z=42-48). The left, center, and right columns show the cell membrane dye, FP-tagged desmosomes (via desmoplakin), and desmosome segmentation images, respectively. Scale bar is 3 μm. In slice z=42, the cell membrane dye clearly identifies the true cell boundary (yellow arrows), which are also correctly detected by the cell segmentation. In z=48, the cell membrane dye, in focus desmosomes, and cell segmentation also all line up well, confirming the true cell boundary. However, at z=46 the cell membrane dye indicates two possible cell boundaries due to the slanted nature of the top of this cell and the out of focus light spreading from slices above and below that slice (yellow arrows and cyan arrows). Here, it is the location of the in focus desmosomes that identifies the inner possible cell boundary as the true cell boundary (yellow arrows). However, the segmentation detects the incorrect cell boundary (cyan arrows). Locating the cell boundary in z=44 is similar to z=46, but in this case, the desmosomes are not yet in focus, thus the true cell boundary is likely somewhere between that determined in z=42 and z=46. This error has a minor effect on the overall segmentation of the cell but a significant effect on the assignment of desmosome location in the cell. In this specific example, desmosomes are not located directly at the cell periphery but still close by such that their contribution to the total volume of desmosomes in this cell is still maintained. However, it is equally likely that that desmosomes (in this case), or any other structure localizing to the cell periphery at the top of the cell could be miss-assigned to a neighboring cell and thus these structures were not validated for cellular structure volume analyses (see Methods). **C)** Structures localizing or partially localizing to a thin 3D surface (such as the cell or nuclear periphery) may suffer from non-uniform accuracy between the middle and the top/bottom of that structure due to the anisotropic resolution of the images. Nuclear pores (via Nup153), which localize to the nuclear periphery demonstrate this caveat well. Here, seven sequential z-slice images of nuclear pores on the nuclear surface are shown (left column) along with their target structure segmentations (right column). Scale bar is 3 μm. The density of segmented nuclear pores is greatest in the first panel (z=36) and declines as the imaging plane moves upward through the nucleus. Consistently accurate detection for nuclear pores at both the center and the top of the nucleus was not possible due to this effect and would require further algorithm development. The accuracy of this target structure segmentation was sufficient to identify the general location of nuclear pores in the cell for the location-based analyses but not sufficient to be validated for use in the cellular structure volume analysis. This nuclear periphery caveat was also observed for other structures localizing to the nuclear and cell periphery (see Methods). **D)** The segmentation target for cohesins (via SMC-1A) is to detect the most contrasted locations of cohesins in nuclei. This segmentation works well for nuclei in most of interphase (see example in (A)). However, SMC-1A moves from the cytoplasm back into the nucleus after mitosis. Therefore, depending on how far along a cell is in interphase affects the amount of tagged SMC-1A protein in the nucleus and also the detection and segmentation of these most contrasted SMC-1A signal locations. Tagged SMC-1A is shown in three examples (left panel within each pair of panels) along with the target structure segmentations (right panel within each pair of panels). Scale bar is 3 μm. The left most example shows a cell in early interphase with SMC-1A both in the cytoplasm and nucleus, but at low levels such that the target segmentation is quite sparse. The right example shows a nucleus well into interphase with the expected cohesin target segmentation as in (A). In the center is a nucleus that is in between, with moderate levels of SMC-1A and thus fewer cohesin locations segmented. **E)** Example demonstrating the cell membrane Training Assay concept. The top row shows the tagged cell membrane channel (via CAAX; left) and the cell membrane dye channel (right) images as single slices near the center of the z-stack. The second row shows corresponding single slice side views of the same z-stacks. The bottom row shows the result of the CAAX-based segmentation (left) and the filled version for the cell membrane dye-based segmentation performed on the dye image after training via the cell membrane Training Assay (right). Scale bar is 5 μm. The cell membrane at the top of cells is often very dim in the cell membrane dye channel images (yellow arrows) due to both the very thin nature of the top membrane and photobleaching during z-stack acquisition. This creates a significant challenge for a cell membrane dye-based segmentation. However, the top of these same cells is much more visible in the tagged plasma membrane cell line (via CAAX; cyan arrows), permitting successful CAAX-based segmentations. We leveraged the image information contained in the CAAX images by using the CAAX-based segmentation result as the training target for a deep learning cell membrane dye-based segmentation model. **F)** Example demonstrating the DNA dye Training Assay concept. The top row shows the tagged nuclear envelope channel (via lamin B1; left) and the DNA dye channel (right) images as a single slice near the center of the z-stack. The second row shows the corresponding single slice side views of the same z-stacks. The bottom row shows the result of the lamin B1-based segmentation (left) and the filled version for the DNA dye-based segmentation performed on the dye image after training via the DNA dye Training Assay (right). Scale bar is 5 μm. The top boundary of nuclei is often very blurry in the DNA dye channel images (yellow arrows) due to the “filled” nature of how the DNA dye demarcates the nuclear boundary combined with the diffraction of light and lower axial resolution. This creates a significant challenge for accurately identifying the top nuclear boundaries to define a segmentation target. However, the top boundaries of nuclei in these same cells are clearly identifiable in the tagged nuclear envelope cell line (via lamin B1; cyan arrows), permitting accurate identification and segmentation of nuclei via lamin B1. We leveraged the image information contained in the lamin B1 images by using the lamin B1-based segmentation result as the training target for a deep learning DNA dye-based segmentation model.

**Figure S2**

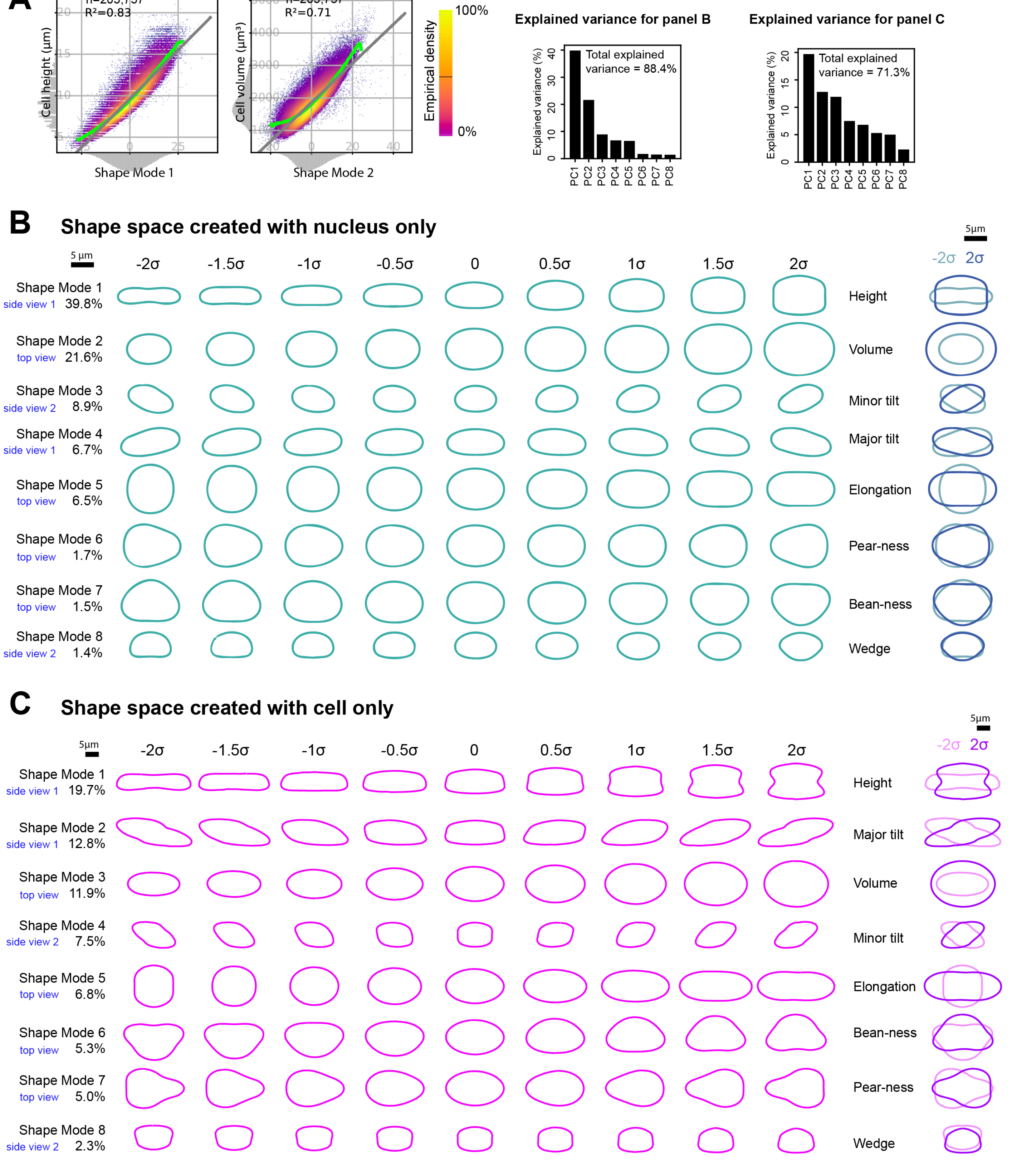

A
n=203,737
R²=0.83
Cell height (μm)
Shape Mode 1
n=203,737
R²=0.71
Cell volume (μm³)
Shape Mode 2
Empirical density
100%
0%
Explained variance for panel B
Total explained variance = 88.4%
Explained variance (%)
PC1 PC2 PC3 PC4 PC5 PC6 PC7 PC8
Explained variance for panel C
Total explained variance = 71.3%
Explained variance (%)
PC1 PC2 PC3 PC4 PC5 PC6 PC7 PC8
B
Shape space created with nucleus only
5 μm
-2σ -1.5σ -1σ -0.5σ 0 0.5σ 1σ 1.5σ 2σ
5μm
-2σ 2σ
Shape Mode 1 side view 1 39.8% Height
Shape Mode 2 top view 21.6% Volume
Shape Mode 3 side view 2 8.9% Minor tilt
Shape Mode 4 side view 1 6.7% Major tilt
Shape Mode 5 top view 6.5% Elongation
Shape Mode 6 top view 1.7% Pear-ness
Shape Mode 7 top view 1.5% Bean-ness
Shape Mode 8 side view 2 1.4% Wedge
C
Shape space created with cell only
5μm
-2σ -1.5σ -1σ -0.5σ 0 0.5σ 1σ 1.5σ 2σ
5μm
-2σ 2σ
Shape Mode 1 side view 1 19.7% Height
Shape Mode 2 side view 1 12.8% Major tilt
Shape Mode 3 top view 11.9% Volume
Shape Mode 4 side view 2 7.5% Minor tilt
Shape Mode 5 top view 6.8% Elongation
Shape Mode 6 top view 5.3% Bean-ness
Shape Mode 7 top view 5.0% Pear-ness
Shape Mode 8 side view 2 2.3% Wedge

**Figure S2.** Supplemental figure for **Figure 2**. **A)** Cell height as a function of Shape Mode 1 (left graph), and cell volume as a function of Shape Mode 2 (right graph). Each point represents a single cell (n=203,737). Points are color-coded based on an empirical density estimate. The gray line represents the best linear fit and the green curve represents the rolling average. **B and C)** The nuclear (B) and cell (C) shape spaces obtained when only nuclear (B) and cell (C) SHE coefficients are used as input for the PCA dimensionality reduction described in **Figure 2,** respectively. The top bar graph plots the total variance explained by each principal component. The bottom panel shows 2D projections of 3D meshes obtained for each of the nine map points of each of the eight shape modes. At the far right is an overlay of nuclear mesh projections for the two most extremes map points (at -2σ, lighter shade and +2σ, darker shade) of each shape mode. All three views for each shape mode and map point can be seen in **Movies S3&4**.

**Figure S3**

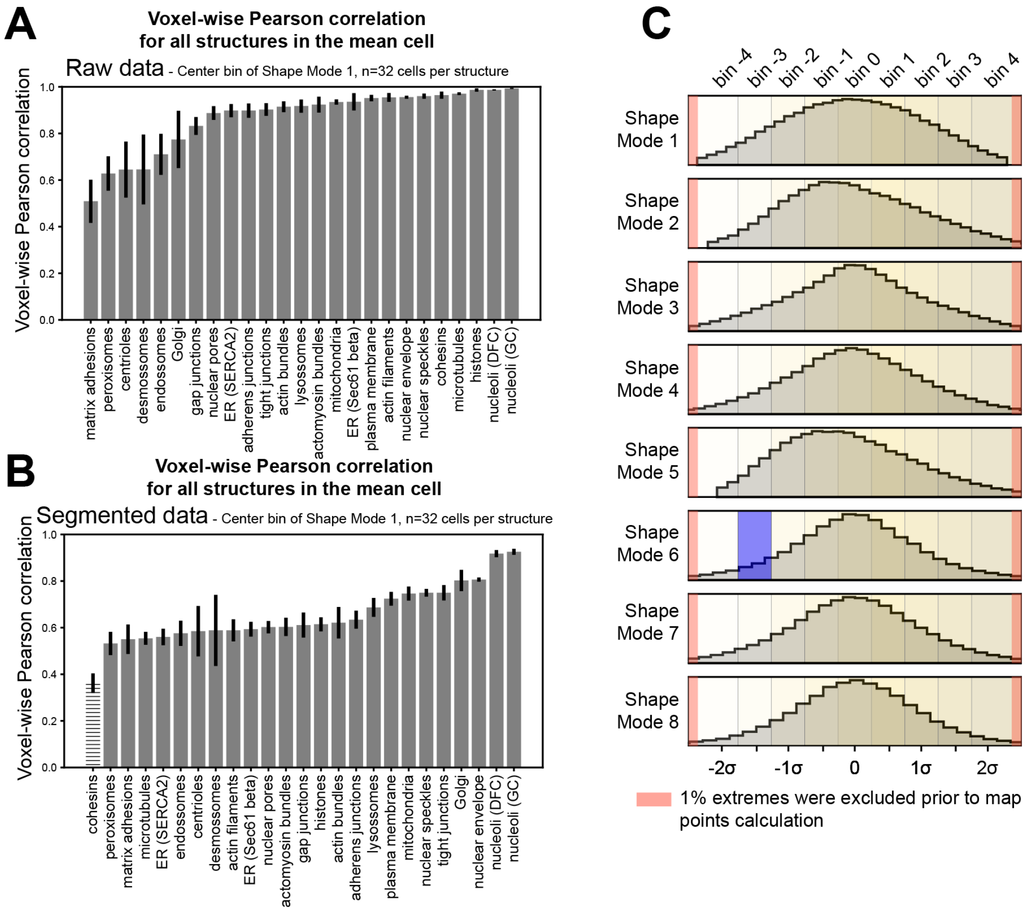
A
Voxel-wise Pearson correlation
for all structures in the mean cell
Raw data - Center bin of Shape Mode 1, n=32 cells per structure
Voxel-wise Pearson correlation
matrix adhesions
peroxisomes
centrioles
desmossomes
endossomes
Golgi
gap junctions
nuclear pores
ER (SERCA2)
adherens junctions
tight junctions
actin bundles
lysossomes
actomyosin bundles
mitochondria
ER (Sec61 beta)
plasma membrane
actin filaments
nuclear envelope
nuclear speckles
cohesins
microtubules
histones
nucleoli (DFC)
nucleoli (GC)
B
Voxel-wise Pearson correlation
for all structures in the mean cell
Segmented data - Center bin of Shape Mode 1, n=32 cells per structure
Voxel-wise Pearson correlation
cohesins
peroxisomes
matrix adhesions
microtubules
ER (SERCA2)
endossomes
centrioles
desmossomes
actin filaments
ER (Sec61 beta)
nuclear pores
actomyosin bundles
gap junctions
histones
actin bundles
adherens junctions
lysossomes
plasma membrane
mitochondria
nuclear speckles
tight junctions
Golgi
nuclear envelope
nucleoli (DFC)
nucleoli (GC)
C
bin -4
bin -3
bin -2
bin -1
bin 0
bin 1
bin 2
bin 3
bin 4
Shape Mode 1
Shape Mode 2
Shape Mode 3
Shape Mode 4
Shape Mode 5
Shape Mode 6
Shape Mode 7
Shape Mode 8
-2σ
-1σ
0
1σ
2σ
1% extremes were excluded prior to map points calculation

Figure S3 continued

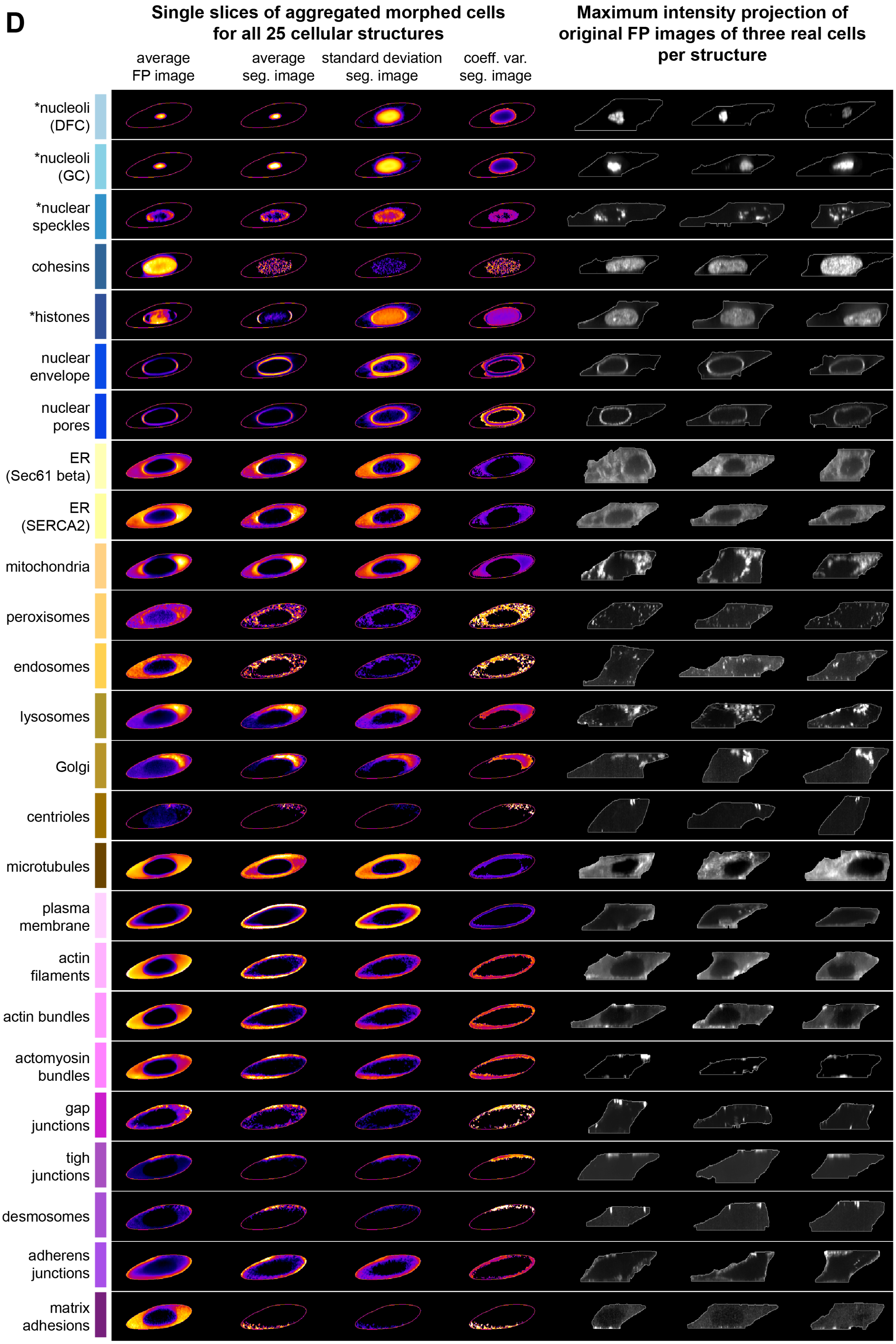

**Figure S3.** Supplemental figure for **Figure 3. A)** Voxel-wise Pearson correlation between original intensity images of FP-tagged proteins and the image reconstructed from the parameterized intensity representation. Error bars represent ± std around the mean (n=32 cells per structure). Cells were selected from center bin of Shape Mode 1. **B)** Same as in (A) for segmented images instead of original intensity images. The correlation for cohesins (via SMC-1A) is indicated with a striped fill pattern. This structure has a significantly changing target structure segmentation depending on how much tagged cohesin has re-entered the nucleus after mitosis, causing the much lower correlation value (**Figure S1 D**). Error bars represent ± std around the mean (n=32 cells per structure). **C)** Probability density distribution of principal component values. Cells that fall into the ranges from 0-1$^{st}$ percentile and from 99$^{th}$ to 100$^{th}$ percentile of each principal component were not used for principal components interpretation. These ranges are shown in red. Remaining data is normalized to units of standard deviation and binned into 9 equal size bins around the map points -2σ, -1.5σ, -1.0σ, -0.5σ, 0, 0.5σ, 1.0σ, 1.5σ and 2.0σ. As an example, the bin corresponding to the map point (0,0,0,0,0,-1.5σ,0,0) is highlighted in blue. **D)** The first four columns show side view 1 of four different types of aggregated morphed cells at the Shape Mode 3 map point (0,0,1.5σ,0,0,0,0,0) for all 25 cellular structures. The first two columns are average morphed cell based on the original FP images and the target structure segmentations, respectively. The next two columns are the standard deviation and structure-localized coefficient of variation morphed cells based on the target structure segmentations. The last three columns show three examples per cellular structure of a partial maximum intensity projection of the original FP images of real cells that are located within this map point bin. Maximum intensity projections were performed either on the center 25 slices or center 13 slices (marked with an asterisk) the cells. Brightness and contrast were adjusted per cell to best represent each FP-tagged cellular structure.

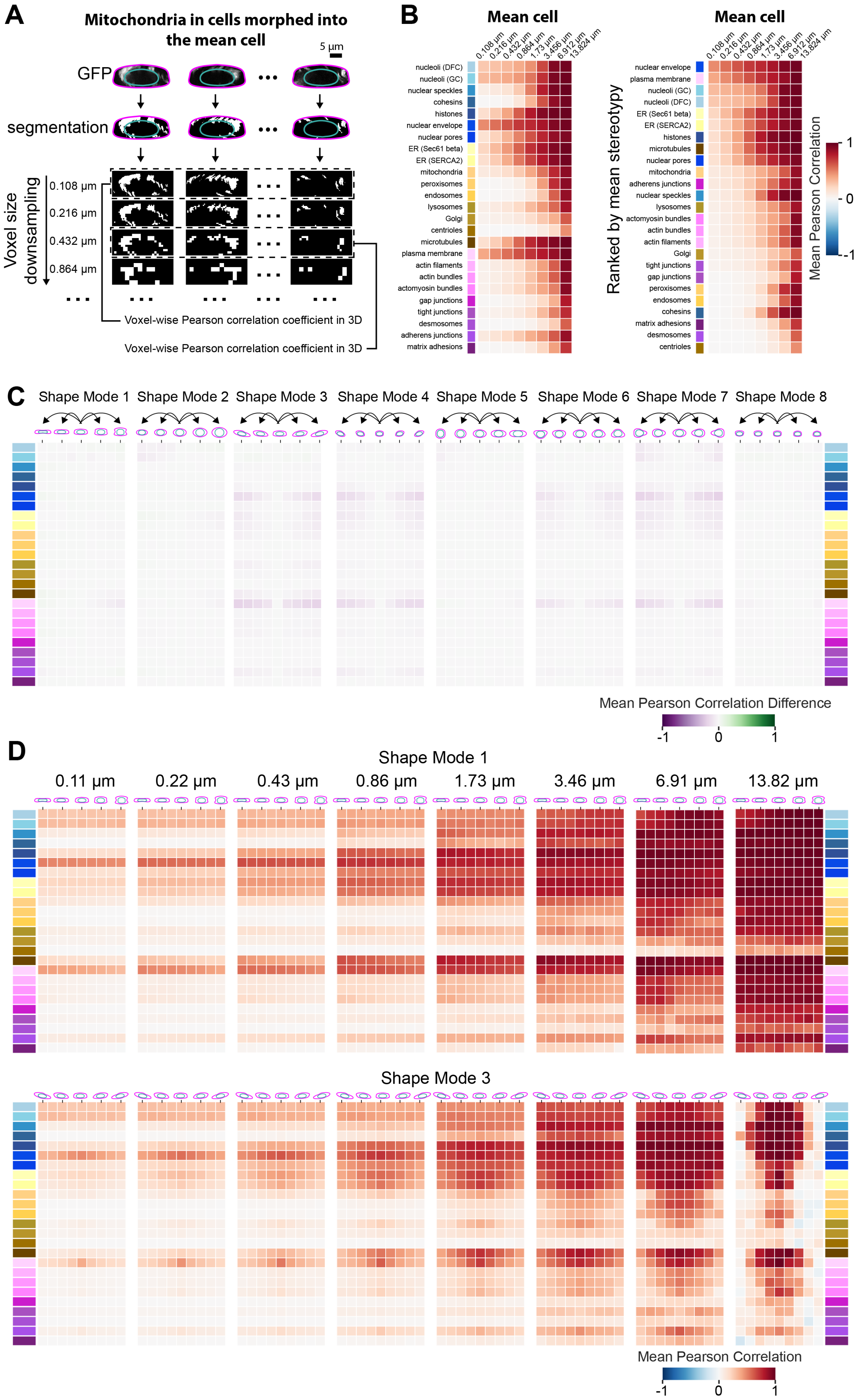
A
Mitochondria in cells morphed into the mean cell
5 µm
GFP
segmentation
Voxel size downsampling
0.108 µm
0.216 µm
0.432 µm
0.864 µm
Voxel-wise Pearson correlation coefficient in 3D
Voxel-wise Pearson correlation coefficient in 3D
B
Mean cell
0.108 µm
0.216 µm
0.432 µm
0.864 µm
1.73 µm
3.456 µm
6.912 µm
13.824 µm
nucleoli (DFC)
nucleoli (GC)
nuclear speckles
cohesins
histones
nuclear envelope
nuclear pores
ER (Sec61 beta)
ER (SERCA2)
mitochondria
peroxisomes
endosomes
lysosomes
Golgi
centrioles
microtubules
plasma membrane
actin filaments
actin bundles
actomyosin bundles
gap junctions
tight junctions
desmosomes
adherens junctions
matrix adhesions
Mean cell
Ranked by mean stereotypy
nuclear envelope
plasma membrane
nucleoli (GC)
nucleoli (DFC)
ER (Sec61 beta)
ER (SERCA2)
histones
microtubules
nuclear pores
mitochondria
adherens junctions
nuclear speckles
lysosomes
actomyosin bundles
actin bundles
actin filaments
Golgi
tight junctions
gap junctions
peroxisomes
endosomes
cohesins
matrix adhesions
desmosomes
centrioles
Mean Pearson Correlation
1
0
-1
C
Shape Mode 1
Shape Mode 2
Shape Mode 3
Shape Mode 4
Shape Mode 5
Shape Mode 6
Shape Mode 7
Shape Mode 8
Mean Pearson Correlation Difference
-1
0
1
D
Shape Mode 1
0.11 µm
0.22 µm
0.43 µm
0.86 µm
1.73 µm
3.46 µm
6.91 µm
13.82 µm
Shape Mode 3
Mean Pearson Correlation
-1
0
1

**Figure S4.** Supplemental figure for **Figure 4**. **A)** The segmented images of mitochondria in cells morphed into the mean cell shape were down sampled via iterative array reduction using a block size of (2,2,2) until the image reaches a voxel size of 1x1x1. The voxel-wise Pearson correlation was calculated for every pair of morphed cells at each down sampled resolution and results were organized as a correlation matrix. **B)** Heatmaps of Pearson correlations showing the variation of mean stereotypy across down sampled resolutions for the mean cell shape. Heatmaps were either ranked by structure compartment order (left) as in **Figure 4B** or ranked by their mean stereotypy at the original voxel size (right). Each column represents a particular down sampled voxel size. **C)** Each heatmap value corresponds to the mean stereotypy difference between the cell shape in the indicated shape mode bin and the mean cell shape. Each heatmap represents a shape mode and each column of the heatmaps represent the different map points within that shape mode. **D)** Heatmaps are regenerated for shape modes 1 (top) and 3 (bottom) with successive voxel-size down sampling. Each heatmap corresponds to one voxel size and each column in the heatmap corresponds to one shape mode map point. Colors on the left and right of panels (C) and (D) indicate the same cellular structures as represented in panel (B). N = 300 morphed cells for each cellular structure and shape mode bin or the maximum number of cells available (see **DataFile S1** and Methods).

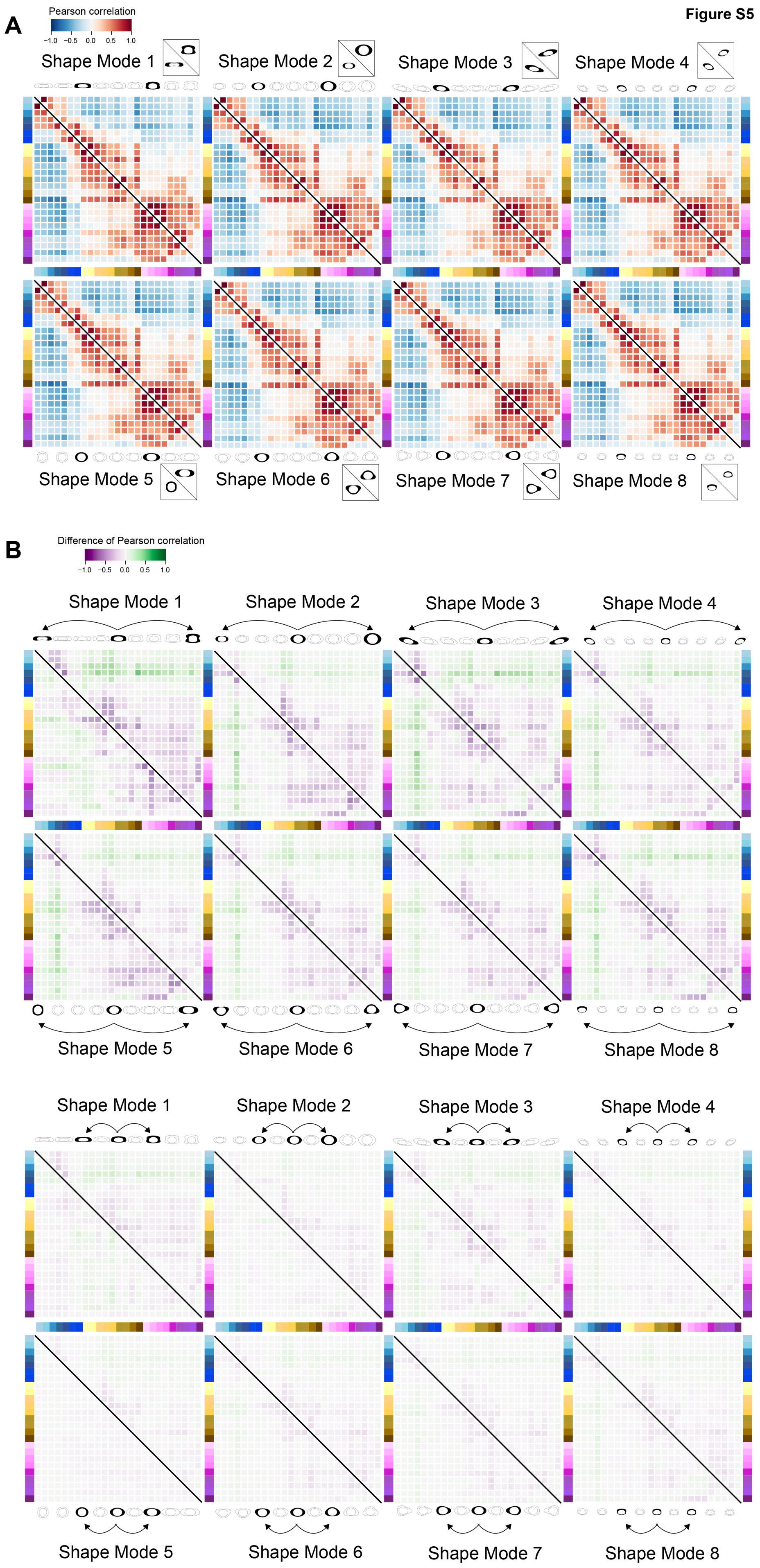
Figure S5
A
Pearson correlation
−1.0 −0.5 0.0 0.5 1.0
Shape Mode 1
Shape Mode 2
Shape Mode 3
Shape Mode 4
Shape Mode 5
Shape Mode 6
Shape Mode 7
Shape Mode 8
B
Difference of Pearson correlation
−1.0 −0.5 0.0 0.5 1.0
Shape Mode 1
Shape Mode 2
Shape Mode 3
Shape Mode 4
Shape Mode 5
Shape Mode 6
Shape Mode 7
Shape Mode 8
Shape Mode 1
Shape Mode 2
Shape Mode 3
Shape Mode 4
Shape Mode 5
Shape Mode 6
Shape Mode 7
Shape Mode 8

C

Figure S5 continued

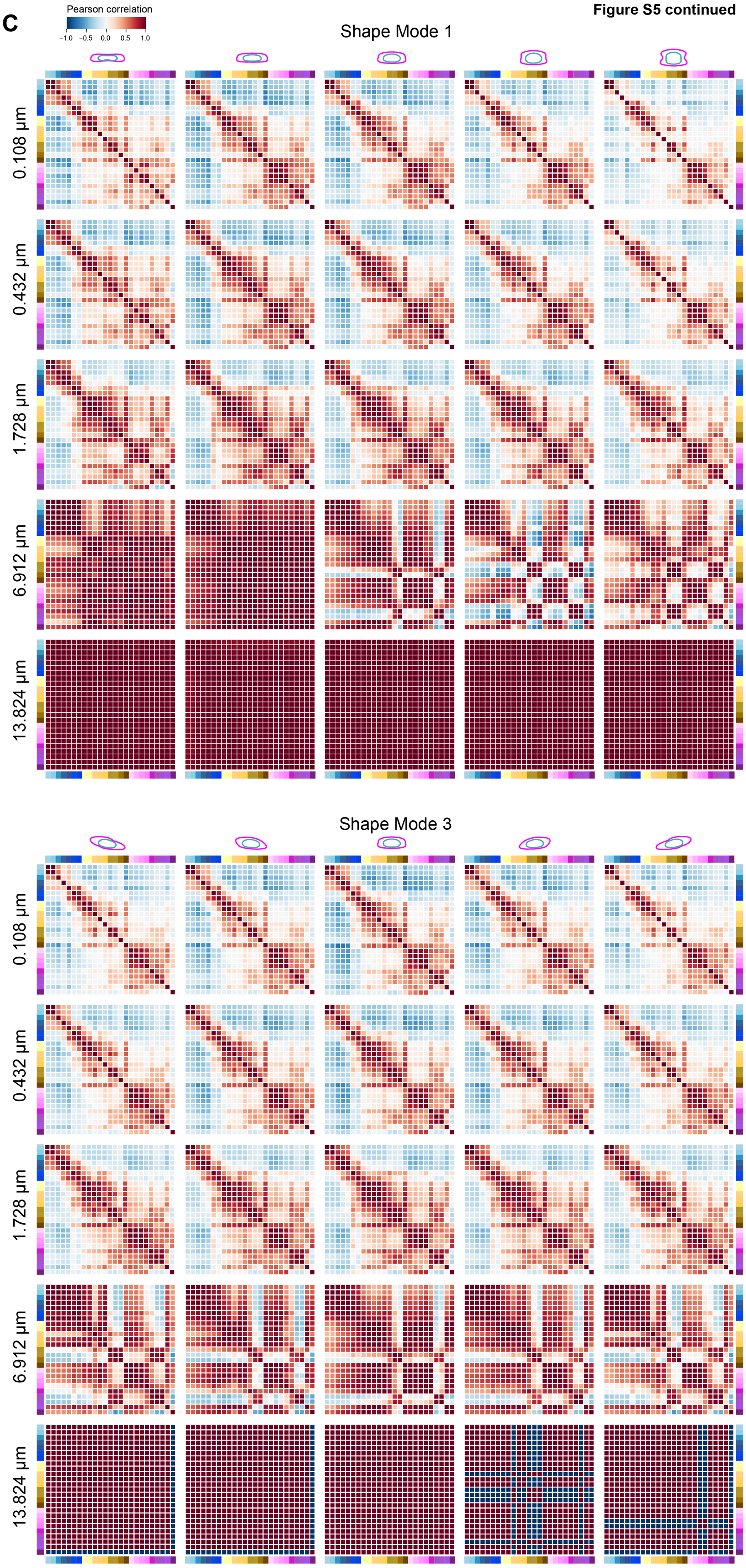

**Figure S5.** Supplemental figure for **Figure 5**. **A)** The process to create the concordance heatmap for the mean cell shape in **Figure 5B** was repeated for the reconstructed cell and nuclear shapes at the -1σ and 1σ shape space map points for each of the 8 shape modes. Each heatmap represents one shape mode. The lower triangle represents shape space map point -1σ and the upper triangle represents shape space map point 1σ. For sake of clarity, diagonals are colored in white and black lines are used to separate the lower and upper triangles. **B)** Each heatmap value corresponds to the difference in concordance between the cell shape in the indicated shape mode bin and the mean cell shape. The top panel of eight heatmaps corresponds to the -2σ and 2σ map points and the bottom panel to the -1σ and 1σ map points. **C)** Concordance heatmaps for five shape mode bins (columns) for Shape Mode 1 (top panel of 25 heatmaps) and for Shape Mode 3 (bottom panel of 25 heatmaps) with successive downsampling of voxel size (rows). The number of cells analyzed for each cellular structure and shape mode bin can be found in **DataFile S1**.

**Figure S6**

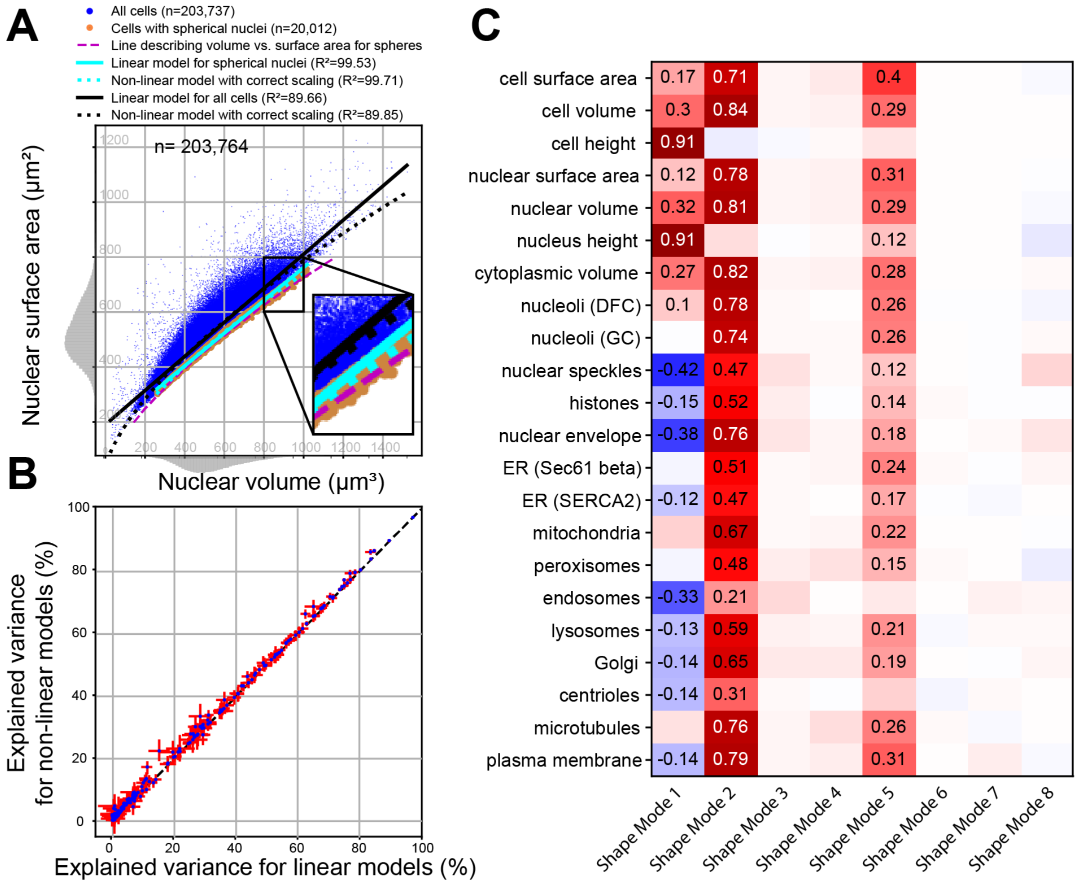
A
All cells (n=203,737)
Cells with spherical nuclei (n=20,012)
Line describing volume vs. surface area for spheres
Linear model for spherical nuclei (R²=99.53)
Non-linear model with correct scaling (R²=99.71)
Linear model for all cells (R²=89.66)
Non-linear model with correct scaling (R²=89.85)
n= 203,764
Nuclear surface area (μm²)
Nuclear volume (μm³)
B
Explained variance for non-linear models (%)
Explained variance for linear models (%)
C
cell surface area 0.17 0.71 0.4
cell volume 0.3 0.84 0.29
cell height 0.91
nuclear surface area 0.12 0.78 0.31
nuclear volume 0.32 0.81 0.29
nucleus height 0.91 0.12
cytoplasmic volume 0.27 0.82 0.28
nucleoli (DFC) 0.1 0.78 0.26
nucleoli (GC) 0.74 0.26
nuclear speckles -0.42 0.47 0.12
histones -0.15 0.52 0.14
nuclear envelope -0.38 0.76 0.18
ER (Sec61 beta) 0.51 0.24
ER (SERCA2) -0.12 0.47 0.17
mitochondria 0.67 0.22
peroxisomes 0.48 0.15
endosomes -0.33 0.21
lysosomes -0.13 0.59 0.21
Golgi -0.14 0.65 0.19
centrioles -0.14 0.31
microtubules 0.76 0.26
plasma membrane -0.14 0.79 0.31
Shape Mode 1
Shape Mode 2
Shape Mode 3
Shape Mode 4
Shape Mode 5
Shape Mode 6
Shape Mode 7
Shape Mode 8

**Figure S6.** Supplemental figure for **Figure6**. **A)** Linear fit between nuclear volume and nuclear surface area. The volume (V) and surface area (A) of a sphere don't scale linearly, instead $A \sim V^{2/3}$. However, on this dataset a linear model of nuclear volume explains as much variation in nuclear area as a model with the theoretically correct non-linear scaling factor. The scatter plot depicts the nuclear volume (x-axis) and nuclear surface area (y-axis) for all cells in the dataset (n=203,737; blue points). We selected a set of n=20,012 cells with nuclei that are approximately spherical. These cells were defined as those falling within the lowest $10^{th}$percentile for nuclear surface area after binning the cells in 250 equally spaced bins in the nuclear volume range from 254 $\mu m^3$ to 1,017 $\mu m^3$. (light brown points). We verified that these cells were spherical in two ways: First, we created 3D images of perfect spheres with different radii and calculated the volume and surface area using the same method (and code) that was used to calculate the volumes and areas of the cells and nuclei. This yielded the dashed magenta line, which visually intersects with the light brown colored cells. We also visually inspected and confirmed that cells in the light brown set were close to spherical. A linear model (y=ax+b) that was fit on the cells with spherical nuclei explained 99.53% of the variation in nuclear area (solid cyan line). A non-linear, theoretically correct model, $y=ax^{2/3}+b$, explained 99.71%; i.e. only a fraction more (dashed cyan line). Similarly, a linear model that was fit on all cells explained 89.66% (solid black line), whereas the $y=ax^{2/3}+b$ non-linear model explained 89.85% of the variation, once again only a fraction more (dashed black line). The notion that a linear model provides a good fit even for relationships known to be non-linear is not unexpected (and has frequently been observed before; (Marshall, 2020), as this non-linear relationship is not strongly manifested on the relatively small range of observed nuclear volumes. Moreover, as indicated by the bottom and side histograms, the large majority of cells are concentrated in an even smaller range. **B)** Non-linear models do not increase explained variance compared to linear models. We compared the explained variance of linear models to non-linear models across all 190 cases reported in the heatmaps of **Figure 6A**, and found that non-linear models cannot explain more variation than the linear models. The scatterplot depicts the explained variance for linear models (x-axis) and non-linear models (y-axis) for all (n=190) cases originally reported in the Heatmap part 1 of **Figure 6A**. The blue points represent the median explained variance across 100 bootstraps of the regression model. The red lines indicate the 95% confidence interval (from 2.5% to 97.5% across the 100 bootstraps). The non-linear models were calculated as described in the Methods. Since these non-linear models are more expressive and the explained variance is computed on the fitted data (and not an external validation set) the non-linear models, per definition, explain the same or more variation in the data. However, the differences are minimal. Specifically, out of the 190 cases, there are only ten cases in which the 95% confidence intervals between the linear and non-linear model do not overlap and the non-linear model explains more than 1% more variance than the linear model. There are only two cases in which the non-linear model explains more than 5% more variance than the linear model. Both these cases involve the endosomes, the structure with the lowest explained variance overall. The multivariate non-linear model that includes the four cell and nuclear size metrics explained 22.5% of endosome volume, whereas the linear model explained 15.0%. Similarly, the unique variance explained by "nuc a+v" (the nuclear area and volume; **Figure 6A**) in the non-linear case is 17.4%, and 11.2% in the linear case. **C)** Size scaling metrics only correlate with Shape Modes 1, 2 and 5. This heatmap depicts Pearson correlation scores between size scaling metrics (rows) and shape modes (sm, columns). Absolute Pearson correlation scores below 0.1 are not printed. For the seven cell and nuclear metrics (the seven top rows) all cells (n=203,737) were used in the correlation calculations. For the structure volumes (15 bottom rows) only those cells that correspond to the structure in question were used (**Table S1**) Correlations of structure volume to Shape Mode 5 likely occur due the moderate correlation between Shape Mode 5 (elongation) and cell surface area.

**Figure S7**

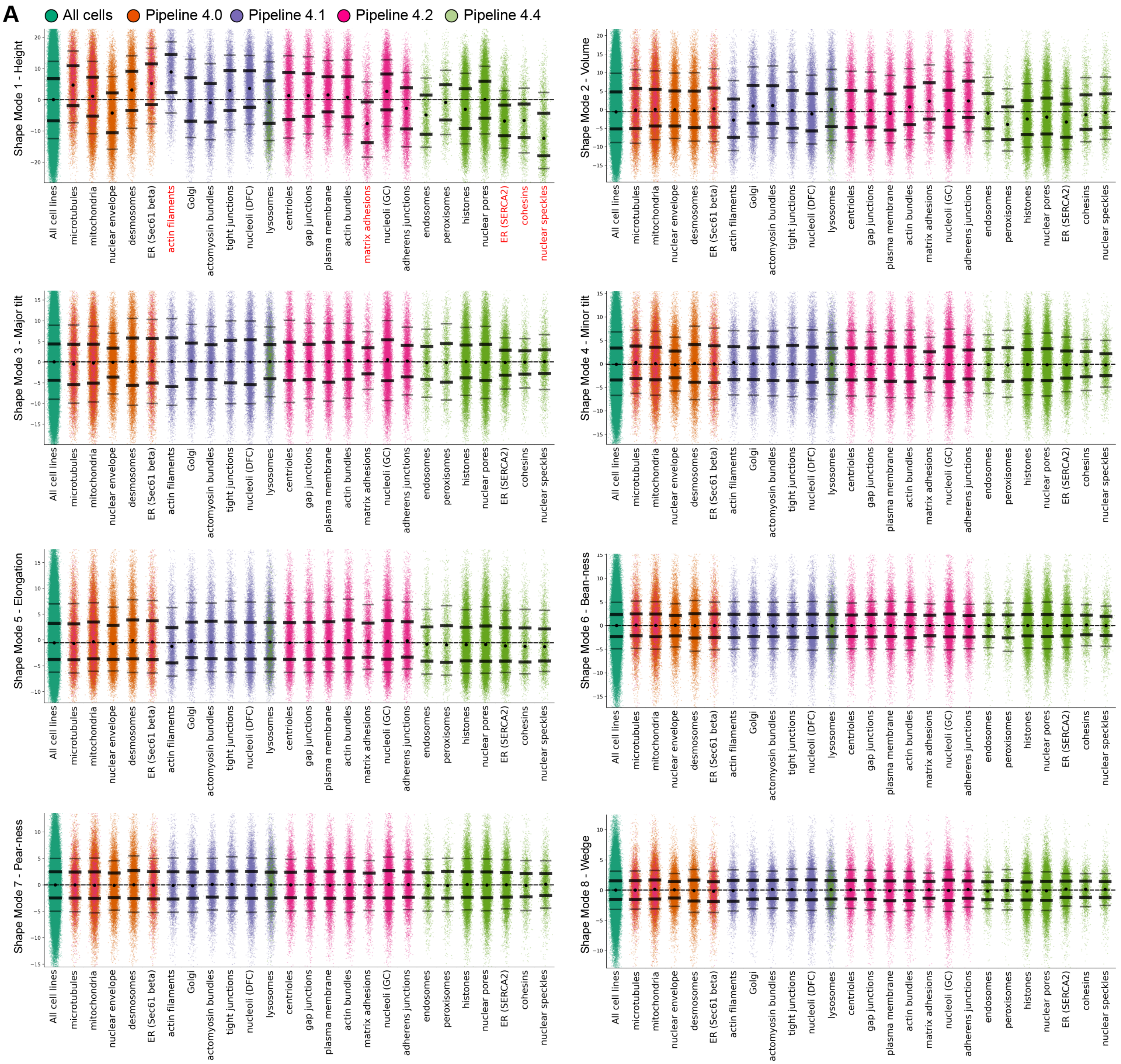
A
All cells
Pipeline 4.0
Pipeline 4.1
Pipeline 4.2
Pipeline 4.4
Shape Mode 1 - Height
Shape Mode 2 - Volume
Shape Mode 3 - Major tilt
Shape Mode 4 - Minor tilt
Shape Mode 5 - Elongation
Shape Mode 6 - Bean-ness
Shape Mode 7 - Pear-ness
Shape Mode 8 - Wedge
All cell lines
microtubules
mitochondria
nuclear envelope
desmosomes
ER (Sec61 beta)
actin filaments
Golgi
actomyosin bundles
tight junctions
nucleoli (DFC)
lysosomes
centrioles
gap junctions
plasma membrane
actin bundles
matrix adhesions
nucleoli (GC)
adherens junctions
endosomes
peroxisomes
histones
nuclear pores
ER (SERCA2)
cohesins
nuclear speckles

### Figure S7 continued

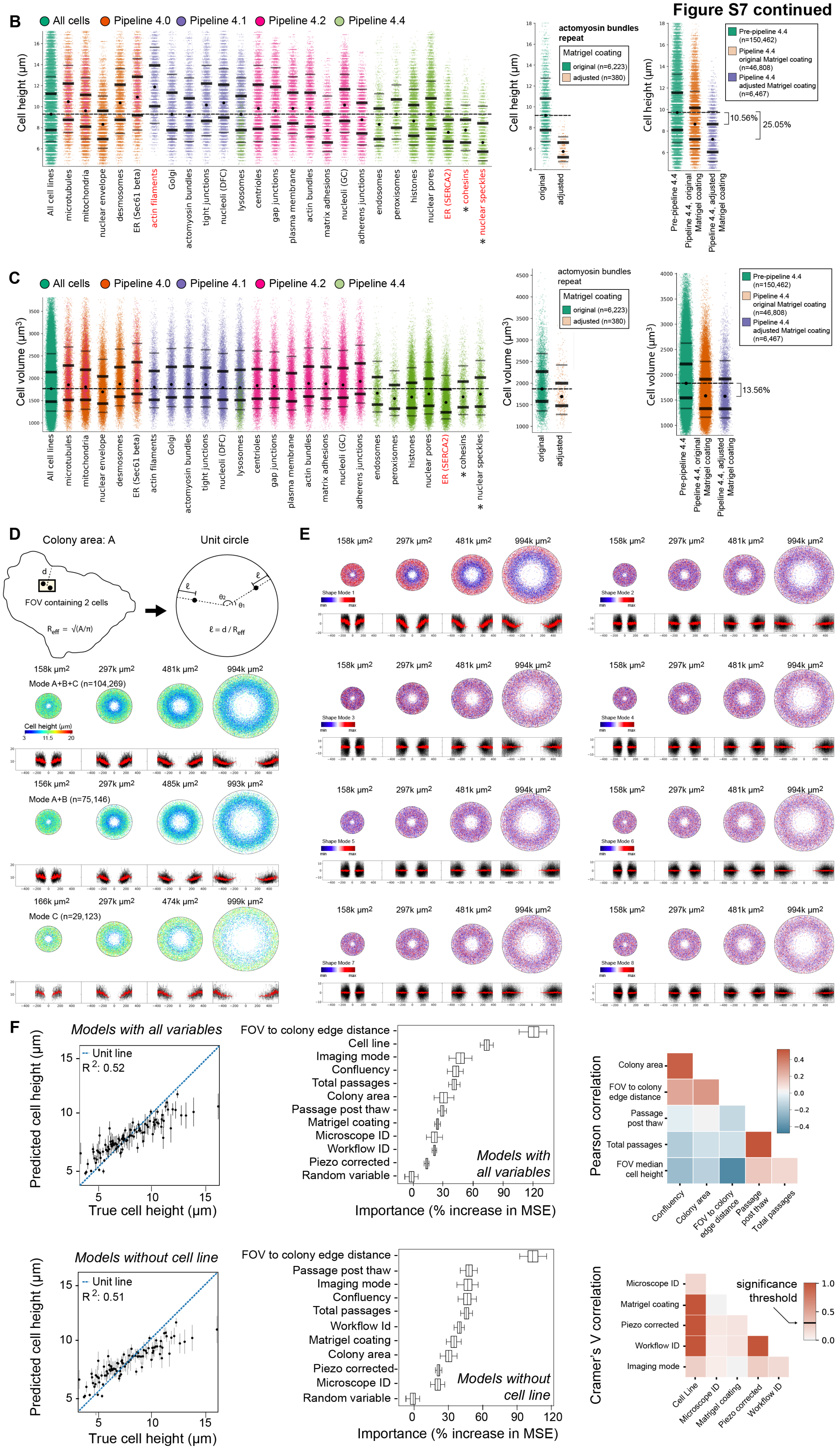

**Figure S7.** Statistical analysis for quality control of the hiPSC Single-Cell Image Dataset. **A)** Distributions of principal component values for all cell lines together (first bin in dark green) and per tagged structure cell line, plotted in pipeline timeline order, the order that structure datasets were collected (total n=203,737). The dashed horizontal line represents the median value for all cell lines together (first bin in dark green). The black dots in each structure bin represent the median of the distribution for all cells from the corresponding cell line. Thick and thin solid tick lines plotted for each structure bin represent the interquartile range (IQR; 25% above and below the median) and the 10$^{th}$ and 90$^{th}$ percentiles of the distribution. The colors for each cell line refer to the pipeline workflow (see Methods for details). Structure names along the bottom are red if the interquartile range does not overlap with the mean value for all cell lines. **B)** Left plot shows the distributions of cell height for all cell lines together (first bin in dark green) and per tagged structure cell line, plotted in pipeline timeline order (same n and plot format as in **A**). The asterisks next to the structure names indicate those structures imaged with an adjusted Matrigel coating protocol towards the end of the pipeline timeline. The center plot shows a comparison of cell height between actomyosin bundle-tagged cells (via non-muscle myosin IIB) in the main dataset (Pipeline 4.1; n=6,223) and in a repeat dataset imaged with Pipeline 4.4 settings with the adjusted Matrigel coating protocol (n=380). The right plot shows a comparison of cell height between all cell lines imaged pre-Pipeline 4.4, during Pipeline 4.4 with original Matrigel coating and during Pipeline 4.4 with adjusted Matrigel coating. Percentages shown in the plot are the relative height reduction compared to the mean height of cell lines imaged pre-Pipeline 4.4. **C)** The left, center, and right plots are the same as the top row in B except for cell volume instead of cell height. **D)** The top image diagrams circular mapping of imaged colonies (via the 12X overview images). Two cells are represented by two black dots within an FOV, represented by a rectangle. The FOV center is at distance *d* from the closest edge of the colony. The two cells are then mapped into a unit circle that serves as a template to visualize the radial location of the two cells. The radial location is the FOV relative distance to the edge of the colony, $\ell = d/R_{eff}$, where $R_{eff}$ represents the effective radius of the colony. The angular location of a cell ($\theta_1$ and $\theta_2$ for the two cells in the image) is independently drawn from a uniform distribution of angles in the range [0,2π]. Cells from the dataset that were associated with a colony size (see Methods) were grouped into four bins, each with similar number of cells, based on the area of the colony where they came from. The colony area range of each bin is 15k-230k µm$^2$, 230k-377k µm$^2$, 377-620k µm$^2$ and 620k-14,285k µm$^2$. Each point represents one cell within the colony area bin that was mapped into the unit circle. The unit circle was then rescaled to match the mean colony area for that bin. Points are color coded by their corresponding cell height. Listed above each circle is the mean colony area in that bin to which the unit circle is scaled. Below each circle are profile plots of cell height as a function of the radial distance for each of the cell (in black). The red curve represents the rolling average. Each row of circular colony mappings represents a different aggregation of the data based on the imaging mode: the first row is for all imaging modes (modes A, B and C; n=104,269), the second row is for modes A and B only (n=75,146) and third row is for mode C only (n=29,123). **E)** Circular colony mappings as in (D) where points (cells) are now color coded by values of shape modes 1 to 8 (all imaging modes, n=104,269). **F)** Scatter plots on the far left show true values of cell height compared to cell height values predicted by random forest regression models (see Methods) that include either all experimental variables (top plot) or all experimental variables except for the cell line identity (bottom plot). The error bars on the predicted values are obtained via bootstrapping. The center column shows box plots representing the feature importance for each of the two models as measured by the increase in the mean squared error (MSE) when all values of that corresponding feature are shuffled across samples. Box plots indicate the median, $Q_1$-0.95xIQR, IQR and $Q_3$+0.95xIQR. The right top plot is the Pearson correlation matrix between five continuous experimental variables used in training the regression models. The bottom right plot is the Cramer's V correlation matrix between six categorical experimental variables used in training the regression models. Variables with correlation above the significance threshold 0.3 are assumed to be highly correlated (McHugh, 2013).

**Table S1:** Number of cells for each cellular structure in the hiPSC Single-Cell Image Dataset

| Workflow | Acquisition Order | Structure | Single-Cell Image Dataset | Mitotic Cells | Interphase Cells | Outliers Cells (Interphase) | Analyzed Dataset (Interphase) |
|---|---|---|---|---|---|---|---|
| Pipeline 4.0 | 1 | microtubule | 9692 | 538 | 9154 | 28 | 9126 |
| | 2 | mitochondria | 24426 | 1292 | 23134 | 78 | 23056 |
| | 3 | nuclear envelope | 12409 | 514 | 11895 | 36 | 11859 |
| | 4 | desmosomes | 10583 | 430 | 10153 | 354 | 9799 |
| | 5 | ER (Sec61 beta) | 6714 | 296 | 6418 | 9 | 6409 |
| Pipeline 4.1 | 6 | actin bundles | 4010 | 186 | 3824 | 0 | 3824 |
| | 7 | Golgi | 6498 | 309 | 6189 | 13 | 6176 |
| | 8 | actomyosin bundles | 6596 | 373 | 6223 | 14 | 6209 |
| | 9 | tight junctions | 5881 | 294 | 5587 | 40 | 5547 |
| | 10 | nucleoli (DFC) | 10446 | 460 | 9986 | 38 | 9948 |
| | 11 | lysosomes | 10745 | 604 | 10141 | 27 | 10114 |
| Pipeline 4.2 | 12 | centrioles | 7780 | 452 | 7328 | 210 | 7118 |
| | 13 | gap junctions | 6548 | 373 | 6175 | 2 | 6173 |
| | 14 | plasma membrane | 8107 | 436 | 7671 | 18 | 7653 |
| | 15 | actin filaments | 8224 | 561 | 7663 | 5 | 7658 |
| | 16 | matrix adhesions | 3880 | 306 | 3574 | 86 | 3488 |
| | 17 | nucleoli (GC) | 12550 | 685 | 11865 | 48 | 11817 |
| | 18 | adherens junctions | 6223 | 374 | 5849 | 6 | 5843 |
| Pipeline 4.4 | 19 | endosomes | 2605 | 190 | 2415 | 4 | 2411 |
| | 20 | peroxisomes | 1997 | 144 | 1853 | 0 | 1853 |
| | 21 | histones | 15877 | 784 | 15093 | 25 | 15068 |
| | 22 | nuclear pores | 18719 | 932 | 17787 | 15 | 17772 |
| | 23 | ER (SERCA2) | 10177 | 457 | 9720 | 16 | 9704 |
| | 24 | cohesins | 2392 | 105 | 2287 | 12 | 2275 |
| | 25 | nuclear speckles | 2983 | 143 | 2840 | 3 | 2837 |
| | | Total | 216062 | 11238 | 204824 | 1087 | 203737 |

**Supplemental Movie Captions:**

**Movie S1:** Animated gif of the cell and nuclear shape space shown in **Figure 2E**. 2D projections of 3D meshes obtained for each of the nine map point bins of each of the eight shape modes. All three views are shown for each mode, as indicated along the top. Human-interpretable names for these shape modes are indicated on the right. Mesh projections of the cell are in magenta and of the nucleus are in cyan. Each frame of the movie shows a successive map point location along each shape mode, from $-2\sigma$ to $2\sigma$ in steps of $0.5\sigma$ ($\sigma$ = standard deviation).

**Movie S2:** 3D visualization of eleven structures rendered simultaneously to illustrate their relative spatial relationships as in **Figure 3G**.

**Movie S3:** Animated gif of the "*cell-only*" shape space shown in **Figure S2B**. 2D projections of 3D meshes obtained for each of the nine map point bins of each of the eight shape modes. All three views are shown for each mode, as indicated along the top. Human-interpretable names for these shape modes are indicated on the right. Mesh projections of the cell are in magenta. Each frame of the movie shows a successive map point location along each shape mode, from $-2\sigma$ to $2\sigma$ in steps of $0.5\sigma$ ($\sigma$ = standard deviation).

**Movie S4:** Animated gif of the "*nucleus-only*" shape space shown in **Figure S2B**. 2D projections of 3D meshes obtained for each of the nine map point bins of each of the eight shape modes. All three views are shown for each mode, as indicated along the top. Human-interpretable names for these shape modes are indicated on the right. Mesh projections of the nucleus are in cyan. Each frame of the movie shows a successive map point location along each shape mode, from $-2\sigma$ to $2\sigma$ in steps of $0.5\sigma$ ($\sigma$ = standard deviation).
