## Supplementary figures and images for "Robust integrated intracellular organization of the human iPS cell: where, how much, and how variable"

### Supplemental Movie S1

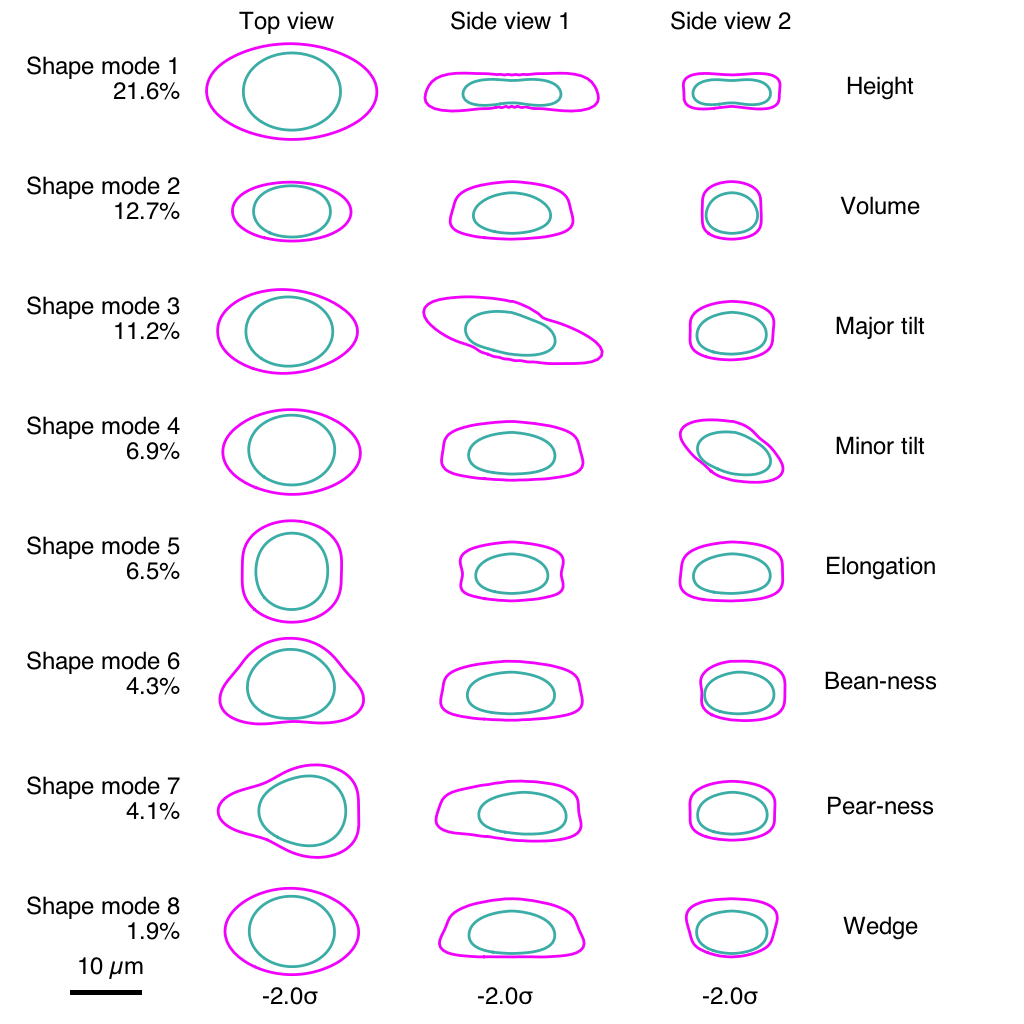

### Supplemental Movie S3

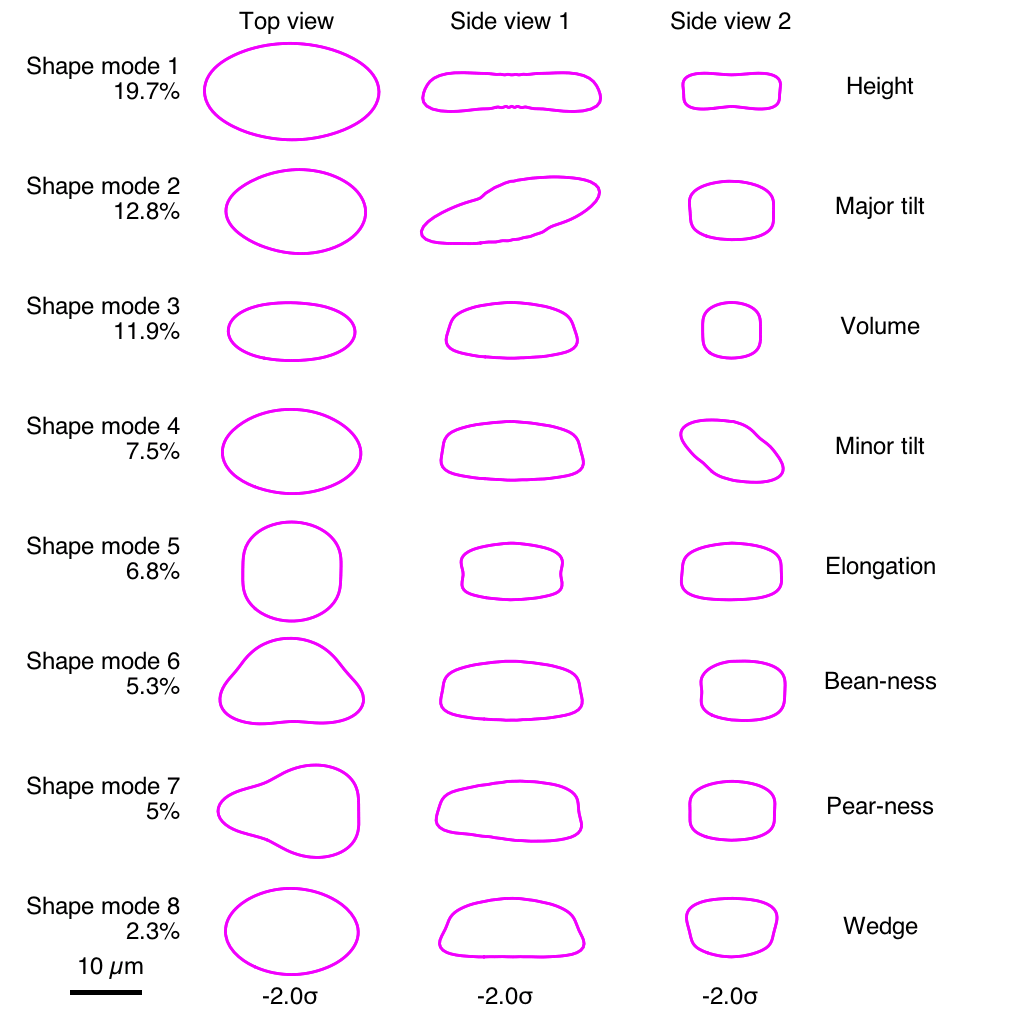

### Supplemental Movie S4

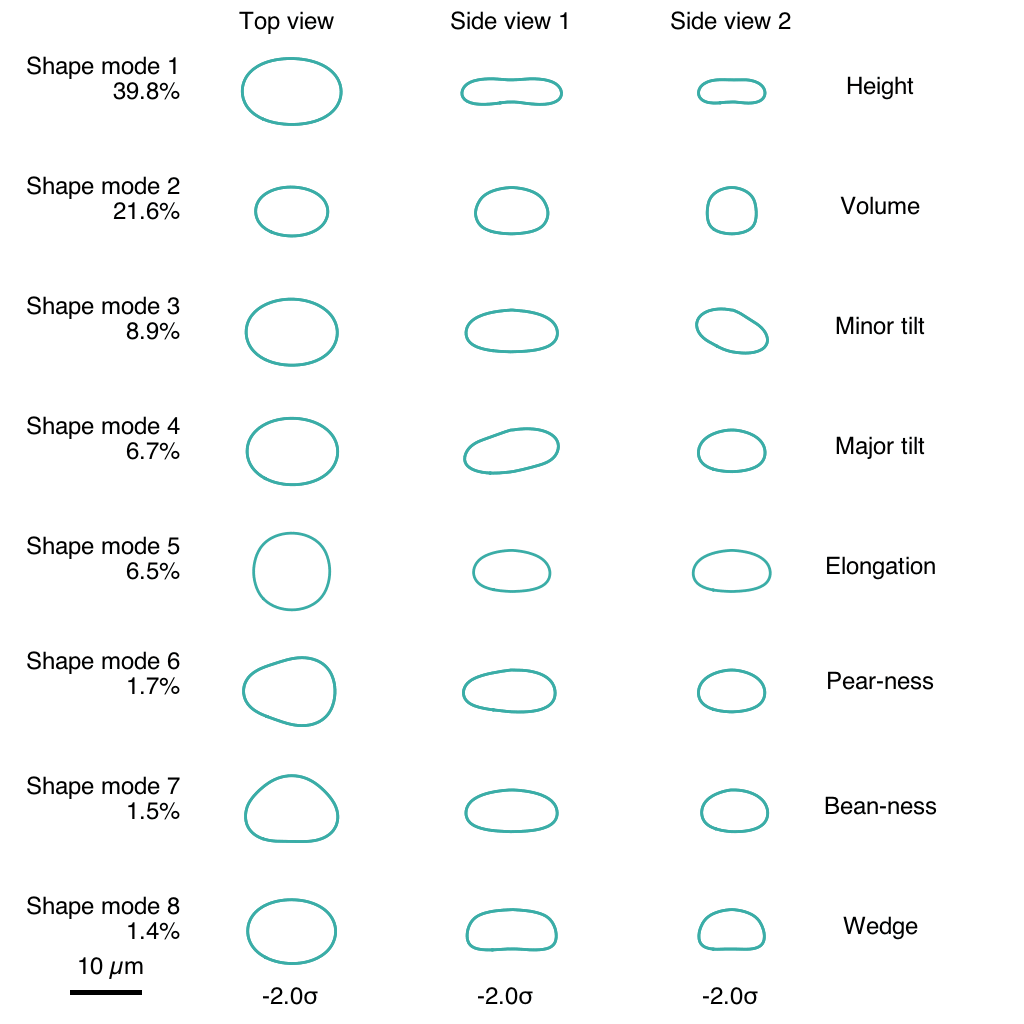
